# Lipid Bilayer Induces Contraction of the Denatured State Ensemble of a Helical-Bundle Membrane Protein

**DOI:** 10.1101/2021.05.17.444377

**Authors:** Kristen A. Gaffney, Ruiqiong Guo, Michael D. Bridges, Daoyang Chen, Shaima Muhammednazaar, Miyeon Kim, Zhongyu Yang, Anthony L. Schilmiller, Nabil F. Faruk, Xiangda Peng, A. Daniel Jones, Liangliang Sun, Wayne L. Hubbell, Tobin R. Sosnick, Heedeok Hong

## Abstract

Defining the denatured state ensemble (DSE) and intrinsically disordered proteins is essential to understanding protein folding, chaperone action, degradation, translocation and cell signaling. While a majority of studies have focused on water-soluble proteins, the DSE of membrane proteins is much less characterized. Here, we reconstituted the DSE of a helical-bundle membrane protein GlpG of *Escherichia coli* in native lipid bilayers and measured the DSE’s conformation and compactness. The DSE was obtained using steric trapping, which couples spontaneous denaturation of a doubly biotinylated GlpG to binding of two bulky monovalent streptavidin molecules. Using limited proteolysis and mass spectrometry, we mapped the flexible regions in the DSE. Using our paramagnetic biotin derivative and double electron-electron resonance spectroscopy, we determined the dimensions of the DSE. Finally, we employed our *Upside* model for molecular dynamics simulations to generate the DSE including the collapsed and fully expanded states in a bilayer. We find that the DSE is highly dynamic involving the topology changes of transmembrane segments and their unfolding. The DSE is expanded relative to the native state, but only to 55–90% of the fully expanded condition. The degree of expansion depends on the chemical potential with regards to local packing and the lipid composition. Our result suggests that the *E. coli*’s native lipid bilayer promotes the association of helices in the DSE of membrane proteins and, probably in general, facilitating interhelical interactions. This tendency may be the outcome of a general lipophobic effect of proteins within the cell membranes.

**Significance:** Here, we delineate the conformation of the denatured state ensemble (DSE) of a membrane protein confined in a native lipid bilayer and assay whether the bilayer permits full expansion or nonspecific collapse of the DSE. Using the intramembrane protease GlpG as a model, we find that the denatured state is a dynamic ensemble involving topological changes and local unfolding of transmembrane segments. The bilayer tends to contract the DSE relative to the fully lipid-solvated, expanded conformations while the degree of compactness is determined by the balance between protein-lipid, lipid-lipid and protein-protein interactions. These findings provide new insights into the lipid bilayer as a solvent that mediates folding, chaperone action, turnover and protein-protein interactions in cell membranes.

---

Denatured states and intrinsically disordered proteins are involved in a variety of cellular events serving as targets for degradation, chaperone action, translocation and cell signaling (1–5). They also influence thermodynamic stability and direct early folding events. Thus, delineating the denatured state ensemble (DSE) has been a subject of extensive study (1, 6–8). For water-soluble proteins, DSE can be described as a collection of disordered conformations that interconverts on a timescale much faster than folding (9–11).

For the quantitative description of the DSE, polymer theory has proven extremely useful (12–14): For a given polymer/solvent combination, the solvent quality can be classified into three limiting regimes depending on the relative strengths between intrachain and chain–solvent interactions (13). In a “good” solvent, chain–solvent interactions are much more favorable than the intrachain interactions and the DSE is well described as a self-avoiding random walk (SARW). In a “*θ*” solvent, the intrachain and chain–solvent interactions are balanced such that the intrachain interaction cancels out the expansion caused by excluded chain volume and the protein behaves like a RW. In a “poor” solvent, the intrachain interaction exceeds the chain–solvent interactions, inducing a contraction or even collapse of the chain into a globule.

For these three scenarios, solvent quality can be quantified by the Flory exponent, *ν*, in the relationship between the radius of gyration (*R*_G_) and the number of monomeric units (*N*): *R*_G_ ∝ *N^ν^*. The exponent is *ν* = 3/5, 1/2 and 1/3 for a good, *θ* and poor solvent, respectively. The *ν* value for SARW is observed for unfolded proteins in elevated denaturant concentrations with small angle X-ray scattering (SAXS) (15–21) and Förster resonance transfer (FRET) (21, 22). However, for a variety of intrinsically disordered proteins (IDPs) with protein-like sequences and the DSE of soluble proteins, *ν* values are typically above 1/2 under native conditions, much higher than anticipated given the general perception that soluble proteins undergo a hydrophobic collapse prior to folding (17-19, 21-23).

In contrast, the properties of the DSE are not as well understood for membrane proteins. The concept of solvent quality is harder to quantitatively apply due to the physical constraints of the quasi-two-dimensional (2D) hydrophobic lipid bilayer. Generally, membrane proteins are not well represented by simple polymers and the folding of helical membrane proteins occurs through at least two distinct stages (24). In Stage I, hydrophobic segments in a polypeptide chain insert into the lipid bilayer as transmembrane (TM) helices. These TM helices can diffuse around within the bilayer although the extent is limited by the length of the interconnecting loops and the requirement that they remain in the bilayer. In cells, insertion is co-translationally mediated by a translocon complex (25, 26). In Stage II, TM helices associate into a compact native structure.

The underlying principle of the two-stage model is that the folding and transfer of helices into a bilayer and their subsequent association are driven by distinct driving forces. In Stage I, the formation and insertion of TM helices are typically favored by the hydrophobic nature of TM segments and a high desolvation cost of unpaired backbone hydrogen bonding within the bilayer (25, 27, 28). The role of water is much less important in Stage II as the individual TM helices have already been dehydrated and are favorably solvated with lipids. The association of TM helices typically requires good van der Waals packing and polar interactions (29–32). A combination of molecular forces determines how favorable TM helix association is and whether the lipid bilayer functions as a good solvent for DSEs of membrane proteins in general.

So far, the DSEs of helical membrane proteins have been characterized under denaturing conditions induced by urea or guanidine hydrochloride (GdnHCl) (33–35), anionic detergents including sodium dodecyl sulfate (SDS) (30, 36–39) or mechanical force (40, 41) in various hydrophobic environments (*e.g.*, micelles, lipid–detergent mixtures, or lipid bilayers). These studies find that relative to native states, DSEs are expanded but their TM helices are constrained by unfolded interhelical loops. The degree of expansion also depends on the choice of denaturant and may be limited by the size of micelles (33, 37–40, 42). Furthermore, lipid bilayers provide a distinct but biologically relevant 2D environment. As a result, the conformational features of the DSE obtained under denaturing conditions may be different from those in cellular membranes.

Here, we reconstitute the DSE of a helical membrane protein GlpG of *E. coli* in native lipid bilayers in the absence of denaturant and study its conformation and compactness. GlpG is a member of the rhomboid intramembrane protease family possessing moderate thermodynamic stability and high unfolding activation energy (***SI Appendix* Table S1**) (40–44). As with any stable protein, the characterization of the DSE under native conditions is challenging due to its low population and short lifetime (45, 46). We generate GlpG’s DSE using our steric trapping method, which couples the spontaneous denaturation of a doubly biotinylated protein to the binding of two bulky monovalent streptavidin (mSA) molecules (**Fig. 1a**). By combining multiple experimental measurements and MD simulations to probe the flexibility and physical dimensions of the DSE at the local and global levels, we demonstrate that the DSE is highly dynamic involving extensive helix unfolding and changes in membrane topology. The DSE expands relative to the native state but retains a degree of compactness in the lipid bilayers. The degree of expansion depends on the local stability in the native state and the lipid composition. We suggest that the tendency of the DSE to partially contract is attributed to the lipophobic effect of TM segments.

**Fig. 1:**
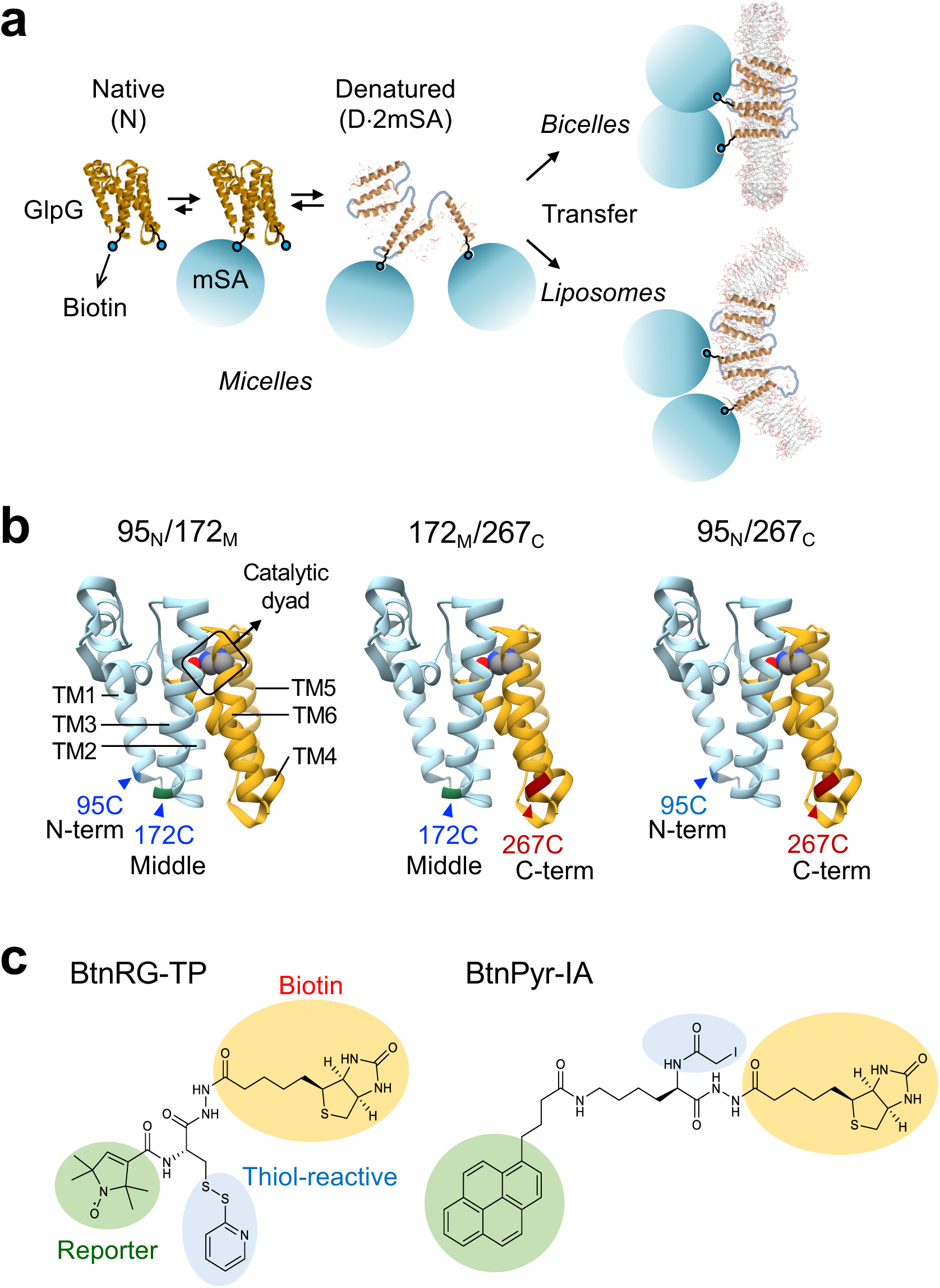
Steric trapping strategy to reconstitute denatured GlpG in the lipid bilayers. **(a)** Doubly biotinylated GlpG was first denatured using steric trapping by the addition of excess mSA in DDM micelles. Sterically denatured GlpG doubly bound with mSA was transferred to preformed bicelles (DMPC:DMPG:CHAPS, molar ratio = 3:1:1.4) or liposomes composed of *E. coli* phospholipids. **(b)** Double cysteine variants employed for the steric trapping of the denatured states of GlpG. In each variant, designated cysteine residues were labeled with a thiol-reactive biotin derivative with a spectroscopic reporter group as shown in **(c)**. The regions of the backbone colored in cyan and orange indicate the N- and C-subdomain, respectively. **(c)** Thiol-reactive biotin derivatives with a paramagnetic spin label (*left*, BtnRG-TP) and fluorescent pyrene (*right*, BtnPyr-IA) used in this study (42). TP: thiopyridine; IA: iodoacetamide

## RESULTS

### Reconstitution of the on-pathway DSE of GlpG in lipid bilayers

To measure the properties of the DSE, we applied our steric trapping method (**Fig. 1a**) (42). GlpG is labeled with biotin tags at two specific residues that are spatially close in the folded state but on different structural segments. The first mSA can bind unhindered to either biotin tag. The binding of a second mSA, however, is sterically disallowed if the two biotins are close in the native state but is allowed when the protein is denatured. This steric trapping strategy takes advantage of the stability and long life-time of the mSA–biotin complex (*K*_d,biotin_ = ∼10^–14^ M; *k*_off,biotin_ = ∼days) (47), which enables the study of the DSE even under native conditions.

Previously, we have shown that the TM domain of GlpG (residues 87–276) can be separated into the N- (TM1–L1–TM2–L2–TM3–L3) and C-subdomains (TM4–L4–TM5–L5– TM6), each having their own distinct folding behaviors (**Fig. 1b**) (42). The three sites selected for biotinylation were located at the **N**- and **C**-termini, and on the middle loop (**M**) connecting the helices TM2 and TM3 (Pro95_N_, Val267_C_, and Gly172_M_, respectively) (42). These three sites were used two at a time to disrupt either the entire protein (with the residue pair 95_N_267_C_) or the N- or C-subdomain (with 95_N_172_M_ or 172_M_267_C_). Each of the three residue pairs was substituted with a pair of cysteine residues that were labeled with the thiol-reactive biotin derivative possessing a spectroscopic probe (*e.g.*, fluorescent pyrene, BtnPyr, or a nitroxide spin label, BtnRG) (**Fig. 1c** and ***SI Appendix* Fig. S1**) (42). Conjugation of the biotin derivative to each single cysteine variant and binding of a mSA molecule to each biotin label do not inhibit the proteolytic activity of GlpG relative to wild type (WT) for the model substrate LYTM2 (the second TM segment of a *E. coli* lactose permease, LacY) (***SI Appendix* Fig. S2a**) (42). We reconstituted sterically denatured GlpG into two lipid bilayer environments: *1)* Large negatively charged bicelles, which are discoidal bilayer fragments edge-stabilized by detergent, composed of *di*C_14:0_-phosphatidylcholine (PC), *di*C_14:0_-phosphatidylglycerol (PG), and 3-[(3-cholamidopropyl) dimethylammonio]-1- propanesulfonate hydrate (CHAPS) (molar ratio = 3:1:1.4, disk diameter = ∼30 nm) (48). *2)* Large unilamellar liposomes composed of *E. coli* phospholipids (diameter = 120–170 nm, ***SI Appendix* Fig. S3**) to provide the native lipid environment for *E. coli* GlpG.

We initially attempted to obtain the DSE using steric trapping with native GlpG reconstituted in bicelles and liposomes. However, incubation with excess mSA did not yield noticeable denaturation for a week when GlpG activity was used as a folding indicator, implying that native GlpG has high thermodynamic or kinetic stability in bilayers (***SI Appendix* Fig. S4**). However, denaturation was achieved for all three constructs within 48 h of incubation with mSA in dodecylmaltoside (DDM) detergent micelles (***SI Appendix* Figs. S2b** and **S4**). With doubly conjugated BtnRG, the activities of the 95_N_172_M_, 172_M_267_C_, and 95_N_267_C_ constructs in micelles were reduced to 43 ± 15%, 26 ± 9%, and 12 ± 7% of their native enzymatic activity, respectively (***SI Appendix* Figs. S2c**). The residual activity indicates incomplete denaturation. An examination of the biotinylation levels indicates that this activity likely stemmed from incomplete biotin labeling (***SI Appendix* Figs. S1 and S2c**).

For reconstitution in bilayers, we first prepared sterically denatured GlpG in DDM micelles (**Fig. 1a**). Then, denatured GlpG was transferred to bicelles or to liposomes by direct injection (for the liposomes, detergents were further removed using polystyrene beads, ***SI Appendix* Fig. S3**). To verify incorporation of sterically denatured GlpG into bicelles, we employed a fluorescence-quenching assay (**Fig. 2a**). Here, the double cysteine variants of GlpG were labeled with fluorescent BtnPyr (**Fig. 1c**). Native and sterically denatured GlpG were injected into the bicelles containing the quencher-labeled lipid, DOPE (*di*C_18:1c9_-phosphatidylethanolamine)-dabcyl. After injection, pyrene fluorescence from both the native and denatured samples of GlpG was quenched to a level close to that observed for full incorporation.

**Fig. 2:**
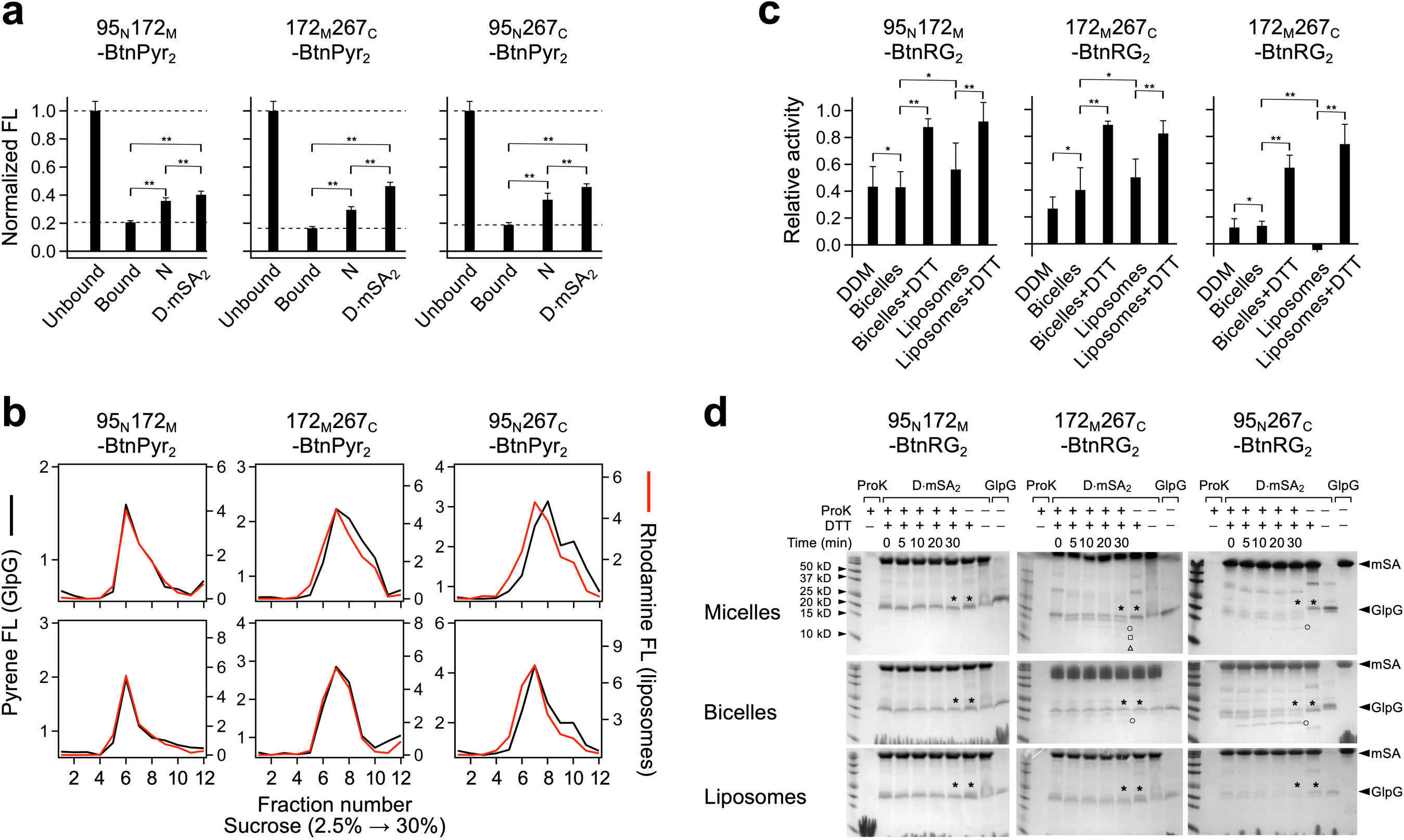
Reconstitution of denatured GlpG in the lipid bilayers. (a) Fluorescence quenching assay to measure incorporation of native (N) and sterically denatured (D*′*mSA_2_) into bicelles. GlpG was doubly labeled with BtnPyr and bicelles contained quencher-labeled lipids, DOPE-dabcyl. Negative control (unbound GlpG): pyrene-labeled mSA, which was highly soluble in water. Positive control (bound): native GlpG doubly labeled with BtnPyr, which was first reconstituted in DMPC:DMPG:DOPE-dabcyl liposomes and then solubilized by CHAPS to form bicelles. Error bars denote ± SEM (*n* = 3). Based on Student’s *t*-test, significant and insignificant differences between a data pair are marked with single (*P* >0.05) and double asterisks (*P* <0.05). **(b)** Liposome flotation assay in a sucrose gradient to measure membrane-association of native and denatured GlpG. GlpG was doubly labeled with BtnPyr and liposomes contained DPPE-rhodamine. **(c)** Proteolytic activity of GlpG as a measure of the extent of denaturation and refolding. GlpG doubly labeled with BtnRG was first denatured upon addition of excess mSA in micelles (“Micelles”). The denatured state was maintained upon reconstitution in bicelles (“Bicelles”) or liposomes (“Liposomes”). The addition of DTT, which cleaved the disulfide linkage between GlpG and the biotin label bound with mSA, allowed for refolding in both bicelles (“Bicelles + DTT”) and liposomes (“Liposomes + DTT”). The activity was normalized relative to that of native GlpG without mSA in each hydrophobic phase. Error bars denote ± SEM (*n* = 3). Based on Student’s *t*-test, significant and insignificant differences between a data pair are marked with single (*P* >0.05) and double asterisks (*P* <0.05). **(d)** Limited proteolysis of native (N) and sterically denatured GlpG (D*×*mSA_2_) by Proteinase K (ProK) in (*top*) DDM micelles, (*middle*) bicelles, and (*bottom*) liposomes. After the termination of proteolysis at each time point, DTT was added to release bound mSA from GlpG. A comparison of the intensities of GlpG bands in the presence and absence of ProK confirmed the proteolysis of GlpG (asterisk marks). The proteolytic peptide fragment larger than 10 kDa is marked with a symbol (open circles, 17 kDa; open squares, 13 kDa; open triangles, 11 kDa) on the right side of each band.

To verify the incorporation of denatured GlpG into liposomes, we employed a liposome-flotation assay (**Fig. 2b** and ***SI Appendix* Fig. S5a**). Native and denatured GlpG were doubly labeled with BtnPyr and reconstituted in the liposomes containing the fluorophore-labeled lipid, DPPE (*di*C_16:0_-PE)-rhodamine). After centrifugation in a sucrose gradient, both native and denatured GlpG co-floated with the liposomes, indicating membrane association. Native and denatured GlpG reconstituted in liposomes were resistant to sodium carbonate extraction, indicating membrane integration (***SI Appendix* Fig. S5b**).

Next, we tested whether the denaturation status of GlpG initially prepared in micelles was retained after reconstitution in bicelles and liposomes (**Fig. 2c** and ***SI Appendix* Fig. S6)**. To trap the denatured state, we used BtnRG to doubly biotinylate GlpG. This biotin label forms a disulfide linkage to cysteine, which can be broken with the addition of a reducing agent, dithiothreitol (DTT). After reconstitution of the GlpG samples containing mSA, the activities relative to the native forms of the double biotin variants 95_N_172_M_, 172_M_267_C_, and 95_N_267_C_ were 43 ± 12%, 40 ± 17% and 12 ± 7% in bicelles, and 56 ± 20%, 49 ± 14% and 0 ± 4% in liposomes, respectively, similar to the values in micelles. Thus, sterically denatured GlpG in micelles largely remained denatured after transfer to bilayer environments.

We tested whether denatured GlpG in the lipid bilayers can refold after the release of the steric repulsion between bound mSA molecules (**Fig. 2c**). Upon addition of DTT to dissociate the BtnRG labels bound with mSA from GlpG, the relative activity increased to >70%. Thus, sterically denatured GlpG in the bilayers can refold to its native structure and is not a dead-end product incapable of refolding.

### Limited proteolysis reveals the flexibility of the DSE under native conditions

To investigate the conformational features of GlpG’s DSE in the native lipid bilayer and other hydrophobic environments, we first employed limited proteolysis using Proteinase K (ProK). ProK is a robust endopeptidase, which nonspecifically proteolyzes water-exposed unstructured regions rather than structured or membrane-buried regions (49, 50).

Upon denaturation, the conformational flexibility and solvent accessibility increased throughout the protein. Time-dependent proteolysis was measured for native and sterically denatured GlpG trapped for the three biotin combinations (**Fig. 2d**). In contrast to native GlpG (23 kDa, the TM domain), which was resistant to proteolysis except at the termini (***SI Appendix* Fig. S7**), denatured GlpG was rapidly proteolyzed within 10 min in all hydrophobic environments. However, 20–50% of the GlpG sample containing mSA was not proteolyzed probably due to incomplete biotin labeling (***SI Appendix* Figs. S1** and **S2c**): GlpG doubly bound with mSA was preferentially proteolyzed while unbound or singly bound GlpG was protease resistant (***SI Appendix* Figs. S7–S8**). Notably, whereas denatured GlpG trapped with the biotin pair at the N-subdomain (95_N_172_M_) was extensively proteolyzed into small fragments (<8 kDa), the denatured protein trapped at either the C-subdomain (172_M_267_C_) or the N- and C-termini (95_N_267_C_), yielded larger fragments in micelles and bicelles (∼11, ∼13, or ∼17 kDa). This result supports our previous finding that in micelles, the disruption of the N-subdomain leads to global denaturation whereas the C-subdomain undergoes subglobal denaturation with the balance of the protein remaining intact (42). In liposomes, however, denatured GlpG was extensively proteolyzed regardless of the position of the biotin pair, implying that the entire chain is less structured, contracted or buried in liposomes than in micelles and bicelles.

To identify the unstructured and solvent-accessible regions that are subject to proteolysis, we employed capillary-zone electrophoresis (CZE)-tandem mass spectrometry (MS/MS) or reverse-phase liquid chromatography (RPLC)-MS/MS (***SI Appendix* Tables S2–S7**). Each method utilizes a distinct principle for peptide separation (electrophoretic mobility for CZE and solubility for RPLC), allowing for complementary peptide identification (51, 52). We chose the double-biotin variant, 172_M_267_C_–BtnRG_2_, because it had a higher biotin-labeling efficiency than the other variants and yielded a larger fraction of denatured GlpG molecules. Overall, the RPLC-MS/MS approach enabled the detection of the larger peptides from the TM regions in the native structure, while the smaller peptides from the loops were obtained from CZE-MS/MS.

The native state ensemble (NSE) of GlpG in the bilayer likely undergoes functional motions that permit cleavage by ProK. We identified several peptide fragments from native GlpG in the liposomes (**Fig. 3a** *top*). These sequences were mapped onto the kinked cytoplasmic end of TM2 and the middle of TM4 and TM6. Notably, these regions are directly involved in the proteolytic mechanism of GlpG: The interface between TM2 and the gating helix TM5 forms the substrate binding site (53). TM4 and TM6 are packed through the conserved glycine-zipper motif harboring the catalytic dyad Ser201–His254 (54). Upon denaturation, the entire length of TM4 and TM6 became susceptible to proteolysis in addition to the periplasmic half of TM3 and the flanking loops of TM5. The middle regions of TM3, TM4 and TM6 were highly susceptible to proteolysis, and these three helices are the least hydrophobic in GlpG (**Fig. 3b**), suggesting that they can transiently become unstructured and solvent exposed in the DSE.

**Fig. 3:**
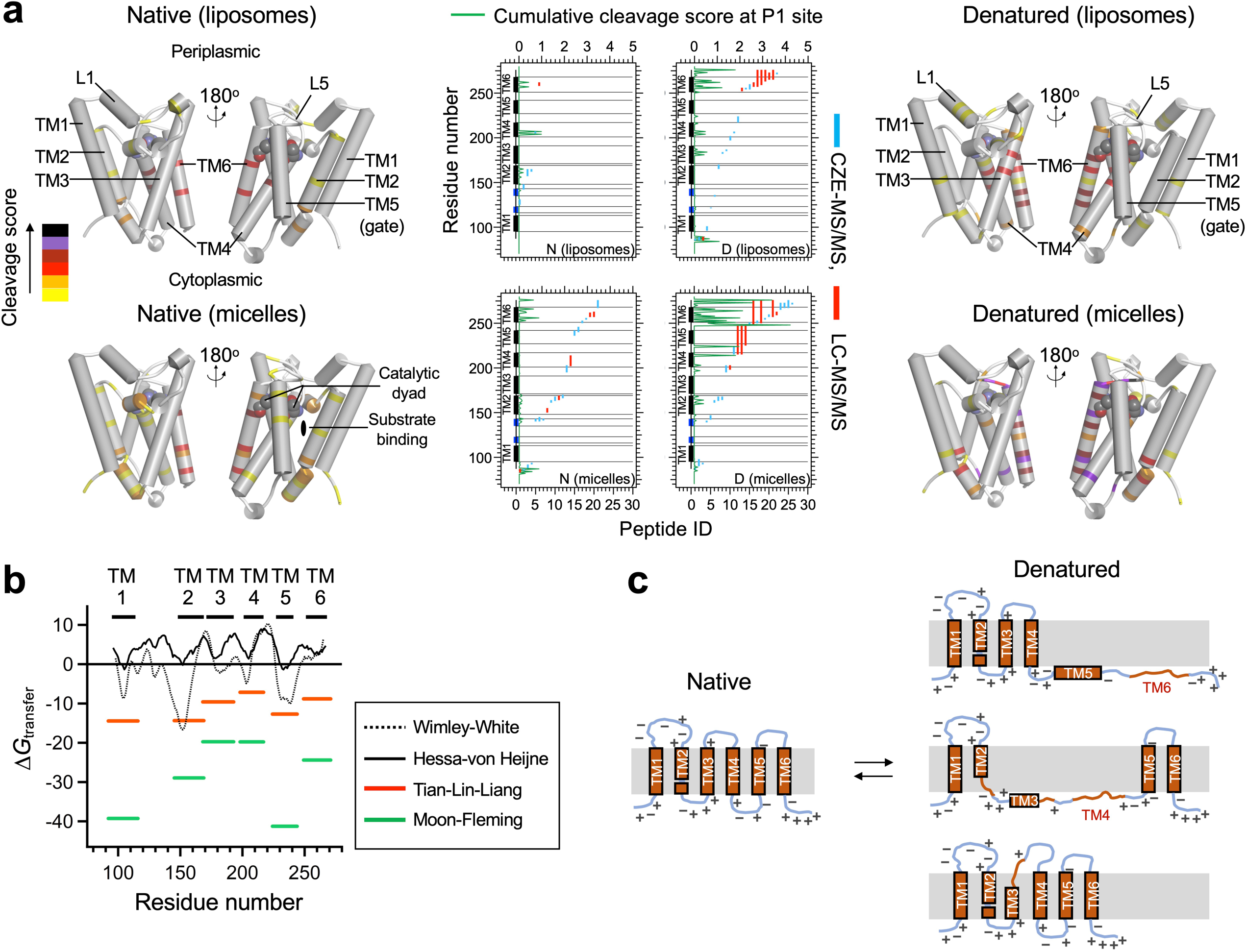
Mapping the flexible structural regions in native and denatured GlpG using limited proteolysis and mass spectrometry. **(a)** Native (*left*) and denatured (*right*) GlpG (the double biotin variant 172_M_267_C_) reconstituted in liposomes (*top*) and micelles (*bottom*) were proteolyzed using ProK. The proteolysis products were analyzed using CZE- and LC-MS/MS and mapped on to the primary and secondary structures (*center*). The scores reflecting the abundance and reliability of the analyzed peptides (104) were summed at each P1 cleavage site (*i.e.*, the substrate residue N-terminal to the cleaved peptide bond) by ProK and mapped on each plot (green lines) and structure (the heat map in the far *left*). The catalytic dyad (Ser201 and His254) is shown in spheres in each structure. **(b)** Hydrophobicity analysis of the TM domain of GlpG using the Wimley-White (WW) water-octanol (106), Hessa-von Heijne (HvH) translocon-membrane (25), Tian-Lin-Liang (TLL) depth-dependent (62, 107), and Moon-Fleming (MF) water-membrane (108) hydrophobicity scales. For the WW and HvH scales, a 19-residue window was slid along the sequence of GlpG summing *ΔG*_transfer_ of each residue. For the TLL and MF scales, the *ΔG*_transfer_ values only for the residues in each TM helix from the structure were summed. **(c)** Possible modes of the membrane-topology dynamics and unfolding in the DSE of GlpG in *E. coli* liposomes. The modes were deduced on the basis of the limited proteolysis, MS/MS, MD simulations and the charge distributions in the membrane-water interfacial regions of GlpG.

In micelles, the overall proteolysis pattern was similar to that in the bilayers for native and denatured GlpG, although in micelles, cleavage was more pronounced in the C-subdomain (**Fig. 3a** *bottom*). The extensive proteolysis in the C-subdomain in both bilayers and micelles points to higher dynamics at this end of the protein, being an intrinsic property of the protein rather than stemming from steric repulsion in this subdomain. Supporting this reasoning, denatured GlpG trapped using the two biotinylation sites (95_N_172_M_) in the N-subdomain still underwent extensive proteolysis in the C-subdomain (***SI Appendix* Fig. S9**).

In summary, our proteolysis study reveals dynamic features of the DSE under native conditions. The DSE involves various modes of membrane-topology changes including the unfolding of TM segments due to their low hydrophobicity and, probably, the charge distribution in the flanking regions (**Fig. 3c**). Although we minimized the duration of proteolysis (5–7 min) to capture unstructured regions in the DSE of full-length GlpG, it is likely that a certain fraction of the identified peptides were obtained from the polypeptides already cleaved by proteolysis (“double hit”). Even so, the proteolysis patterns nonetheless would reflect intrinsic conformational features of individual structural segments in their isolated forms.

### Contraction of the DSE measured with DEER

We next examined the compactness of the DSE in micelles, bicelles and liposomes using DEER spectroscopy, which is suitable for measuring distances between 15‒70 Å. Specifically, we measured the interspin distances between a pair of paramagnetic biotin labels (BtnRG) in native and sterically denatured GlpG for the three constructs 95_N_172_M_–BtnRG_2_, 172_M_267_C_–BtnRG_2_, and 95_N_267_C_–BtnRG_2_ (**Fig. 4**). The probability distribution of distances between the spin labels was obtained from the time-dependent dipolar evolution data using Tikhonov regularization or model-based (*e.g.*, Gaussian distribution) fitting algorithm (55). To minimize unwanted intermolecular dipolar coupling between multiple spin-labeled GlpG molecules in a single liposome, we increased the lipid-to-protein molar ratio by six-fold (from 2,000 to 12,000) compared to our activity and proteolysis measurements and incorporated unlabeled GlpG (an inactive variant, S201A) at a three- or six-times molar excess relative to spin-labeled GlpG (***SI Appendix* Figs. S10–S11**). For the three denatured variants, fitting of the time-dependent dipolar evolution data using a non-negative Tikhonov regularization model yielded uncharacteristic, highly heterogeneous distance distributions with multiple local maxima (***SI Appendix* Figs. S11–S12**). Thus, considering the dynamic and heterogeneous nature of the DSE, we chose to fit the data to a single Gaussian distribution model (**Fig. 4**; ***SI Appendix* Fig. S12** and **Tables S8–S9** for comparison of the two fitting methods).

**Fig. 4:**
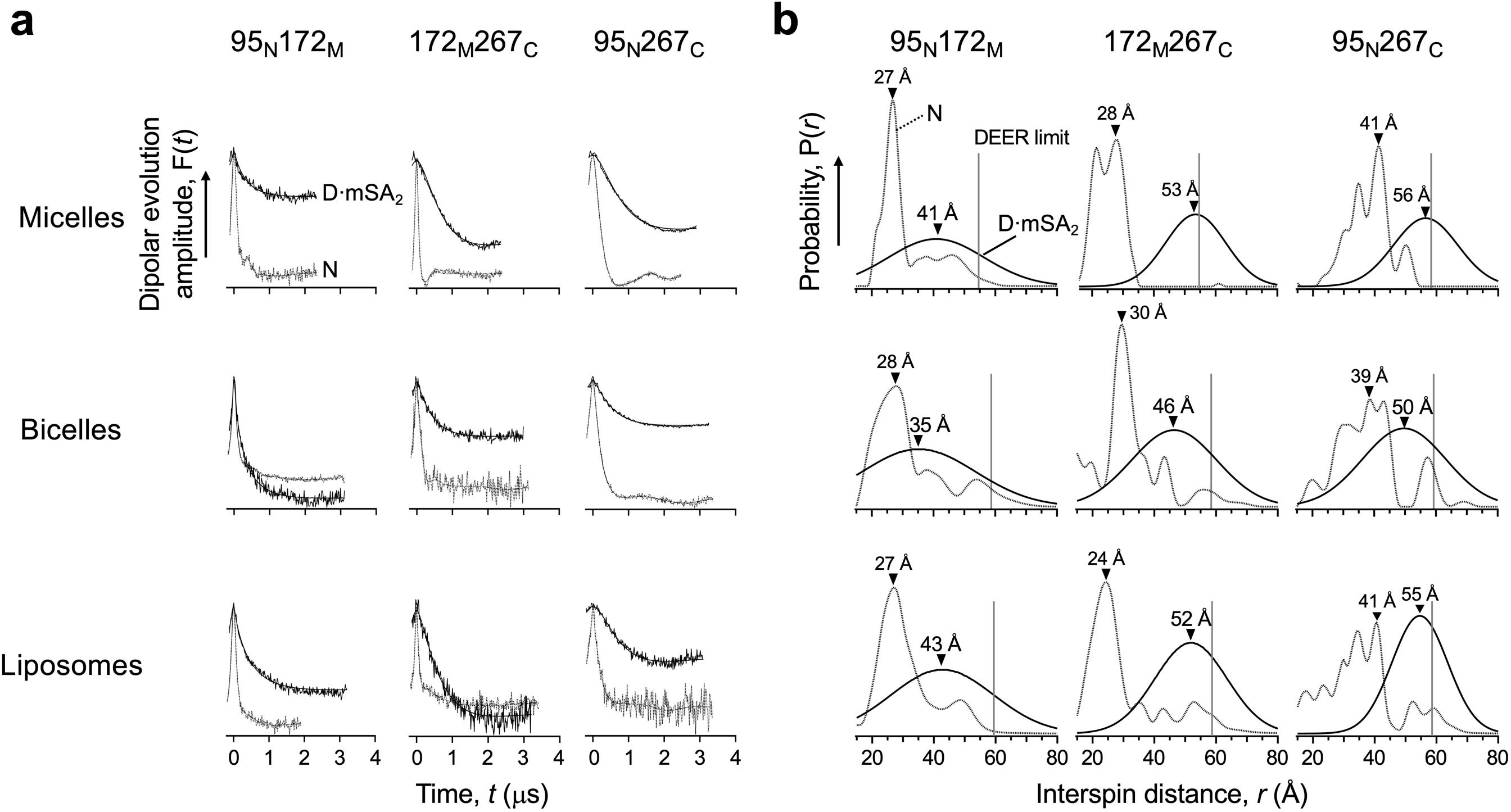
The physical dimension of the denatured states of GlpG measured by DEER. **(a)** Time-dependent dipolar evolution data for native (N) and sterically denatured GlpG (D·mSA_2_). The data were fitted via non-negative Tikhonov regularization for native GlpG and a single Gaussian model for sterically denatured GlpG to obtain the interspin distances as in **(b)**. **(b)** Distance distributions between the spin labels at the designated residues in native and sterically denatured GlpG. In each distribution, the most probable distances (*r*_prob_) are shown. N (dotted lines): the distributions obtained with a non-negative Tikhonov regularization algorithm; D·mSA_2_ (solid lines): the distributions obtained from fitting to a single Gaussian model.

We have previously shown that upon denaturation by steric trapping in DDM micelles, the most probable interspin distances (*r*_Prob_) increase from 27 to 41 Å (1.5-fold) for 95_N_172_M_ and from 28 to 53 Å (1.9-fold) for 172_M_267_C_, respectively (42). For the new 95_N_267_C_ variant, we found that the N-to-C distance increased from *r*_Prob_ = 41 to 56 Å (1.4-fold) upon denaturation (**Fig. 4**). Thus, steric trapping overall induced an expansion of the DSE under non-denaturing micellar conditions.

In the native state, *r*_Prob_’s in bicelles and liposomes were overall similar to those in micelles (**Fig. 4b**). We expected that the quasi-2D lipid bilayer would confine the movements of the TM segments in the DSE, resulting in a more compact DSE relative to micelles, which are less organized assemblies. Indeed, the DSEs in bicelles expanded less than in micelles (1.3-fold for 95_N_172_M_, 1.5-fold for 172_M_267_C_, and 1.3-fold for 95_N_267_C_ relative to the native state). Interestingly, the DSE in liposomes expanded more than in bicelles despite the same quasi-2D bilayer constraint (1.6-fold for 95_N_172_M_, 2.2-fold for 172_M_267_C_, and 1.3-fold for 95_N_267_C_ relative to the native state). Overall, the expansion of the DSE depended on the position of the biotin pair at which the steric repulsion was exerted as well as the lipid composition (*e.g.,* DMPC:DMPG:CHAPS *vs E. coli* phospholipids).

To test the effect of doubly bound mSA molecules on the compactness of the DSE, we measured the interspin distances in the SDS-induced DSE of a double biotin variant with and without mSA (***SI Appendix* Fig. S13**). In SDS, mSA is resistant to denaturation and can still bind to the biotin labels (***SI Appendix* Fig. S1**). Upon mSA binding in SDS, the *r*_prob_ increased by ∼3 Å, implying that doubly bound mSA molecules may induce a mild additional expansion of the DSE. SDS molecules are expected to bind the surface of mSA (56). The resulting negatively charged mSA–SDS complexes may repel each other when doubly bound to GlpG, inducing further expansion of the DSE. Nonetheless, the *r*_prob_ measured with bound mSA in a high concentration of SDS (∼4% w/v) was similar to that in neutral DDM micelles (***SI Appendix* Fig. S13**). Thus, repulsive or attractive interactions between bound mSA molecules, even if they exist, do not noticeably affect the compactness of sterically denatured GlpG. This result supports our modeling of mSA-bound as hard spheres in the DSE simulations as described in the next section.

### DSE simulations

To generate plausible models of compact and expanded DSEs and provide a reference for comparison to the DEER distance distributions, we used our *Upside* MD simulation algorithm suitably modified for mSA-bound membrane proteins. This recently created algorithm can fold small proteins with accuracy comparable to all-atom methods (57, 58). *Upside* employs six atoms per residue: three backbone atoms (N, C*_α_* and C), the carbonyl oxygen and amide proton along with a side chain represented by a directional-dependent bead. After every MD step, the side chains are globally repacked to have the lowest global free energy. Our mild coarse-graining and the use of an implicit solvent along with the global side-chain packing step explains much of the 10^3^–10^4^-fold speed up compared to standard all-atom simulations (57, 58).

Our membrane-burial potential includes knowledge-based, depth-dependent energies for side-chain burial and backbone hydrogen bonding within the bilayer. Helices are allowed to unfold and refold in the bilayer during the simulations (59, 60). At every MD step, side-chain burial is recalculated to account for the exchange of protein-lipid interactions for protein-protein interactions as helices come into contact with each other. This recalculation avoids overestimating (double counting) the energetics of helix association within the membrane. Through the careful treatment of the bilayer boundary position and the exposure of side chain and backbone within the bilayer, our potential can accurately predict the optimal bilayer thickness and protein location, as compared to the OPM (Orientations of Proteins in Membranes) method (61). Our force-induced unfolding studies using the membrane potential for GlpG and bacteriorhodopsin are in good agreement with experiment (59).

Twenty 4 msec simulations were run for native GlpG and for the three doubly mSA-bound versions (95_N_172_M_, 172_M_267_C_ and 95_N_267_C_) at five temperatures, *T* = 274 K, 308 K, 343 K, 377 K, and 411 K (400 simulations total, ***SI Appendix* Figs. S14a***−***f**). To attach the mSA molecules, each of the two biotinylation sites on GlpG was linked with a stiff 4 Å spring to Asn49 on each mSA, a position near the biotin-binding pocket (**Fig. 5a**). For modeling purposes, each mSA was held in its native conformation using a set of stiff restraint springs. To eliminate steric overlap, only short-range repulsive interactions were allowed between the two bound mSA molecules with each other and with GlpG (a sigmoid potential with a ∼1 Å width and drop-off at 8 Å, and the repulsive energy of 2*RT* in which *R* is a gas constant). To inhibit the mSA molecules from entering the bilayer, they were subject to the same membrane burial potential as GlpG.

**Fig. 5:**
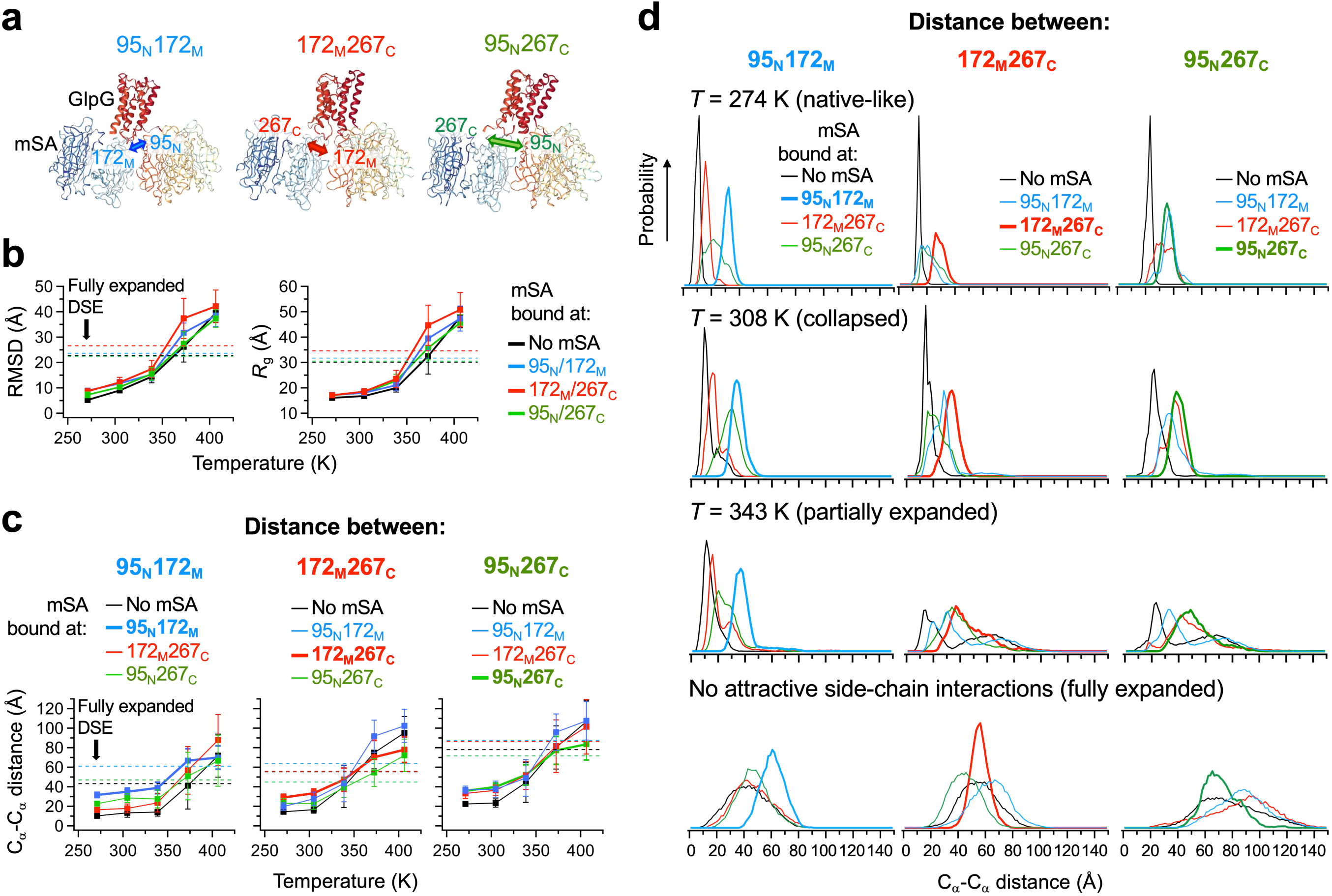
*Upside* MD simulation under the statistical membrane-burial potential. **(a)** Modeling of GlpG with doubly bound mSA molecules. Two mSA molecules were attached to the designated residue pair on GlpG through a stiff 4 Å virtual spring (spring constant = 30 kcal·mol^-1^·Å^-1^). **(b)** The C*_α_*-RMSD and radius of gyration (*R*_g_) of GlpG from temperature-dependent simulation. Each value corresponds to the average from 20 independent simulations at each temperature. Each error bar denotes ± SEM (*n* = 20). **(c)** The C*_α_*–C*_α_* distances between each designated residue pair monitored at an increasing temperature. When the distances were monitored between a specific residue pair (“Distance between”), mSA molecules are bound to the same residue pair, or the other residue pairs where the distances were not monitored (“mSA bound at”). **(d)** Distribution of the C*_α_*–C*_α_* distances between the designated residue pair in GlpG with or without bound mSA. The simulations were carried out with all energy terms (“native-like”, “collapsed” and “partially expanded”) or missing attractive side-chain interactions (“Fully expanded”).

During the simulations, GlpG and the three doubly mSA-bound versions maintained a native and near-native fold at the lowest two temperatures, 274 K and 308 K (***SI Appendix* Figs. S14a−b**), respectively. The C*_α_*-RMSD for mSA-free GlpG was 5 ± 1 Å and 9 ± 1 Å at 274 K and 308 K, respectively (**Fig. 5b**). In support of our steric trapping strategy, the RMSDs of the three mSA-bound GlpG versions were larger (C*_α_*-RMSD = 7–9 Å and 10–12 Å at 274 K and 308 K, respectively) with the average radius of gyration (*R*_g_) within 1.2–2.5 Å of the native value of 16 Å. Given these RMSD and *R*_g_ values, we selected the 274 K and 308 K simulations to represent the NSE (“native-like”) and a compact DSE (“collapsed”), respectively.

We ran simulations at higher temperatures in hopes of creating a prototypical expanded DSE (***SI Appendix* Fig. S14c**). At 343 K, the protein displayed significantly larger dynamics than at 308 K. However, the protein generally remained compact (*R*_g_ <24 Å) even though the C*_α_*-RMSD was often above 10 Å, indicating that the helices remained in contact but no longer retained their native arrangement (**Fig. 5b**). Notably, some trajectories underwent large excursions where TM6 unfolded and remained outside the bilayer allowing the mSA molecules to sample a much larger region of space. Nonetheless, higher temperatures increased the separation between the TM segments and broadened the corresponding distance distributions (**Figs. 5c–d**). Likewise, the attachment of doubly bound mSA preferentially increased the distance between the structural segments to which their steric repulsion was directly exerted. Based on these conformational features, we referred to this simulated model as “partially expanded DSE”.

Given the compactness at 343 K and the persistence of helix-helix contacts, we adopted an alternative strategy to create a more appropriate expanded DSE (**Figs. 5b–d** and ***SI Appendix* Fig. S14f**). Twenty additional 4 msec simulations were run retaining hydrogen bonding and the membrane potential but with the attractive side chain-side chain terms turned off, leaving only a repulsive term to avoid their steric overlap. Without interhelical interactions, we reduced the temperature to 240 K to help maintain the helices within the bilayer. Under this protocol, the TM1– TM5 helices independently diffused around in the membrane with their separation restricted by interhelical loops. As observed at 343 K, TM6 occasionally unfolded and partitioned into the surface of the membrane. The two small helices between TM1 and TM2 also unfolded, which allowed TM1 to separate more than the other helices. The overall C*_α_*-RMSD and *R*_g_ values were large, 23 Å and 30 Å, respectively. We selected this DSE for use as our reference “fully expanded DSE”.

The experimental *T*_m,app_ determined by heat-induced aggregation or inactivation is 343– 348 K in DDM micelles and above 358 K in bicelles or *E. coli* liposomes (43, 62). In our simulations, mSA-free GlpG started to melt at ∼300 K with the N-subdomain being more temperature resistant than the C-subdomain by ∼30 K (**Figs. 5b–d**). We have previously shown that the N-subdomain has a higher thermodynamic stability than the C-subdomain by *ΔΔG*°_D-N_ = 1.1 kcal/mol in micelles (42). The unfolding of TM6 at 343 K on the extracellular side observed in simulations (***SI Appendix* Fig. S14c**) agrees with our experimental result that TM6 and the flanking loops of TM5 are prone to proteolysis by ProK (**Fig. 3**).

### Comparison between DEER distance distributions and simulations

Of primary interest is the comparison between the experimental distance distributions and the simulated distance distributions for sterically denatured GlpG in bicelles and liposomes (**Fig. 6a** and ***SI Appendix* Tables S8–S10**). By this comparison, we can determine whether the DSE of GlpG is collapsed, partially expanded or fully expanded. To quantitatively describe the degree of expansion of the DSE relative to the collapsed and fully expanded states, we defined the normalized expansion ratio, *R*_Expansion_ (**Fig. 6b**):

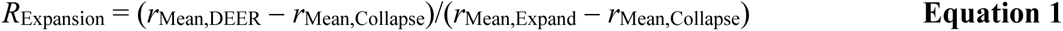

**Fig. 6:**
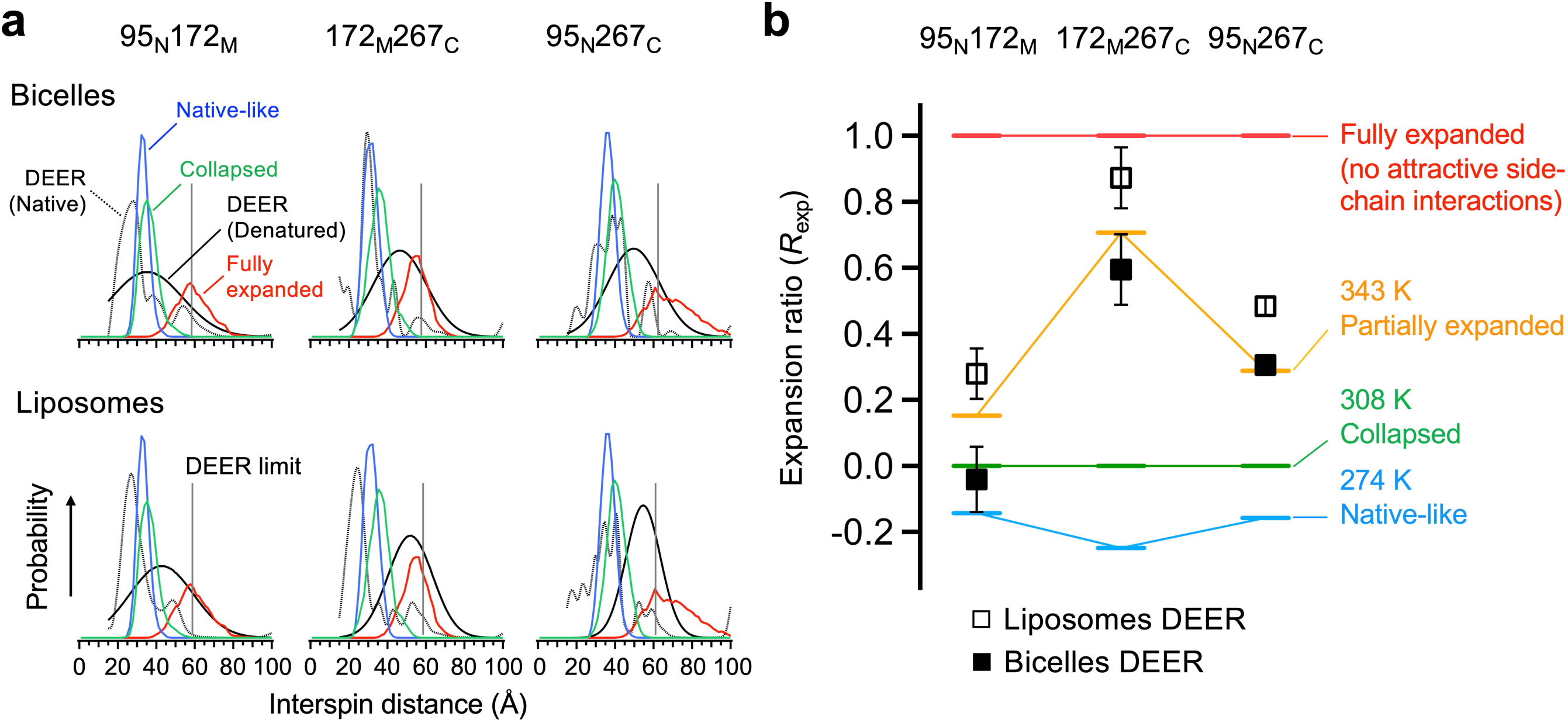
Dimension of the denatured states of GlpG measured by DEER and *Upside* simulation. **(a)** Comparison of the distance distributions obtained from DEER and the corresponding distances in the simulated DSEs doubly bound with mSA. To mimic the interspin distances measured from DEER, the distances from the simulations were measured between the Asn49 C*_α_*’s on bound mSA molecules, which were the attachment sites to GlpG. **(b)** Expansion ratio (*R*_Expansion_, **Eq.1**) of the DSE measured between each residue pair in the lipid bilayers (*i.e.*, bicelles and liposomes). The *R*_Expansion_’s of the simulated DSEs at an increasing temperature are overlayed.

where *r*_Mean,DEER_ is the mean distance between a specific residue pair in denatured GlpG determined by DEER, while *r*_Mean,Collapse_ and *r*_Mean,Expand_ denote the mean distances between the same pair in the simulated collapsed and fully expanded DSE, respectively. When the DSE was trapped using mSA at the more stable N-subdomain (95_N_172_M_) with extensive inter-residue contacts (***SI Appendix* Fig. S15**), *R*_Expansion_ was *−*0.04 ± 0.10 in bicelles, that is, this region in the DSE was as compact as the simulated collapsed state. In contrast, when the DSE was trapped using mSA at the less stable C-subdomain (172_M_267_C_), *R*_Expansion_ reached 0.59 ± 0.11, that is, the DSE expanded close to the partially expanded state. In the DSE trapped at the termini (95_N_267_C_), *R*_Expansion_ was 0.31 ± 0.02, which is likely to be an outcome of the collapse in the N-subdomain being offset by the expansion in the C-subdomain.

As noted, the DSE in the liposomes composed of *E. coli* phospholipids expanded more than in the bicelles composed of DMPC:DMPG:CHAPS throughout the protein while the N-subdomain is still more compact than the C-subdomain in liposomes (**Figs. 4** and **6a–b**): Upon denaturation, the *R*_Expansion_ values in liposomes increased by 0.32 ± 0.13 for the N-subdomain, 0.28 ± 0.14 for the C-subdomain, and 0.18 ± 0.03 for the termini from the corresponding values in bicelles. This result is consistent with the finding that the DSE was more extensively proteolyzed in liposomes than in bicelles regardless of the position of the biotin pair (**Fig. 2d**).

Given the general trends, we infer that for the DSE of GlpG, bicelles and liposomes can qualitatively behave as solvents somewhere between a poor and good solvent limits. The precise behavior depends on the specific construct and bilayer, suggesting an inherent degree of variability to solvent quality in different combinations of proteins and bilayers.

## DISCUSSION

By combining experiment and simulation, we provided a molecular-level description of the DSE of a membrane protein in a native lipid bilayer at ambient temperature. The overall picture is a partially collapsed DSE with considerable dynamics and susceptibility to proteolytic cleavage. TM helices can transiently dock against each other but also can unfold. For example, the simulations find that TM6 unfolds and exits the bilayer, a finding supported by our MS data. The comparison of the interhelical distance distributions from DEER measurements to the corresponding distances in the simulations suggests that bicelles and liposomes behave as solvents somewhere between a poor and good solvent limits, depending on the bilayer composition and the region of the protein being considered.

### Conformational diversity of the DSE

The two-stage model has served as a useful conceptual framework to describe the folding of helical membrane proteins (24). Our experimental data and simulations indicate that GlpG’s DSE is more diverse than what is expected from the canonical two-stage model, where the formation of TM helices occurs before their association. Whereas individual TM segments undergo restricted motions within the lipid bilayer, less hydrophobic segments can unfold and become solvent exposed, suggesting that the hydrophobicity of the TM segments is a key determinant of the conformational distribution in the DSEs.

In line with these results, diversity in membrane topology has been observed locally (*e.g.*, the N-terminal TM helix of an multidrug transporter EmrE) (63) and globally (*e.g.*, EmrE and lactose permease LacY) (64–68) depending on the hydrophobicity of the TM segments as well as the distribution of the positively charged residues in the flanking regions. Notably, a hydropathy analysis of the multi-spanning membrane proteins in *E. coli* predicts that about half of the TM segments have a low tendency to insert into the membrane (*i.e.*, *ΔG*_app,insertion_>0) (63). This observation combined with our current findings suggest that in general, multiple segments that are TM helices in the native state may not remain so in the DSEs of the membrane proteome.

### General principles of the DSE contraction and folding in the lipid bilayer

Another major finding of our study is that degree of contraction of GlpG’s DSE depends on the local stability in the native state. The DSE contracts upon transfer from micelles to bicelles with the more stable N- subdomain contracting as if in the poor solvent limit.

The lipid-driven contraction of the DSE reflects the balance between protein-protein, protein-lipid and lipid-lipid interactions. If the TM segments in the DSE are fully solvated by lipids (*i.e.*, protein-lipid interactions are maximal), the combined chemical potentials for lipid-lipid and protein-protein interactions should be higher (weaker) than for protein-lipid interactions due to the perturbation of the lipid bilayer and the disruption of native TM helix interactions (**Fig. 7**). Nevertheless, these two chemical self-potentials can induce partial contraction of the DSE and promote folding to the native state further releasing solvated lipids into the bulk lipids. This scenario is consistent with the general trend of lipid-lipid interactions being stronger than detergent-detergent interactions (69), which may induce compaction of GlpG’s DSE in bicelles upon transfer from micelles (**Fig. 4b**).

**Fig. 7:**
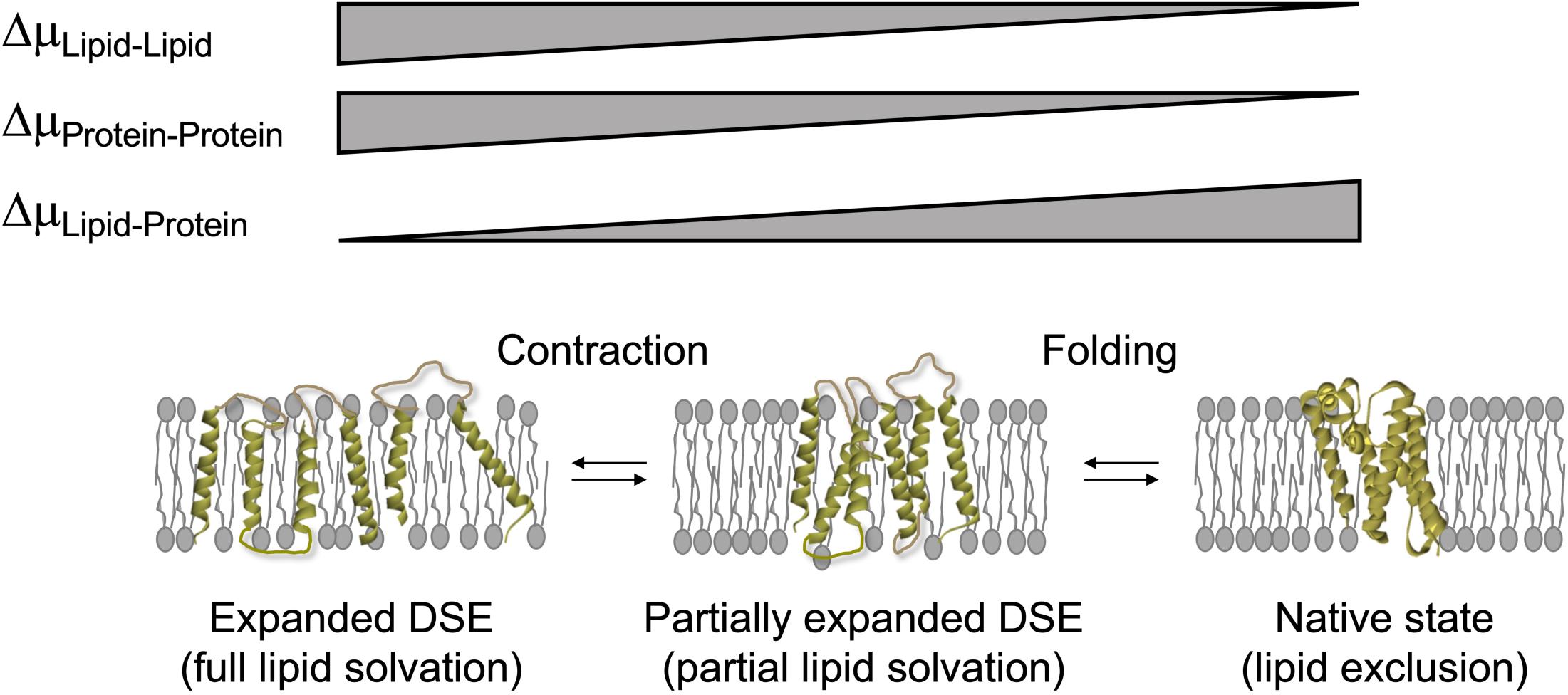
Proposed model on the denatured state contraction of helical membrane proteins in the lipid bilayers. (*Left*) The hypothetical fully expanded DSE in which the TM segments are fully solvated by lipids. (*Middle*) The partially expanded DSE in which three driving forces (*i.e.*, protein-protein, protein-lipid and lipid-lipid interactions) are balanced. (*Right*) The native state in which the lipid molecules solvating the DSE are fully excluded to the protein surface. The possible amplitude of the chemical potential (*Δμ*) of each interaction type is represented by the thickness in a trend bar during the stepwise transition from the fully expanded DSE model to the native state.

The lipid-driven compaction of the DSE may reflect a lipophobic effect (the solvophobic effect in a broader context) in the membrane, which has long been postulated but remains elusive regarding its molecular basis. In case of globular proteins, the hydrophobic effect is largely entropic, related to the release of bound waters (8, 70). In contrast, the chemical similarity between the surface of TM helices and lipid molecules implies that the lipophobic effect in the membrane may be a combination of enthalpic and entropic contributions stemming from the relatively unfavorable lipid-lipid interactions and lipid ordering, respectively, in the DSE. In both soluble and membrane proteins, however, the solvophobic effect may be insufficient to drive collapse and folding on its own (20). Favorable lipid-lipid interactions, intrachain van der Waals packing and polar interactions are needed to tip the energetic balance toward the native state.

### Lipophobic effect depends on the context of native structure and lipid environment

While we propose that the lipophobic effect generally induces protein compaction in a lipid bilayer, recent studies indicate that this solvent-dependent effect occurs through multiple mechanisms that can be modulated by the native structure and lipid environment:

*Mode 1)* Lipid deformation at the protein surface (71): For example, in the dynamic monomer-dimer equilibrium of the CLC-ec1 Cl^–^/H^+^ antiporter, a thicker lipid bilayer induces a hydrophobic mismatch between the dimer interface on each monomer and the bilayer, resulting in local thinning of the bilayer by deformation of the lipid molecules at the interface (72). This local thinning is pronounced in the monomers and shifts the equilibrium towards dimers, which can be overcome by the incorporation of lipids with shorter acyl chains that preferentially solvate the interface (72), *Mode 2)* Exposure of the polar residues in the bilayer: The isolated N- and C-domains in each CLC-ec1 monomer associate with more water molecules than the folded monomer at the domain interface due to the exposure of the polar residues in the bilayer core (73). The energetic penalty caused by the hydration of the membrane-buried domain interface can drive the folding of each monomer (73), *Mode 3)* Chemical and physical properties of the bulk lipid bilayer: TM helix-helix interactions can be modulated by lipid composition with negligible changes in the bilayer thickness for various types of single membrane-spanning helices, including the TM domain of glycophorin A (74, 75), a proton channel M2 (76), a transcriptional regulator Mga2 (77) and an endoplasmic reticulum stress sensor Ire1*α* (78). The origin of such lipid dependence has been attributed to the changes in lateral tension profile (75), lipid packing density (77), membrane fluidity (78), and the electrostatic interactions between the negatively charged membranes and the positively charged residues flanking the TM helices (75).

We find that the DSE of GlpG expands more in *E. coli* liposomes than in bicelles despite a similar quasi-2D physical constraint. Although different modes likely come into play, this difference could be explained by the different lipid composition between the two bilayer systems. Bicelles are composed of the saturated lipids, neutral *di*C_14:0_PC (75 mol-%) and negatively charged *di*C_14:0_PG (25%), and CHAPS while *E. coli* lipids mainly contain neutral PE (∼75 %), and negatively charged PG (∼20%) and cardiolipin (5%) headgroups with 16:0 (∼40%), 16:1c9 (∼30%) and 18:1c11 (∼15%) fatty acyl chains (79). Each monolayer leaflet in the bilayer containing the small headgroup and unsaturated PE or cardiolipin has a tendency to bend outward (*i.e.*, negative spontaneous curvature) compared to that containing the larger headgroup and saturated PC or PG lipids (80–84). Hence, the lipid bilayer is “frustrated” by the difference between the spontaneous and actual curvatures with an increased repulsive force between the hydrocarbon chains (lateral tension) in the bilayer core. Thus, the increase in lateral tension in the core of the *E. coli* lipid bilayer may facilitate the separation of TM segments, inducing increased lipid solvation of the TM helices and expansion of the DSE.

### Challenges in determining the solvent quality of lipid bilayers

Solvent quality has been obtained for disordered soluble proteins using the Flory scaling relationship, *R*_G_ ∝ *N^ν^*(85). This powerful formalism, however, is not readily adaptable to DSEs of membrane proteins. Although a tempting 2D counterpart would be a chain in a quasi-2D slab (86), this is a poor model of an actual DSE of a membrane protein as there are multiple TM helices positioned orthogonal to the lipid bilayer and connected by loops that largely remain in solution above the bilayer. For example, when a pair of TM helices are linked by a flexible loop of increasing length, the scaling behavior would largely be determined by that for the loop above the bilayer. Furthermore, different membrane proteins have different topologies, helical content, variable length linkers, and some helices may even lie along the bilayer. Thus, deriving generalized scaling exponents for the DSE of membranes may be difficult.

To circumvent these issues, we modelled GlpG’s DSE allowing its TM helices to dissociate and unfold with either attractive or repulsive side-chain interactions to represent poor and good solvent conditions, respectively. These simulations are used to generate reference ensembles for comparing to the DEER data and quantify compaction levels of the DSE to assess solvent quality. **Implications in quality control of membrane proteins.** The solvent quality of the lipid bilayer for membrane proteins is relevant to a variety of biological events including folding, association, chaperone action and proteostasis. Our results imply that the lipid bilayer overall facilitates contraction of DSEs of membrane proteins and generally promotes protein-protein interactions in membranes.

The adhesion of TM helices can present both challenges and benefits. TM segments make both native and nonnative contacts with each other during the co-translational insertion (26, 87, 88) while the efficiency of the cellular folding and maturation of newly synthesized membrane proteins is surprisingly small (20–50%) (89, 90). In addition, various types of TM chaperones are either components of the translocon complex or function independently (91–95). These observations imply that cellular membranes are not necessarily an optimal environment for efficient folding and assembly of membrane proteins. The bilayer may promote nonspecific compaction and misfolding, resulting in species that become targets for protein quality control mechanisms in the cells (89, 96, 97). On the other hand, protein-protein interactions are necessary for the formation of membrane protein complexes and the clustering of receptors for cellular signaling (98–101). Presumably, the interaction strength is tuned by varying lipid composition in the different bilayers in the cell to achieve “a livable compromise” in each one.

## MATERIALS AND METHODS

Detailed information on materials, preparation and biotinylation of GlpG, activity determination, mass spectrometry, and *Upside* MD simulation can be found in ***SI text***.

### Preparation of biotinylated GlpG

Double cysteine variants (95C172C and 172C267C and 95C267C) of *E. coli* GlpG with an N-terminal His_6_-tag were expressed in *E. coli* BL21(DE3) RP cells and purified in DDM (Anatrace) (42). The double cysteine variants were labeled with thiol-reactive BtnPyr or BtnRG labels as previously described (42).

### Activity assay of GlpG

We used the model substrate LYTM2–His_6_ fused to staphylococcal nuclease (SN) with a unique cysteine residue in the upstream of the scissile bond (42). To measure GlpG activity in micelles and bicelles, SN–LYTM2–His_6_ in DDM was labeled with the thiol-reactive environment-sensitive fluorophore, iodoacetyl-7-nitrobenz-2-oxa-1,3-diazol (IA-NBD, Setareh Biotech) (42). For assaying in bicelles, SN–LYTM2–His_6_ was first incorporated into bicelles (3% w/v, DMPC:DMPG:CHAPS, molar ratio = 3:1:1.4) and then mixed with GlpG incorporated in the same bicelles. Time-dependent decrease of NBD fluorescence was monitored with *λ*_Ex_ = 485 nm and *λ*_Em_ = 535 nm. To measure GlpG activity in liposomes, SN–LYTM2–His_6_ was separately labeled with 5-(iodoacetamido) fluorescein (Sigma Aldrich) or 4-dimethylaminophenylazophenyl-4’-maleimide (DABMI, Setareh Biotech). After labeling, LYTM2–His_6_ were reconstituted in *E. coli* liposomes at a 1:1 molar ratio. The proteoliposomes containing LYTM2–His_6_ were mixed with the proteoliposomes containing GlpG in 20 mM Na_2_HPO_4_, 40 mM NaCl, pH 7.5. Mixing of GlpG and LYTM2 was initiated by inducing liposomal fusion upon addition of PEG_3350_ (final 14% in w/v). For the activity assay, the increase in fluorescein fluorescence was monitored with *λ*_Ex_ = 494 nm and *λ*_Em_ = 520 nm at 37°C.

### Denaturation of GlpG using steric trapping in DDM

GlpG labeled with BtnPyr or BtnRG in DDM was incubated with a 5 times molar excess of mSA at 25°C until maximum denaturation was reached. The extent of denaturation was monitored using GlpG activity for LYTM2. Maximum denaturation was reached within 48 h for 95_N_172_M_-BtnRG_2_ and 24 h for 172 _M_267_C_- BtnRG_2_ and 95_N_267_C_-BtnRG_2_.

### Reconstitution of GlpG in bicelles

Native or sterically denatured GlpG in DDM was directly injected into 3% bicelles (DMPC:DMPG:CHAPS, molar ratio = 3:1:1.4) at the final concentrations of 5 μM GlpG and 25 μM mSA in 20 mM Na_2_HPO_4_, 40 mM NaCl, pH 7.5. Incorporation of native or denatured GlpG labeled with BtnPyr was measured using quenching of pyrene fluorescence from BtnPyr by dabcyl-DOPE (Avanti Polar Lipids) mixed at the 0.1% lipid-to-lipid molar ratio. As a negative control (*i.e.*, no incorporation to bicelles), water-soluble mSA-S83C (S83C mutation on the active subunit) labeled with pyrene was used. To be used as a positive control (*i.e.*, full incorporation in bicelles), GlpG labeled with pyrene was first reconstituted in DMPC:DMPG (molar ratio = 3:1) liposomes and then desired amount of CHAPS was added to form bicelles with the lipid-to-CHAPS molar ratio of 2.8. Pyrene fluorescence was measured with the *λ*_Excitation_ = 345 nm and *λ*_Emission_ = 390 nm.

### Reconstitution of GlpG in liposomes

Native or sterically denatured GlpG (5 μM) labeled with BtnPyr in DDM was mixed with the extruded liposomes (10 mM *E. coli* polar extract) pre-soaked with 10 mM DDM, 20 mM Na_2_HPO_4_, 40 mM NaCl (pH 7.5). Detergents were removed using Bio-Beads (Bio-Rad) followed by additional extrusion. To test incorporation in liposomes, native and denatured GlpG was reconstituted in *E. coli* liposomes with rhodamine-labeled DPPE (Avanti Polar Lipids) at 0.1% lipid-to-lipid molar ratio. The proteoliposomes were mixed with 60% (w/v) sucrose and loaded at the bottom of a 10 ml polycarbonate centrifuge tube (Beckman Coulter). A discontinuous sucrose gradient was prepared (20%, 10%, 5% and 2.5% from bottom to top). The samples were ultracentrifuged at 35,000 rpm for 2 h in a 50.4 Ti rotor. Each 50 μL fraction was solubilized in 2% (w/v) *n*-octyl-β-D-glucopyranoside (*β*-OG). Rhodamine and pyrene fluorescence from each fraction was measured at *λ*_Ex_ = 560 nm and *λ*_Em_ = 583 nm for rhodamine and *λ*_Ex_ 345 nm and at *λ*_Em_ = 390 nm for pyrene.

### Proteinase K digestion

Native or sterically denatured GlpG (5 μM) doubly labeled with BtnRG without and with mSA (25 μM) was prepared in 10 mM DDM, 3% bicelles or 10 mM liposomes. Proteolysis was carried out by Proteinase K (0.14 μg/mL, Sigma) and quenched after a specified time by permethylsulfoxide (0.1 mM). After proteolysis, bound mSA molecules were dissociated using DTT (4 mM) that cleaved the disulfide bond between cysteine and the BtnRG label bound with mSA. For mass spectrometry, native and denatured GlpG reconstituted in liposomes were solubilized by mixed detergents (2% *β*-OG and 1% DDM). Excess mSA molecules were removed by passing the sample through biotin–conjugated agarose resin (Sigma). Free thiols on GlpG were alkylated by iodoacetamide (10 mM). The reaction was quenched by DTT (10 mM) for 1 h.

### Mass spectrometry

For CZE-MS/MS, the proteolysis products of free mSA, native GlpG and sterically denatured GlpG were obtained using the single-pot, solid-phase-enhanced sample preparation method (102). After washing with 90% acetonitrile, 100 mM NH_4_HCO_3_ buffer solution (pH 8) was added to elute bound peptides. Recovered peptides were separated by CZE on an ECE-001 capillary electrophoresis system (CMP Scientific) (103). For mass detection, a Q-Exactive HF tandem mass spectrometer (Thermo Fisher Scientific) coupled to the separation capillary (∼100 cm long) was used. LC-MS/MS was carried out using a Q-Exactive interfaced with a Vanquish Flex UPLC system (Thermo Fisher Scientific). The sample was injected onto a Acquity BEH-C4 column (Waters). Initial conditions were 99% Solution A (water + 0.1% formic acid)/1% Solution B (acetonitrile + 0.1% formic acid). The subsequent LC was carried out by ramping to 99% Solution B and then returning to 1% Solution B. Proteins were ionized by electrospray operating in a positive ion mode. Data were acquired using a data-dependent MS/MS method with a scan range of 200 to 2000 *m/z*. The MS/MS data were analyzed by Byonic™ (v3.9.6, Protein Metrics) (104).

### DEER Spectroscopy

Detailed procedures for sample preparation are described in ***SI text***. Final concentration of spin-labeled GlpG in DDM, bicelles or liposomes were 50*−*100 μM. Four-pulse DEER spectroscopy data were collected on a Q-band Bruker ELEXSYS 580 spectrometer using a 10 W amplifier and an EN5107D2 probe head (Bruker Biospin). Samples were loaded into sealed quartz capillaries and flash frozen in a dry ice/acetone slurry prior to data collection at 80 K. The distance distributions were determined from fits to the background-corrected dipolar evolution data using the model-free algorithm or the Gaussian model on the LongDistances program (http://www.biochemistry.ucla.edu/biochem/Faculty/Hubbell/).

### Simulations of DSEs of GlpG

Simulations used *Upside*, our near-atomic, implicit solvent model that conducts Langevin dynamics on just the N, C_α_, and C atoms (57, 58). The amide protons and carbonyl oxygens as well as side chain beads participate in the simulations with the forces on them being back-propagated onto the N, C_α_, and C atoms to determine their new positions and velocities. The side chains are represented by a multi-position, amino acid- and directional-dependent beads. The bead position probabilities of all side chains are determined simultaneously in a single global side-chain packing step that produces the lowest free energy conformation.

The energy function used here, FF1.5, has been improved over the original function FF1.0 through the use of an updated contrastive divergence training procedure that includes more extensive sampling (***SI text*)**. We added a new membrane burial potential that dynamically accounts for the level of side-chain exposure to lipids and includes unfavorable energies for unsatisfied H-bond donors and acceptors in the membrane to allow helices to fold and unfold within the bilayer (59, 60). Side-chain burial energies are determined from the statistics of a training set of proteins and accounts for the burial depth. The membrane thickness was set to 2.88 nm as predicted by OPM (PDB: 2xov) (61).

Procedures for determining precise time and temperature scale of the *Upside* model are described in ***SI text***. Briefly, from the transition rates between folded and unfolded states (well-to-well barrier-crossing process), we estimate the time unit for barrier-crossing events to be ∼1 ps. From transition rate between Ramachandran basins in the extended state (*i.e.*, chain motions within a thermodynamic well), we estimate the time unit to be ∼0.1 ps. Using a value of 0.36 ps per Verlet step, each *Upside* time unit is 40 ns. Irrespective of the issues with defining an absolute time scale, the equilibrium population distribution is expected to be approximately correct. As the *Upside* temperature scale is in arbitrary units of *RT*, the experimental stability data at 298 K for 13 small Rosetta designed soluble proteins (105) is used to calibrate the temperature scale by identifying a value of *RT* that produces the best correlation between the simulated and experimental stabilities. The best correlation occurs when 0.87*RT* equates to *T* = 298 K. Accordingly, the simulation temperatures of *RT* = 0.7, 0.8, 0.9, 1.0, 1.1 and 1.2 correspond to 240, 274, 308, 343, 377 and 411 K, respectively. All data and analysis codes are available upon reasonable request. The full simulation package of *Upside* as well as the parameter files are available to public access on GitHub: https://github.com/sosnicklab/upside-md.

### Data Availability

All data supporting the findings of this study are available within this article and ***SI Appendix***.

## FOOTNOTES

### Author contributions

K.A.G., R.G., M.D.B., W.L.H., T.R.S. and H.H. conceived the problem and designed research. K.A.G., R.G., M.D.B., D.C., S.M., M.K., Z.Y., A.S., N.F., X.P., T.R.S. and H.H. performed research. M.D.B., D.C., A.S., N.F., X.P., A.D.J., L.S., W.L.H. and T.R.S. provided new analytical tools. K.A.G., R.G., M.D.B., D.C., S.M., M.K., Z.Y., N.F., X.P., W.L.H., T.R.S. and H.H. analyzed data. K.A.G., R.G., M.D.B., D.C., A.S., W.L.H., T.R.S. and H.H. wrote the paper.

^†^These authors equally contributed to this work.

The authors declare no conflict of interest.

## Acknowledgement

We thank the Hong lab members for critical comments. This work is supported by NIH grant R01GM118685 and the Hunt for a Cure foundation to H.H., NIH Grants GM055694, GM130122, NSF MCB-2023077 to T.R.S., the Jules Stein Professorship endowment and NIH Grants R01 EY05216, T33 EY07026, and 5P41EB001980 to W.L.H., and NIH grant R01GM125991 and NSF DBI1846913 to L.S. A.D.J. acknowledges support from Michigan AgBioResearch through the USDA National Institute of Food and Agriculture, Hatch project number MICL02474.

