## Supporting_Information for "Lipid Bilayer Induces Contraction of the Denatured State Ensemble of a Helical-Bundle Membrane Protein"

### *SI Text*

**GlpG expression and purification.** *E. coli* GlpG gene was encoded in pET15b plasmid with an N-terminal $His_6$-tag as previously described (1). GlpG was expressed in the *E. coli* BL21(DE3) RP strain. Cells were grown at 37°C until $OD_{600nm}$ reached 1.0. Protein expression was induced with 0.5 mM isopropyl β-thiogalactopyranoside (IPTG, GoldBio), followed by additional cultivation at 15°C for 16 h. GlpG was isolated from the total membrane fraction obtained by ultracentrifugation (Beckman Coulter, Type 45 Ti rotor, 50,000*g*, for 2 h) using $Ni^{2+}$-nitrilotriacetic acid (NTA) affinity chromatography (Qiagen) after solubilization with 2% (w/v) *n*-dodecyl-β-D-maltoside (DDM, Anatrace).

**Biotin labeling of GlpG.** The stock solution of the double cysteine variants (95C172C, 172C267C and 95C267C) of GlpG in DDM was diluted to ~50 μM in 0.2% DDM, 50 mM tris-(hydroxymethyl) aminomethane hydrochloride (TrisHCl, Fisher Scientific) (pH 8.0), 200 mM NaCl, and incubated with a ten times molar excess of tris (2-carboxyethyl) phosphine hydrochloride (TCEP-HCl, Pierce) for 1 h at room temperature. A 40 times molar excess of BtnPyr-IA or BtnRG-TP dissolved in dimethyl sulfoxide (DMSO) was added to the mixture while vortexing. Labeling reaction was incubated at room temperature overnight in the dark with gentle stirring. After labeling with BtnPyr-IA, excess free labels were removed by extensive washing of the proteins bound to $Ni^{2+}$-NTA affinity resin using 0.1% DDM, 50 mM TrisHCl (pH 8.0), 200 mM NaCl solution followed by dialysis against 0.02% DDM, 50 mM TrisHCl (pH 8.0), 200 mM NaCl to remove imidazole. After labeling with BtnRG-TP, excess free labels were removed running desalting column (Bio-rad) twice equilibrated with 0.1% DDM, 50 mM TrisHCl (pH 8.0), 200 mM NaCl. The samples were concentrated using a centrifugal concentrator (Millipore, MWCO=30 kD). The final protein concentration was determined using DC protein assay (Bio-Rad). For the BtnPyr label, the labeling efficiency of GlpG was determined using the protein concentration determined by DC assay and the pyrene concentration determined by absorbance at 346 nm ($\varepsilon_{Molar}$ = 43,000 $M^{-1}cm^{-1}$). For the BtnRG label, the labeling efficiency of GlpG was determined using SDS-PAGE gel shift assay (in the absence of reducing agents) as previously described (***SI Appendix*** **Fig. S1**) (1). Briefly, the band intensities that corresponded to GlpG singly bound with mSA (monovalent streptavidin) and GlpG doubly bound with mSA were compared accounting for the molecular mass of GlpG and mSA. The labeling efficiency typically ranged from 1.5 to 1.8 per GlpG. The detailed procedures for preparing mSA are described in the literature (1, 2).

**Bicelle preparation.** 15% (w/v) stock of bicelles composed of DMPC (1,2-dimyristoyl-*sn*-glycero-3-phosphocholine)/DMPG (1,2-dimyristoyl-*sn*-glycero-3-phospho-(1'-rac-glycerol))/CHAPS (3-[(3-Cholamidopropyl) dimethylammonio]-1-propanesulfonate) (lipid-to-detergent molar ratio, $q$ = 2.8) were prepared by hydrating DMPC/DMPG (molar ratio = 3:1) lipids with water. 20% (w/v) CHAPS was added to reach the desired $q$ value. Bicelle samples were homogenized through three cycles of freeze-thaw using liquid $N_2$ and a water bath at 42°C. Bicelle stocks were kept at –20°C prior to use.

**Transfer of native and denatured GlpG to bicelles.** GlpG labeled with either BtnPyr or BtnRG in DDM was incubated with a 5 times molar excess of mSA at room temperature until maximum denaturation was reached. The extent of denaturation was monitored using the proteolytic activity of GlpG for the transmembrane (TM) substrate, LYTM2 (the second TM segment of *E. coli* lactose permease LacY), monitored every 24 h. Maximum denaturation was reached within 48 h for $95_N172_M$-BtnR$G_2$ and 24 h for 172 $_M267_C$-BtnR$G_2$ and $95_N267_C$-BtnR$G_2$. Native and denatured GlpG were directly injected into preformed bicelles to the final concentrations of 5 μM GlpG, 25 μM mSA, and 3% (w/v) DMPC/DMPG/CHAPS bicelles in 20 mM $Na_2HPO_4$, 40 mM NaCl, pH 7.5 and incubated overnight at room temperature.

**Measuring incorporation of native and denatured GlpG into bicelles.** 7.5% bicelles containing dabcyl-DOPE (quencher-labeled lipid, Avanti Polar Lipids) at the 0.1% lipid-to-lipid molar ratio were prepared in 20 mM HEPES buffer (pH 7.5). Double cysteine GlpG variants were labeled with BtnPyr-IA as described above. Incorporation of native or denatured GlpG into bicelles was measured using quenching of pyrene fluorescence from GlpG by the dabcyl label localized in the bilayer region in bicelles.

As a negative control (*i.e.*, no incorporation to bicelles), highly water-soluble mSA-S83C (S83C mutation was made on the active subunit) labeled with pyrene was used. mSA-S83C was labeled using the following procedures: 1 mL of 30 μM mSA-S83C in 0.3 mM TCEP, 20 mM HEPES buffer (pH 8.0) was incubated with a 20 times molar excess of thiol-reactive N-(1-pyrene) maleimide solubilized in DMSO for 2 h at room temperature. Excess free labels were removed on a desalting column equilibrated with 20 mM HEPES buffer (pH 7.5). To the final pyrene-labeled mSA stock, DDM was added to a final concentration of 5 mM to match the DDM concentration in GlpG stock.

To be used as a positive control (*i.e.*, full incorporation in bicelles), GlpG labeled with pyrene was first reconstituted in DMPC/DMPG liposomes using the following procedures: Mixed dried lipid ([DMPC]:[DMPG] = 3:1) was resuspended in 20 mM HEPES buffer (pH 7.5) to a final lipid concentration of 4% (w/v). The lipid suspension was homogenized by three cycles of freeze-thaw and then extruded through 0.2 μm pore-size polycarbonate membrane (Whatman). DDM was added to the liposome suspension to a final concentration of 40 mM and incubated for 30 min. Then, GlpG labeled with BtnPyr from a stock solution was added to a final concentration of 10 μM. The lipid-protein-detergent mixture was incubated for 30 min. Three portions of Bio-Beads (Bio-Rad) were added (20 mg/mL for each) stepwise to remove detergent DDM. In the first removal step, the samples were gently stirred at 4 °C for 2 h and then moved to room temperature in the subsequent removal steps (1–2 h of incubation in each step). The resulting proteoliposomes were extruded again using 0.2 μm pore size membrane. The total phospholipid concentration was determined using an organic phosphate assay. Based on the measured total lipid concentration, a desired amount of CHAPS was added to form bicelles with $q = 2.8$. Then, the 7.5% bicelle stock containing dabcyl-labeled lipid (see above) was added to the final bicelle concentration of 3%, during which the bicelle constituents (labeled and unlabeled lipids and GlpG) are homogeneously mixed.

In the samples for negative and positive controls, the final pyrene and dabcyl concentrations were matched to those of the experimental samples (see below).

To be used as experiment, native or sterically trapped denatured GlpG labeled with BtnPyr in DDM was directly injected into preformed 7.5% bicelles containing DOPE-dabcyl at the final concentrations of 3% bicelles and the final pyrene concentration of 5 μM as measured by UV-Vis absorbance at 346 nm ($\varepsilon_{Molar}$ = 43,000 $M^{-1}cm^{-1}$). After mixing, the samples were equilibrated overnight at room temperature.

Pyrene fluorescence from the control and experiment samples was measured on a 96-well plate using SpectraMax M5e plate reader (Molecular Devices) with the excitation and emission wavelengths of 345 nm and 390 nm, respectively. The ratio of the pyrene fluorescence intensities from the experimental and positive control samples to the intensity from the negative control sample, respectively, was used as a measure of GlpG incorporation to bicelles.

**Preparation of *E. coli* liposomes.** Dried *E. coli* lipid (Avanti Polar Lipids) films were hydrated with 20 mM $Na_2HPO_4$ (pH 7.5), 40 mM NaCl buffer to a final lipid concentration of 10 mM. The lipid suspension was homogenized by three cycles of freeze-thaw and then extruded through a 0.2 μm pore size polycarbonate membrane.

**Transfer of native and denatured GlpG into *E. coli* liposomes.** DDM was added to empty *E. coli* liposomes (10 mM lipid extruded through a 0.2 μm pore size polycarbonate membrane) to a final concentration of 10 mM (detergent-to-lipid molar ratio = 1) and incubated for 30 min. Native or sterically denatured GlpG in DDM (preparation procedures are described in the subsection "**Transfer of native and denatured GlpG to bicelles.**") was added to the DDM-lipid mixture to a final concentration of 5 μM. The lipid-protein-detergent mixture was incubated for 30 min. For detergent removal, three portions of Bio-Beads (Bio-Rad) were added (20 mg/mL for each) stepwise. In the first removal step, the samples were gently stirred at 4 °C for 2 h and then moved to room temperature in the subsequent removal steps (1−2 h of incubation in each step). The resulting proteoliposomes were extruded using a 0.2 μm pore size membrane.

**Flotation assay of proteoliposome samples.** Pyrene-labeled GlpG ($95_N172_M$–$BtnPyr_2$, $172_M267_C$–$BtnPyr_2$ or $95_N267_C$–$BtnPyr_2$) was reconstituted in *E. coli* liposomes containing rhodamine-labeled lipid (DPPE-Rho, 0.1% lipid-to-lipid molar ratio, Avanti Polar Lipids). The proteoliposomes containing GlpG (50 μL) was mixed with 60% (w/v) sucrose in 20 mM HEPES (pH 7.5) (50 μL). The mixture was loaded at the bottom of the centrifuge tube (polycarbonate tubes, 1 mL capacity, Beckman Coulter) and flash-frozen in liquid $N_2$. A discontinuous sucrose gradient was prepared by the addition and subsequent flash-freezing of the solution of a lower sucrose concentration (100 μL of 20%, 10%, 5% and 2.5%). After thawing, the tube was centrifuged at 35,000 rpm at 4°C for 2 h in a fixed angle rotor 50.4 Ti (Beckman Coulter Optima XE- 90 ultracentrifuge). The tubes were taken out carefully and each ~50 μL fraction was taken from top to bottom. The fractions were solubilized in 2% (w/v) *n*-octyl-β-D-glucopyranoside (β-OG). Rhodamine and pyrene fluorescence from each fraction was measured at the excitation wavelength of 560nm and 345nm and at the emission wavelength at 583 nm and 390 nm, respectively. The amount of protein in each fraction was also analyzed using SDS-PAGE (***SI Appendix*** **Fig. S5a**). 25 μL sample was taken out from each fraction, solubilized with 2% β-OG, and then mixed with the SDS sample loading buffer.

**Sodium carbonate extraction.** There were three liposome samples for each GlpG variant: native GlpG in *E. coli* liposomes, sterically denatured GlpG in *E. coli* liposomes and empty *E. coli* liposomes mixed with water-soluble mSA-WT as a reference (***SI***

***Appendix* Fig. S5b**). 50 μL of each sample was incubated with 500 μL of pre-chilled 0.1 M $Na_2CO_3$ buffer (pH 11.0) for 30 min on ice. Then, the mixture was ultracentrifuged at 4 °C for 30 min at 90,000*g* in Beckman polycarbonate tubes (4 mL tube capacity) in a 50.4 Ti rotor. Separated supernatants and pellets were incubated in 2.5 mL or 0.5 mL of 12.5% (w/w) trichloroacetic acid for at least 15 min on ice to precipitate all the protein content, followed by ultracentrifugation for 30 min at 28,000*g* at 4 °C in a fixed angle rotor 50.4 Ti (Beckman Coulter Optima XE-90 ultracentrifuge). All the pellets after the last centrifugation were first solubilized in 3% (w/v) β-OG, followed by the addition of SDS sample buffer for SDS-PAGE.

**Expression and purification of GlpG substrate SN-LYTM2.** For GlpG activity assays, we used the second transmembrane segment of lactose permease (LYTM2) as the model substrate of GlpG. LYTM2 was fused to staphylococcal nuclease (SN) connected by a linker with a TEV protease recognition site. The resulting DNA construct SN-TEV-LYTM2-$His_6$ possesses a unique cysteine residue, which is located in the upstream of the scissile bond. The procedures for expression and purification of this construct in BL21(DE3) RP *E. coli* strain have previously been described (1). For measuring GlpG activity in DDM micelles and DMPC/DMPG/CHAPS bicelles, SN-TEV-LYTM2-$His_6$ was labeled with the thiol-reactive environment-sensitive fluorophore iodoacetyl-7-nitrobenz-2-oxa-1,3-diazol (IA-NBD amide, Setareh Biotech) (1).

To measure GlpG activity in *E. coli* liposomes (the principle of the assay is described in ***SI Appendix* Fig. S6**), SN-TEV-LYTM2-$His_6$ was labeled with either 5-(iodoacetamido) fluorescein (Sigma Aldrich) or 4-dimethylaminophenylazophenyl-4'-maleimide (DABMI, Setareh Biotech). After labeling, TEV-protease was added to cleave off the SN domain at the TEV cleavage site, leaving LYTM2-$His_6$ with the conjugated fluorophores. TEV protease and SN were removed using $Ni^{2+}$-NTA chromatography.

**Monitoring proteolytic activity of GlpG in micelles, bicelles and liposomes.** The activity assay in micelles or bicelles was initiated by the addition of a 10 times molar excess of the model substrate, NBD-labeled SN-LYTM2 to GlpG in 20 mM $Na_2HPO_4$ (pH 7.5), 40 mM NaCl. Time-dependent decrease of NBD fluorescence, which was a measure of proteolytic activity, was monitored on a 96-well plate using SpectraMax M5e plate reader (Molecular Devices) with the excitation and emission wavelengths of 485 nm and 535 nm, respectively. The change in fluorescence intensity was normalized to a control sample containing NBD-SN-LYTM2 alone. For activity measurement in bicelles, SN-LYTM2 and GlpG were separately incorporated into 3% DMPC/DMPG/CHAPS bicelles by direct injection and the proteolysis reaction was initiated by mixing the bicelle samples with GlpG- and SN-LYTM2.

To measure GlpG activity in liposomes (***SI Appendix* Fig. S6**), LYTM2 labeled with fluorescein and DABMI were incorporated into liposomes composed of *E. coli* phospholipids (Avanti Polar Lipids) at a 1:1 molar ratio with the total substrate concentration of 50 μM and the total lipid concentration of 5 mM. The reconstitution was performed using the following procedures: 5 mM preformed *E. coli* liposomes were incubated with 5 mM DDM at room temperature for 30 minutes. Then 25 μM $LYTM2_{DAB}$ and 25 μM $LYTM2_{FL}$ where added while vortexing, followed by incubation at room temperature for 30 min. For detergent removal, three portions of Bio-Beads (Bio-

Rad) were added (200 mg/mL for each) stepwise. In each step, the mixture was gently stirred for 1–2 h at room temperature. The resulting proteoliposomes were extruded using a 0.2 μm pore size membrane to remove aggregation.

For activity assay, the proteoliposomes containing LYTM2 (10 μL) were mixed with the proteoliposomes containing 5 μM GlpG (5 μL) and 18.5 μL of buffer (20 mM $Na_2HPO_4$, 40 mM NaCl, pH 7.5). Fusion of proteoliposomes was initiated by the addition of 16.5 μL 36% $PEG_{3350}$ containing 365 mM NaCl. Time-dependent change of fluorescein fluorescence was monitored at 37 °C on a 96-well plate using SpectraMax M5e plate reader with the excitation and emission wavelengths of 494 nm and 520 nm, respectively. Fluorescence increase, which is caused by dequenching of fluorescein fluorescence upon cleavage, was normalized to a control sample, the proteoliposomes containing LYTM2 mixed with the liposomes without GlpG.

**Liposome fusion assay induced by PEG.** This assay was for obtaining the time scale of mixing between the enzyme GlpG and the substrate LYTM2, which forms a basis for our GlpG activity assay in liposomes (see above). We employed a FRET-based lipid mixing assay (3). To prepare the proteoliposomes containing the substrate, Cys-less LYTM2 was reconstituted in *E. coli* liposomes containing 0.02 molar fraction of *N*-(7-nitro-2,1,3-benzoxadiazol-4-yl)(ammonium salt) dipalmitoylphosphatidylethanolamine (DPPE-NBD, FRET donor) and 0.02 molar fraction of quenching lipid *N*-(lissamine rhodamine B sulfonyl)(ammonium salt) dipalmitoylphosphatidylethanolamine (DPPE-Rho, FRET acceptor) to the final substrate concentration of 50 μM and the final lipid concentration of 5 mM. GlpG was reconstituted in *E. coli* liposomes without fluorescent label to the final protein concentration of 5 μM and the final lipid concentration of 5 mM. All the samples were prepared in 20 mM HEPES (pH 7.5) and 200 mM NaCl. The protein/lipid molar ratio was adjusted to mimic that in the activity assays described above.

PEG-induced liposome fusion was detected upon lipid mixing between fluorescently labeled (20 μL) and unlabeled liposomes (9.5 μL), which led to dequenching of NBD-fluorescence caused by spatial separation of NBD and Rho within the lipidic phase. The fusion reaction was initiated upon addition of 11% (v/v, final concentration) $PEG_{3350}$. The total volume was 1.4 mL in a Hellma florescence cuvette. NBD Fluorescence was detected with an excitation wavelength at 467 nm and an emission wavelength at 530 nm as a function of time with a 5 s interval (PTI QW4 fluorimeter) with constant stirring at 37°C.

As a negative control that represents no fusion, no PEG was added. As a positive control for a homogeneously mixing state, 12 μL of 100% Triton X-100 was added to a final concentration of 0.08% (w/v) to solubilize the liposomes.

**Proteinase K digestion.** 5 μM GlpG ($95_N172_M$-$BtnRG_2$, $172_M267_C$-$BtnRG_2$ or $95_N267_C$-$BtnRG_2$) in the absence and presence of 25 μM mSA was prepared in 10 mM DDM, 10 mM DMPC/DMPG/CHAPS bicelles and 10 mM *E. coli* liposomes, as described above. 2 mM $CaCl_2$ was added to enhance the stability of proteinase K (Sigma). Proteolysis was initiated upon addition of 0.14 μg/mL proteinase K (final concentration). An aliquot of each sample was taken at a specified time, and the reaction was quenched by the addition of 10 mM phenylmethylsulfonyl fluoride (PMSF). For post-proteolysis removal of bound mSA molecules that had been added to trap the denatured state of GlpG, 4 mM dithiothreitol (DTT) was added to cleave the disulfide bond that links BtnRG label bound with mSA to the cysteine residues. These samples were directly used for SDS-PAGE upon addition of the SDS sample buffer. For mass spectrometry, native and sterically denatured GlpG reconstituted in liposomes were solubilized by the addition of mixed micelles (2% β-OG and 1% DDM). Denatured GlpG which had been digested contained dissociated mSA. Excess mSA molecules were removed by passing the sample through biotin-agarose resin (Sigma). Free thiol groups on GlpG were alkylated by the addition of 10 mM iodoacetamide (final concentration) and incubation for 60 min in the dark at room temperature. The reaction was quenched by the addition of 10 mM DTT and incubation for 1 h at room temperature.

**Mass spectrometry.** Prior to tandem mass spectrometry coupled to capillary-zone electrophoresis (CZE-MS/MS), the proteolysis products of free mSA, native GlpG and sterically denatured GlpG were desalted using the single-pot, solid-phase-enhanced sample preparation (SP3) (4, 5). The carboxylated paramagnetic beads, which are a 1:1 mixture of hydrophilic and hydrophobic Sera-Mag SpeedBeads (Sigma-Aldrich), were washed with water three times. 10 μL of each proteolyzed sample was diluted with 60 μL of acetonitrile. 5 μL of the bead mixture was added to each proteolyzed sample, followed by gentle vortexing and incubation for 20 min. A magnet was placed under the sample tube for 1 min and then the supernatant was removed. Beads were washed by 90% acetonitrile twice. 3 μL of 100 mM $NH_4HCO_3$ buffer solution (pH 8) was added to the beads to elute bound peptides. The samples were stored in –20°C before CZE-MS/MS.

Recovered peptides were separated by CZE on an ECE-001 capillary electrophoresis system (CMP Scientific). To reduce the electroosmotic flow during CZE, the capillary was coated with linear polyacrylamide (LPA) (6). A Q-Exactive HF tandem mass spectrometer (Thermo Fisher Scientific) was used for mass detection. The mass spectrometer was coupled to the separation capillary by a commercialized electrokinetically pumped sheath-flow CE–MS interface (CMP Scientific). A ~100 cm long LPA-coated capillary was used for separation. Samples were injected using 5 psi for 45 s and separated at +30 kV. After 90 min of the separation time, 30 psi was applied for 10 min to flush the capillary.

A top-10 data-dependent acquisition strategy was employed for MS. The NCE (normalized collisional energy) was set to 28 V for peptide fragmentation. For MS, the microscan, resolution, automatic gain control (AGC) target and maximum injection time were set to 1, 60,000, $3 \cdot 10^6$ and 50 ms, respectively, with the scan range of 300 to 2,000 *m/z*. For MS/MS, the microscan, AGC target and maximum injection time were set to 1, 30,000, $10^5$ and 200 ms, respectively with the scan range of 200 to 2,000 *m/z*. The isolation window was set to 2.0 *m/z* and the dynamic exclusion was set to 30 s. Ions with unassigned charge, or with the charge equal to or larger than 8 were excluded for fragmentation.

Protein samples were analyzed by LC/MS/MS using a Thermo Q-Exactive interfaced with a Thermo Vanquish Flex UPLC system. 10 μL of sample was injected onto a Waters Acquity BEH-C4 protein column (2.1x100mm). Initial conditions were 99% Solution A (water + 0.1% formic acid) / 1% Solution B (acetonitrile + 0.1% formic acid). Hold for one min at 1% Solution B, then ramp to 99% Solution B at 7.0 min, hold at 99% Solution B until 8.5 min, return to 1% Solution B at 8.51 min and hold at 1% Solution B until 10.0

min. Flow rate was 0.3 ml/min and column temperature was 30°C. Proteins were ionized by electrospray operating in positive ion mode with a capillary voltage of 3.5 kV, capillary temperature was 256°C, probe heater was 412.5 °C, and S-lens RF level was 50.0. Data were acquired using a data-dependent MS/MS method with a survey scan range of 300–4,000 *m/z*, the resolution of 140,000 at *m/z* = 200, the AGC target of $10^6$ and the maximum inject time of 100 ms. The top 3 ions were selected for MS/MS with the resolution of 35,000 at *m/z* = 200, the AGC target of $10^6$, the maximum inject time of 100 ms, the isolation width of 3.0 Da and the scan range of 200–2,000 *m/z*. Fragmentation in the HCD cell was carried out using a stepped NCE method with 10.0, 30.0 and 60.0 V. Charge states of +1 or +2 were excluded for MS/MS picking and a dynamic exclusion was set to 3.0 s.

The raw files were analyzed by Byonic™ (version 3.9.6, Protein Metrics) (7). The database contains three protein sequences (the double cysteine variant $95_N172_M$, $172_M267_C$ and mSA) in the FASTA format. All settings were as defaulted except that nonspecific digestion was chosen as the digestion specificity and the carbamidomethylation on cysteine was set as a variable modification. We calculated the cumulative Byonic score at each P1 site (*i.e.*, the preceding residue of the hydrolyzed peptide bond) (7). The Byonic score, which is the primary indicator of PSM (Peptide-Spectrum Match) correctness, reflects the absolute quality of the peptide-spectrum match, not the relative quality compared to other candidate peptides (7). When a Byonic score was obtained for an identified peptide, the score was assigned at the P1 site at each end (N- or C-term) of the peptide. Then, the scores were summed up at each P1 site for the peptides sharing the same P1 site in either end. Therefore, the P1 site with a higher score means more frequently cleaved site with more reliability in PSM assignment.

**Sample preparation for DEER** To obtain the sterically denatured state in DDM micelles, 120 µL of GlpG variant $95_N172_M$–BtnRG$_2$, $172_M267_C$–BtnRG$_2$ or $95_N267_C$–BtnRG$_2$ (25 µM) was incubated with a 5 times molar excess of mSA-WT in 40 mM DDM, 20 mM $Na_2HPO_4$ (pH 7.5), 40 mM NaCl at room temperature for three days ($95_N172_M$–BtnRG$_2$) or overnight ($172_M267_C$–BtnRG$_2$ and $95_N267_C$–BtnRG$_2$). Then, the samples were concentrated to about ~50 µL using an Amicon centrifugal concentration filter unit (0.5 ml capacity, MWCO = 10 kDa, Millipore). Glycerol was added to the final concentration of 10% (v/v). Native GlpG samples were obtained in the same way but without addition of mSA-WT.

Native and sterically denatured GlpG samples were prepared in 3 % (w/v) DMPC/DMPG /CHAPS bicelles as described above (see the subsection, **Transfer of native and denatured GlpG to bicelles)**. The samples were then concentrated using a centrifugal concentrator unit (MWCO = 10 kD) and diluted in 20 mM $Na_2HPO_4$ (pH 7.5), 40 mM NaCl, 10% (v/v) glycerol. Final concentrations of GlpG were typically 40–70 µM.

Native and sterically denatured GlpG samples (5 µM) were first prepared in micelles and transferred to *E. coli* liposomes as described above (see the subsection, **Transfer of native and denatured GlpG into *E. coli* liposomes)**. To suppress unwanted inter-molecular dipolar coupling between spin-labeled GlpG in DEER measurement, the lipid concentration was doubled to 20 mM and a 3 or 6 times molar excess of GlpG (inactive variant S201A) was mixed with spin-labeled GlpG in DDM prior to the addition to *E. coli* liposomes. After detergent removal by Biobeads and extrusion, samples were concentrated by spinning down the proteoliposomes using a fixed angle rotor 50.4 Ti at 35,000 rpm for 2 hours. The resulting pellets were resuspended in 20 mM $Na_2HPO_4$ (pH 7.5), 40 mM NaCl, 10% (v/v) glycerol. Final concentrations of spin-labeled GlpG were typically 40–60 μM.

**DEER data acquisition and analysis** Four-pulse DEER spectroscopy data were collected on a Q-band Bruker ELEXSYS 580 spectrometer using a 150 W TWT amplifier (Applied Engineering Systems, Fort Worth, TX) and an E5106400 cavity resonator (Bruker Biospin). Pulse lengths were optimized via nutation experiment but ranged from 12 to 22 ns (π/2) and 24 to 44 ns (π); Observer frequency was set to a spectral position 2 G downfield of the low and central resonance intersection point, and the pump envelope frequency was a 50 MHz-wide square-chirp pulse (generated by a Bruker arbitrary waveform generator) set 70 MHz downfield from the observer frequency.

Each 20–50 μL sample of 50–100 μM double-cysteine variants of GlpG labeled with BtnRG was placed in a 1.4/1.7 mm (inner/outer diameter) quartz capillary jacketed in a 2.0/2.4 mm borosilicate capillary (Vitrocom, Mountain Lakes, NJ) and then flash frozen in liquid nitrogen prior to data collection at 50 K. Sample temperature was maintained at 50 K by a recirculating/closed-loop helium cryocooler and compressor system (Cold Edge Technologies, Allentown, PA).

The experimental distance distributions were determined from fits to the background-corrected dipolar evolution data using the non-negative Tikhonov (*i.e.*, model-free) algorithms or the single-Gaussian model in LongDistances v.593, a custom program written by Christian Altenbach in LabVIEW (National Instruments). The software is available online (http://www.biochemistry.ucla.edu/biochem/Faculty/Hubbell/) and described elsewhere (8).

As aforementioned, dipolar evolution data are processed to yield distance probability distributions representative of all interacting spins in the 15–70 Å range (9). An important feature of the temporal dipolar evolution function is the “modulation depth”, which is the difference between the maximum and equilibrium amplitudes of the dipolar evolution function. For interacting spin pairs (as opposed to systems where $3^+$ spins interact), the modulation depth is *equal to* the experimental “inversion efficiency”, which is proportional to various instrumental and sample parameters, including the number of interacting spin pairs within the detectable range. Thus, in the analysis and comparison of modulation depths for different samples, spectrometer frequency, pulse-nutation efficiency, sample labeling efficiency, sample concentration, and distance distribution of spin pairs must be taken into account. Holding certain experimental parameters constant (spectrometer frequency, nutation efficiency, and ostensibly labeling efficiency) allows us to consider the effects of relative concentration and fraction of spin pairs outside the detectable range on recorded modulation depth. For our Q-band instrument, an interacting spin pair system reports a standard maximum modulation depth of ~0.50 for a completely labeled, optimally-excited sample. Modulation depths reported herein that are smaller than this value, indicate sub-optimal labeling efficiency, concentration effects, or a population fraction of inter-spin interactions beyond the 70 Å detection limit.

**Simulations of native and unfolded GlpG.** Simulations used our *Upside* model (10, 11) that conducts Langevin dynamics only on the N, $C_\alpha$, and C atoms, but the energy function includes hydrogen bonds between the NH and C=O groups as well as those with the side chains (*e.g.*, helix capping). We employ trained neighbor- and residue-dependent ($\phi$, $\psi$) torsion maps. The side chain is represented by a multi-position, amino acid- and directional-dependent bead. All the side chain bead positions are determined in a global side chain packing calculation using Belief Propagation. This step greatly reduces side chain friction, which along with the lack of explicit solvent, explains much of the $10^3$−$10^4$-fold speed up compared to standard MD.

The energy function used here, FF1.5, is improved over the original function FF1.0 through the use of an updated contrastive divergence training procedure that includes more extensive sampling. Critical to the success of *Upside* is the development of a force field having the proper balancing of energy terms. This is achieved by simultaneously training nearly all parameters using our version of the machine learning contrastive divergence method. Here one considers two ensembles, the first restrained to be near the native structure and the second that is free to diffuse away during simulations. For a perfect energy function, the unrestrained training ensemble should remain close to the native ensemble. With an imperfect function, however, differences arise, but they can be corrected. That is, if too many hydrogen bonds form in the training ensemble, the hydrogen bond energy is reduced, and the simulations are rerun to create a new training ensemble. This iterative procedure continues until no energy parameter can be updated to generate a training ensemble that better matches the native ensemble.

In our original training procedure, the simulation time used to produce the training ensemble was short (4000 *Upside* time steps, ~ 10 cpu-minutes). With such short simulation times, the training ensemble stayed largely in the native well, which compromised its ability to adequately challenge the force field which is needed improve the parameters. To overcome this deficiency, we generated new training ensembles with non-native folds by starting the simulations from a fully unfolded conformation that collapsed into nonnative folds. These states were diversified using metadynamics to find 12 diverse low energy structures. Together with the native structure, these were used as initial states for temperature replica exchange. We then conducted the contrastive divergence procedure using the conformations in the lowest temperature replica as the training ensemble.

We recently integrated a new knowledge-based membrane burial potential into that accounts for the changing level of side chain exposure to lipids (thereby correcting for the replacement of lipid-protein interactions by protein-protein interactions as helices approach) (12). The membrane burial potential accounts for changes in bilayer thickness and includes unfavorable energies for unsatisfied hydrogen bond donors and acceptors in the membrane, which allows helices to fold and unfold within the bilayer during the simulations. Energies are determined from the statistics of a large training set of proteins and account for both the depth Z in the membrane and the level of side-chain exposure to the lipid, i.e., $E$(Z, exposure) ∝ $RT$ ln(frequency). We also incorporate depth-dependent energies for unsatisfied backbone hydrogen bond donors and acceptors within the bilayer.

The membrane thickness was set to 28.8 nm as determiend by OPM (Orientations of Proteins in Membrane) for GlpG (PDB: 2xov) (13).

We use a Verlet integration with a time unit of 0.009 *Upside* time steps and random number generator to implement the Langevin dynamics with a thermalization time scale of 5-time units. The time scale of thermalization (related to Langevin friction) is chosen to maximize the effective diffusion rate of chains while effectively controlling simulation temperature. As Langevin dynamics with any friction coefficient produces the same Boltzmann ensemble, we chose to maximize equilibration of our system rather than attempt to match a solvent viscosity.

The precise time and temperature scale of the *Upside* model is unclear because of the coarse graining process. As compared to all-atom, explicit solvent simulations on a decapeptide using Charmm36, *Upside* is about 10,000-fold faster in part due to the lack of solvent and the smoothing of side chain interactions. This smoothing is likely to have a disproportionate effect for condensed structures as compared to extended structures. From the transition rates between folded and unfolded states (well-to-well barrier crossing process), we estimate the time unit for barrier crossing events to be ~1 ps. From transition rate between Ramachandran basins in the extended state (chain motions within a thermodynamic well), we estimate the time unit to be ~0.1 ps. Using a value of 0.36 ps per Verlet step, each *Upside* time unit is 40 ns. In the GlpG simulations, every 50th conformation is stored giving a final output spacing of 2 μs/frame. Irrespective of the issues with defining an absolute time scale, the equilibrium population distribution that determines the free energy is expected to be approximately correct, as well as the order of dynamical folding events.

The *Upside* temperature scale was calibrated using 13 Rosetta designed mini-proteins that *Upside* can reversible fold and having known stability at 298 K (chemical denaturation measurements) (14). Temperature replica exchange simulations were run across a broad enough temperature range to generate melting curves for each protein. These melting curves (hydrogen bonds versus temperature) were fit assuming a two-state model between the native and unfolded state to obtain $\Delta G_{\text{sim}}$. As *Upside* temperature scale is in arbitrary units of $RT$, the experimental stability data is used to calibrate the temperature scale by identifying a value of $RT$ that produces the best correlation between the simulated and experimental stabilities at 298 K. For the test set, the best correlation occurs when 0.87 $RT$ equates to $T = 298$ K. Accordingly, the simulation temperatures of $RT = 0.7$, 0.8, 0.9, 1.0, 1.1 and 1.2 $RT$ correspond to 240, 274, 308, 343, 377 and 411 K.

All data and analysis codes supporting the findings of this study are available from the corresponding authors upon reasonable request. The full simulation package of *Upside* as well as the necessary parameter files are available to public access on GitHub: https://github.com/sosnicklab/upside-md.

**Table S1. Thermodynamic and kinetic parameters of GlpG folding**

| Method | Lipid environment | $\Delta G^{o}_{N\text{-}D}$ (kcal/mol) | $k_D$ ($min^{-1}$) |
|---|---|---|---|
| SDS denaturation (activity)[a] | DDM/SDS mixed micelles | 7.39 ± 0.20 (kinetic)<br>8.23 ± 1.43 (equilibrium) | $6.00 \cdot 10^{-6}$ |
| SDS denaturation (Trp fluorescence)[b] | DDM/SDS mixed micelles | 5.2 ± 0.8 | ND |
| Single-molecule force spectroscopy[c] | DMPC/CHAPSO bicelles | 3.9 | $3.38 \cdot 10^{-3}$ |
| Single-molecule force spectroscopy[d] | DMPC/DMPG/CHPASO bicelles | 9.1 | $4.3 \cdot 10^{-2}$ [e] |
| Steric trapping | DDM micelles | 5.8 ± 0.2 (N-subdomain)[f]<br>4.7 ± 0.1 (C-subdomain)[f] | $2.5 \cdot 10^{-4}$ [g]<br>$4.2 \cdot 10^{-3}$ [g] |
| Steric trapping[h] | DMPC/DMPG/CHAPS bicelles | ND | $1.8 \pm 0.1 \cdot 10^{-4}$<br>$2.7 \pm 0.7 \cdot 10^{-4}$ |

[a] Baker, RP and Urban, S (2012) Architectural and thermodynamic principles underlying intramembrane protease function. *Nat Chem Biol* 8, 759–768.

[b] Paslawski, W, Lillelund, OK, Kristensen, JV, Schafer, NP, Baker, RP, Urban, S, and Otzen, DE (2015) Cooperative folding of a polytopic α-helical membrane protein involves a compact N-terminal nucleus and nonnative loops. *PNAS* 112, 7978.

[c] Min, D, Jefferson, RE, Bowie, JU and Yoon, TY (2015) Mapping the energy landscape for second-stage folding of a single membrane protein. *Nat Chem Biol* 11, 981-987.

[d] Choi, H-K, Min, D., Kang, H., Shon, M., Rah, S-H, Kim, HC, Jeong, H, Choi, H-J, Bowie, JU and Yoon, T-Y (2019) Watching helical membrane proteins fold reveals a common N- to C-terminal folding pathway. *Science* 366, 1150–1156.

[e] Transition from N to $I_2$.

[f] Guo, R, Gaffney, KA, Kim, M, Yang, Z, Sungsuwan, S, Huang, X, Hubbell, WL and Hong, H (2016) Steric trapping reveals a cooperative network in the intramembrane protease GlpG. *Nat Chem. Biol.* 12, 353-360.

[g] Yang, Y, Guo, R, Gaffney, KA, Kim, M, Muhammednazaar, S, Wang, B, Wei, T, Liang, J, and Hong, H (2018) Folding-degradation relationship of a membrane protein mediated by the universally conserved ATP-dependent protease FtsH. *J. Am. Chem. Soc*. 140, 13, 4656-4665.

**Table S2. Identification of the proteolysis products of native GlpG ($172_{M}267_{C}$-$BtnRG_2$) in DDM micelles.** The peptides were obtained from ProK treatment and analyzed using CZE- or LC-tandem mass spectrometry. The mass spectra were analyzed using the Bionyc™ (Protein Metrics) program.

| Method | Query #:*z* | Peptide ID | Peptide | Start | End | Observed *m/z* | *z* | Observed (M+H) | Calc. mass (M+H) | Mass error (ppm) | **Score**[a] | \|Log Prob\|[c] | Scan # | Scan Time |
|---|---|---|---|---|---|---|---|---|---|---|---|---|---|---|
| CZE | 05874:1 | 2 | S.HMA.A | 85 | 87 | 358.154 | 1 | 358.154 | 358.154 | -0.5 | **565.1** | 8.0 | 7740 | 23.821 |
| CZE | 05412:2 | 3 | A.ALRER.A | 88 | 92 | 322.696 | 2 | 644.384 | 644.384 | -0.2 | **38.0** | 1.99 | 7190 | 22.1998 |
| CZE | 06225:1 | 4 | E.RAG.P | 92 | 94 | 303.178 | 1 | 303.178 | 303.178 | 0.0 | **36.6** | 2.32 | 8204 | 25.1352 |
| CZE | 04680:1 | 5 | F.THA.L | 140 | 143 | 328.161 | 1 | 328.161 | 328.162 | -0.7 | **78.3** | 2.26 | 6338 | 19.3901 |
| CZE | 09442:2 | 6 | F.THALMH.F | 140 | 145 | 355.182 | 2 | 709.358 | 709.345 | 17.7 | **2.1** | 1.20 | 11984 | 35.4047 |
| CZE | 06017:1 | 7 | T.HAL.M | 141 | 143 | 340.198 | 1 | 340.198 | 340.198 | -0.1 | **111.1** | 2.32 | 7933 | 24.3891 |
| CZE | 15168:1 | 9 | Y.LGGA.V | 161 | 164 | 317.182 | 1 | 317.182 | 317.182 | -0.4 | **112.0** | 6.45 | 18287 | 52.267 |
| CZE | 05385:2 | 10 | G.GAVEKR.L | 163 | 168 | 330.201 | 2 | 659.395 | 659.384 | 17.0 | **81.1** | 0.97 | 7159 | 22.1107 |
| CZE | 05528:2 | 12 | V.EKRLG.S | 166 | 170 | 301.685 | 2 | 602.362 | 602.362 | -0.4 | **84.6** | 2.54 | 7321 | 22.5918 |
| CZE | 06432:2 | 13 | G.PWFGGLSGV.V | 195 | 203 | 460.241 | 2 | 919.475 | 919.467 | 8.2 | **26.6** | 0.61 | 8467 | 25.8473 |
| CZE | 07475:2 | 15 | W.IVAGWF.D | 236 | 242 | 346.695 | 2 | 692.383 | 692.377 | 9.4 | **10.3** | 0.93 | 9758 | 29.4158 |
| CZE | 05922:2 | 16 | G.WFDLFG.M | 240 | 246 | 392.690 | 2 | 784.373 | 784.367 | 8.7 | **1.9** | 0.93 | 7804 | 24.0104 |
| CZE | 17109:1 | 17 | M.ANGA.H | 250 | 253 | 332.156 | 1 | 332.156 | 332.157 | -1.6 | **12.4** | 3.09 | 20422 | 58.0343 |
| CZE | 06159:1 | 18 | A.HIA.G | 254 | 256 | 340.198 | 1 | 340.198 | 340.198 | 0.2 | **269.2** | 4.32 | 8121 | 24.9135 |
| CZE | 05162:3 | 21 | F.C[+57.021]DSLNARKRK.- | 267 | 276 | 416.559 | 3 | 1247.664 | 1247.664 | 0.1 | **629.8** | 3.51 | 6900 | 21.2808 |
| LC | 00029:1 | 1 | R.GSHMA.A | 83 | 87 | 502.209 | 1 | 502.209 | 502.208 | 2.1 | **264.5** | 0.68 | 32 | 1.2104 |
| LC | 00383:1 | 8 | L.LNFLIH.M | 150 | 155 | 756.425 | 1 | 756.425 | 756.440 | -19.7 | **58.5** | 0.17 | 500 | 4.2178 |
| LC | 00275:2 | 11 | G.AVEKRL.C | 164 | 169 | 358.232 | 2 | 715.457 | 715.446 | 15.2 | **125.1** | 0.29 | 356 | 3.3033 |
| LC | 00324:3 | 14 | G.RLWVYGMLAYVVG.S | 202 | 214 | 509.620 | 3 | 1526.845 | 1526.819 | 17.3 | **24.8** | 0.20 | 421 | 3.7162 |
| LC | 00429:1 | 19 | A.GLAVGL.A | 257 | 262 | 529.335 | 1 | 529.335 | 529.334 | 1.0 | **319.7** | 0.68 | 561 | 4.6045 |
| LC | 00416:1 | 20 | A.GLAVGLA.M | 257 | 263 | 600.372 | 1 | 600.372 | 600.372 | -0.1 | **274.7** | 0.59 | 544 | 4.498 |

[a]**Score** – Byonic score, the primary indicator of PSM correctness. Byonic scores reflect the absolute quality of the peptide-spectrum match, not the relative quality compared to other candidate peptides. Byonic scores range from 0 to about 1000, with 300 a good score, 400 a very good score, and PSMs with scores over 500 almost sure to be correct.

[b]**Delta** – The drop in Byonic score from the top-scoring peptide to the next distinct peptide. In this computation, the same peptide with different modifications is not considered distinct.

[c]**Log Probability** – The log p-value of the PSM. This is the log of the probability that the PSM with such a score and delta would arise by chance in a search of this size (size of the protein database, as expanded by the modification rules). A log p-value of −3.0 should happen by chance on only one of a thousand spectra.

**Table S3. Identification of the proteolysis products of denatured GlpG ($172_M267_C$-BtnRG$_2$·mSA$_2$) in DDM micelles.** The peptides were obtained from ProK treatment and analyzed using CZE- or LC-tandem mass spectrometry. The mass spectra were analyzed using the Bionyc$^{TM}$ (Protein Metrics) program.

| Method | Query #:*z* | Peptide ID | Peptide | Start | End | Observed *m/z* | *z* | Observed (M+H) | Calc. mass (M+H) | Mass error (ppm) | Score | \|Log Prob\| | Scan # | Scan Time |
|---|---|---|---|---|---|---|---|---|---|---|---|---|---|---|
| CZE | 29819:1 | 1 | H.MAAL.R | 86 | 89 | 405.216 | 1 | 405.216 | 405.217 | -0.5 | 143.5 | 3.32 | 33931 | 90.538 |
| CZE | 09318:2 | 2 | L.RERAGPVTW.V | 90 | 98 | 536.289 | 2 | 1071.570 | 1071.569 | 0.7 | 16.8 | 2.97 | 11366 | 32.3082 |
| CZE | 08063:1 | 3 | E.RAG.P | 92 | 94 | 303.177 | 1 | 303.177 | 303.178 | -0.6 | 37.4 | 3.35 | 9963 | 28.4426 |
| CZE | 13014:1 | 4 | Y.FTH.A | 139 | 141 | 404.189 | 1 | 404.189 | 404.193 | -10.4 | 16.8 | 1.74 | 15435 | 42.8452 |
| CZE | 07839:2 | 5 | H.ALMHFS.L | 142 | 147 | 353.177 | 2 | 705.346 | 705.339 | 9.9 | 73.2 | 1.93 | 9703 | 27.7399 |
| CZE | 29246:1 | 6 | Y.LGGA.V | 161 | 164 | 317.182 | 1 | 317.182 | 317.182 | 0.3 | 413.26 | 21.66 | 33301 | 89.0271 |
| CZE | 07916:2 | 7 | G.GAVEKR.L | 163 | 168 | 330.196 | 2 | 659.384 | 659.384 | 0.2 | 53.1 | 3.73 | 9793 | 27.9874 |
| CZE | 06914:2 | 8 | G.AVEKR.L | 164 | 168 | 301.685 | 2 | 602.362 | 602.362 | 0.1 | 135.0 | 4.06 | 8605 | 24.652 |
| CZE | 08433:2 | 9 | G.PWFGGLSGV.V | 195 | 203 | 460.241 | 2 | 919.475 | 919.467 | 7.9 | 36.8 | 1.55 | 10385 | 29.6195 |
| CZE | 23580:2 | 11 | R.GERDPQSGIY.L | 215 | 224 | 561.265 | 2 | 1121.522 | 1121.522 | 0.0 | 1169.39 | 44.17 | 27067 | 73.0651 |
| CZE | 27434:1 | 15 | S.MAN.G | 249 | 251 | 335.138 | 1 | 335.138 | 335.138 | -0.8 | 1823.67 | 43.36 | 31307 | 83.8533 |
| CZE | 28809:1 | 17 | M.ANGA.H | 250 | 253 | 332.156 | 1 | 332.156 | 332.157 | -1.4 | 203.05 | 9.82 | 32819 | 87.7909 |
| CZE | 07471:1 | 19 | A.HIA.G | 254 | 256 | 340.198 | 1 | 340.198 | 340.198 | -0.1 | 1216.74 | 3.90 | 9267 | 26.5312 |
| CZE | 28853:1 | 20 | A.GLAV.G | 257 | 260 | 359.229 | 1 | 359.229 | 359.229 | -0.5 | 297.10 | 10.42 | 32868 | 87.9223 |
| CZE | 20077:2 | 23 | M.AFC[+57.021]DSLNAR.K | 265 | 273 | 527.243 | 2 | 1053.479 | 1053.478 | 0.5 | 2129.08 | 41.22 | 23209 | 63.1161 |
| CZE | 19498:2 | 24 | F.C[+57.021]DSLNAR.K | 267 | 273 | 418.190 | 2 | 835.373 | 835.373 | -0.1 | 289.2 | 5.32 | 22572 | 61.455 |
| CZE | 06580:3 | 25 | F.C[+57.021]DSLNARKRK.- | 267 | 276 | 416.559 | 3 | 1247.663 | 1247.664 | -0.7 | 617.59 | 9.60 | 8201 | 23.4656 |
| CZE | 11060:1 | 26 | L.NAR.K | 271 | 273 | 360.199 | 1 | 360.199 | 360.199 | -0.4 | 145.9 | 3.52 | 13284 | 37.3501 |
| LC | 00168:1 | 10 | V.VGSLGG.F | 198 | 203 | 489.267 | 1 | 489.267 | 489.267 | 0.2 | 255.5 | 255.5 | 0.23 | 197 |
| LC | 00659:3 | 12 | R.GERDPQSGIYLQRGLIIFALIWIVAGWFDLFGMS.M | 215 | 248 | 1290.352 | 3 | 3869.042 | 3869.025 | 4.5 | 370.4 | 2.65 | 828 | 6.3046 |
| LC | 00666:3 | 13 | R.GERDPQSGIYLQRGLIIFALIWIVAGWFDLFGMSM.A | 215 | 249 | 1334.026 | 3 | 4000.062 | 4000.065 | -0.8 | 280.7 | 0.86 | 837 | 6.3614 |
| LC | 00673:3 | 14 | Y.LQRGLIIFALIWIVAGWFDLFGMS.M | 225 | 248 | 922.845 | 3 | 2766.521 | 2766.520 | 0.1 | 800.2 | 8.46 | 847 | 6.4269 |
| LC | 00508:4 | 16 | S.MANGAHIAGLAVGLAMAFCDSLNARKRK.- | 249 | 276 | 722.382 | 4 | 2886.505 | 2886.501 | 1.2 | 1231.5 | 12.03 | 631 | 5.0323 |
| LC | 00497:4 | 17 | A.NGAHIAGLAVGLAMAFCDSLNARKRK.- | 251 | 276 | 671.862 | 4 | 2684.428 | 2684.424 | 1.6 | 612.4 | 6.42 | 616 | 4.9339 |
| LC | 00501:4 | 18 | M.ANGAHIAGLAVGLAMAFCDSLNARKRK.- | 250 | 276 | 689.622 | 4 | 2755.466 | 2755.461 | 1.9 | 462.0 | 4.85 | 621 | 4.9656 |
| LC | 00477:3 | 21 | A.GLAVGLAMAFCDSLNARKRK.- | 257 | 276 | 707.720 | 3 | 2121.146 | 2121.142 | 2.0 | 499.2 | 4.69 | 589 | 4.7595 |
| LC | 00262:1 | 22 | L.AVGLA.M | 259 | 263 | 430.266 | 1 | 430.266 | 430.266 | -0.3 | 901.41 | 0.67 | 315 | 3.0216 |

**Table S4. Identification of the proteolysis products of native GlpG ($172_M267_C$-BtnRG$_2$) in *E. coli* liposomes.** The peptides were obtained from ProK treatment and analyzed using CZE- or LC-tandem mass spectrometry. The mass spectra were analyzed using the Bionyc™ (Protein Metrics) program.

| Method | Query #:*z* | Peptide ID | Peptide | Start | End | Observed *m/z* | *z* | Observed (M+H) | Calc. mass (M+H) | Mass error (ppm) | Score | \|Log Prob\| | Scan # | Scan Time |
|---|---|---|---|---|---|---|---|---|---|---|---|---|---|---|
| CZE | 07785:2 | 1 | A.WPFDPTL.K | 125 | 131 | 438.222 | 2 | 875.437 | 875.430 | 8.5 | 22.8 | 0.78 | 10690 | 32.4472 |
| CZE | 12375:2 | 2 | H.ALMHFSL.M | 142 | 148 | 409.709 | 2 | 818.411 | 818.423 | -14.9 | 77.9 | 1.56 | 16964 | 51.2422 |
| CZE | 15246:2 | 3 | L.WWWYLGGAV.E | 157 | 165 | 569.286 | 2 | 1137.565 | 1137.552 | 11.7 | 3.0 | 0.45 | 20983 | 63.1773 |
| CZE | 21367:1 | 4 | L.GGAV.E | 162 | 165 | 303.166 | 1 | 303.166 | 303.166 | -0.6 | 190.6 | 3.16 | 29248 | 88.2458 |
| CZE | 20599:1 | 5 | V.YAL.M | 205 | 207 | 366.202 | 1 | 366.202 | 366.202 | 0.0 | 988.4 | 11.5 | 28222 | 85.1133 |
| LC | 00237:1 | 6 | G.LAVGL.A | 258 | 262 | 472.313 | 1 | 472.313 | 472.313 | 0.6 | 427.8 | 0.01 | 336 | 4.4918 |

**Table S5. Identification of the proteolysis products of denatured GlpG ($172_M267_C$-BtnRG$_2$·mSA$_2$) in *E. coli* liposomes.** The peptides were obtained from ProK treatment and analyzed using CZE- or LC-tandem mass spectrometry. The mass spectra were analyzed using the Bionyc$^{TM}$ (Protein Metrics) program.

| Method | Query #:*z* | Peptide ID | Peptide | Start | End | Observed *m/z* | *z* | Observed (M+H) | Calc. mass (M+H) | Mass error (ppm) | Score | \|Log Prob\| | Scan # | Scan Time |
|---|---|---|---|---|---|---|---|---|---|---|---|---|---|---|
| CZE | 07554:1 | 1 | S.HMA.A | 85 | 87 | 358.154 | 1 | 358.154 | 358.154 | -0.1 | 655.3 | 11.45 | 9877 | 29.5011 |
| CZE | 09939:1 | 2 | S.HMAAL.R | 85 | 89 | 542.277 | 1 | 542.277 | 542.276 | 1.9 | 175.3 | 2.29 | 13352 | 39.6613 |
| CZE | 11899:2 | 4 | C.VTWVMM.I | 96 | 101 | 383.690 | 2 | 766.373 | 766.363 | 13.6 | 42.1 | 0.33 | 16248 | 48.0877 |
| CZE | 19851:1 | 5 | M.LW.L | 121 | 122 | 318.181 | 1 | 318.181 | 318.181 | -0.5 | 55.6 | 1.26 | 27922 | 83.0845 |
| CZE | 21574:1 | 6 | F.EFW.R | 134 | 136 | 481.205 | 1 | 481.205 | 481.208 | -7.3 | 21.0 | 0.14 | 30479 | 90.7104 |
| CZE | 08476:2 | 7 | A.VEKRLG.S | 165 | 170 | 351.219 | 2 | 701.431 | 701.431 | 0.1 | 9.2 | 1.72 | 11213 | 33.4203 |
| CZE | 20119:1 | 8 | S.ALL.S | 182 | 184 | 316.223 | 1 | 316.223 | 316.223 | -0.3 | 422.8 | 21.40 | 28324 | 84.3086 |
| CZE | 21491:1 | 9 | L.SGY.V | 185 | 187 | 326.135 | 1 | 326.135 | 326.135 | 0.3 | 127.1 | 2.36 | 30387 | 90.5406 |
| CZE | 21115:1 | 10 | F.GGLSG.V | 197 | 202 | 390.199 | 1 | 390.199 | 390.198 | 0.5 | 17.9 | 1.32 | 29840 | 88.9012 |
| CZE | 19971:1 | 11 | V.YAL.M | 205 | 207 | 366.202 | 1 | 366.202 | 366.202 | -0.2 | 478.6 | 11.23 | 28104 | 83.6354 |
| CZE | 17126:2 | 12 | E.RDPQSGIY.L | 217 | 224 | 468.233 | 2 | 935.460 | 935.458 | 1.6 | 143.3 | 2.26 | 4010 | 71.2103 |
| CZE | 07559:1 | 14 | A.HIA.G | 254 | 256 | 340.198 | 1 | 340.198 | 340.198 | -0.3 | 262.8 | 4.38 | 9883 | 29.5118 |
| CZE | 06912:2 | 15 | A.HIAGLAV.G | 254 | 260 | 340.708 | 2 | 680.409 | 680.409 | 0.0 | 17.8 | 1.59 | 8911 | 26.5789 |
| CZE | 09004:1 | 22 | L.NAR.K | 271 | 273 | 360.199 | 1 | 360.199 | 360.199 | -0.8 | 104.1 | 4.40 | 11969 | 35.6212 |
| LC | 00193:1 | 3 | H.MAAL.R | 85 | 89 | 405.217 | 1 | 405.217 | 405.217 | -0.4 | 255.8 | 0.06 | 290 | 3.872 |
| LC | 00061:1 | 13 | N.GAHIA.G | 252 | 256 | 468.258 | 1 | 468.258 | 468.257 | 2.1 | 236.6 | 0.39 | 71 | 1.6147 |
| LC | 00262:1 | 16 | A.GLAVGL.A | 257 | 262 | 529.335 | 1 | 529.335 | 529.334 | 1.2 | 343.0 | 0.85 | 378 | 4.6213 |
| LC | 00282:3 | 17 | A.GLAVGLAMAFC[+57.021]DSLNARKRK.- | 257 | 276 | 726.727 | 3 | 2178.167 | 2178.163 | 1.4 | 364.1 | 2.07 | 405 | 4.8559 |
| LC | 00258:3 | 18 | L.AVGLAMAFC[+57.021]DSLNARKRK.- | 259 | 276 | 670.025 | 3 | 2008.059 | 2008.058 | 0.5 | 365.3 | 1.62 | 373 | 4.5816 |
| LC | 00213:3 | 19 | L.AMAFC[+57.021]DSLNARKRK.- | 263 | 276 | 556.621 | 3 | 1667.848 | 1667.847 | 0.8 | 775.7 | 4.40 | 317 | 4.1045 |
| LC | 00217:2 | 20 | M.AFC[+57.021]DSLNAR.K | 265 | 273 | 527.244 | 2 | 1053.480 | 1053.478 | 1.5 | 453.9 | 0.35 | 322 | 4.1437 |
| LC | 00199:3 | 21 | M.AFC[+57.021]DSLNARKRK.- | 265 | 276 | 489.262 | 3 | 1465.770 | 1465.769 | 0.7 | 290.3 | 1.35 | 298 | 3.9398 |

**Table S6. Identification of the proteolysis products of native GlpG ($95_N172_M$-BtnRG$_2$) in DDM micelles.** The peptides were obtained from ProK treatment and analyzed using LC-tandem mass spectrometry. The mass spectra were analyzed using the Bionyc™ (Protein Metrics) program.

| Method | Query #:*z* | Peptide ID | Peptide | Start | End | Observed *m/z* | *z* | Observed (M+H) | Calc. mass (M+H) | Mass error (ppm) | Score | \|Log Prob\| | Scan # | Scan Time |
|---|---|---|---|---|---|---|---|---|---|---|---|---|---|---|
| LC | 00034:1 | 1 | R.GSHMA.A | 83 | 87 | 502.208 | 1 | 502.208 | 502.208 | 0.4 | 249.3 | 0.03 | 27 | 1.2445 |
| LC | 00029:1 | 2 | S.HMA.A | 85 | 87 | 358.154 | 1 | 358.154 | 358.154 | -0.2 | 179.3 | 0.00 | 24 | 1.2161 |
| LC | 00047:1 | 3 | M.AALR.E | 87 | 90 | 430.277 | 1 | 430.277 | 430.277 | -0.8 | 158.6 | 0.00 | 40 | 1.3506 |
| LC | 00335:1 | 4 | C.VTWV.M | 96 | 99 | 504.285 | 1 | 504.285 | 504.282 | 7.1 | 122.4 | 0.00 | 396 | 4.3618 |
| LC | 00182:1 | 5 | Q.ILGD.Q | 113 | 116 | 417.234 | 1 | 417.234 | 417.234 | -0.4 | 37.9 | 0.00 | 200 | 2.7036 |
| LC | 00252:1 | 6 | D.QEVM.L | 117 | 120 | 506.224 | 1 | 506.224 | 506.228 | -6.9 | 14.4 | 0.00 | 289 | 3.4576 |
| LC | 00282:1 | 7 | L.KFE.F | 132 | 134 | 423.223 | 1 | 423.223 | 423.224 | -1.1 | 3.4 | 0.00 | 329 | 3.7936 |
| LC | 00227:1 | 8 | M.HIL.F | 150 | 152 | 382.244 | 1 | 382.244 | 382.245 | -1.5 | 140.4 | 0.00 | 260 | 3.2168 |
| LC | 00221:1 | 9 | G.KLI.V | 173 | 175 | 373.281 | 1 | 373.281 | 373.281 | -0.7 | 216.6 | 0.00 | 252 | 3.1488 |
| LC | 00030:1 | 10 | G.LSGV.V | 184 | 187 | 375.235 | 1 | 375.235 | 375.224 | 29.6 | 29.9 | 0.00 | 25 | 1.2213 |
| LC | 00260:1 | 11 | V.VYA.L | 204 | 207 | 352.186 | 1 | 352.186 | 352.187 | -1.5 | 27.4 | 0.00 | 300 | 3.5534 |
| LC | 00050:1 | 12 | Y.LQR.G | 225 | 227 | 416.261 | 1 | 416.261 | 416.262 | -0.7 | 66.4 | 0.00 | 44 | 1.3842 |
| LC | 00162:1 | 13 | V.AGW.F | 239 | 241 | 333.156 | 1 | 333.156 | 333.156 | -0.3 | 68.6 | 0.00 | 173 | 2.4703 |
| LC | 00127:2 | 14 | G.WFDLFGM.S | 241 | 247 | 458.211 | 2 | 915.415 | 915.407 | 8.4 | 62.6 | 0.00 | 129 | 2.0988 |
| LC | 00336:1 | 15 | F.DLFG.M | 243 | 246 | 451.219 | 1 | 451.219 | 451.219 | -0.4 | 39.2 | 0.00 | 397 | 4.367 |
| LC | 00110:1 | 16 | H.IAGLA.V | 255 | 259 | 444.282 | 1 | 444.282 | 444.282 | -0.2 | 43.4 | 0.00 | 112 | 1.9586 |
| LC | 00146:1 | 17 | A.GLAVG.L | 257 | 261 | 416.250 | 1 | 416.250 | 416.250 | -1.0 | 97.3 | 0.00 | 152 | 2.2958 |
| LC | 00347:1 | 18 | A.GLAVGLA.M | 257 | 263 | 600.371 | 1 | 600.371 | 600.372 | -0.2 | 265.5 | 0.01 | 412 | 4.4965 |
| LC | 00179:1 | 19 | G.LAMA.F | 262 | 265 | 405.216 | 1 | 405.216 | 405.217 | -0.5 | 114.3 | 0.00 | 196 | 2.6692 |
| LC | 00231:1 | 20 | M.AFVD.S | 265 | 268 | 451.219 | 1 | 451.219 | 451.219 | -0.6 | 124.0 | 0.00 | 265 | 3.2558 |

**Table S7. Identification of the proteolysis products of denatured GlpG ($95_N172_M$-BtnRG$_2$·mSA$_2$) in DDM micelles.** The peptides were obtained from ProK treatment and analyzed using LC-tandem mass spectrometry. The mass spectra were analyzed using the Bionyc$^{TM}$ (Protein Metrics) program.

| Method | Query #:*z* | Peptide ID | Peptide | Start | End | Observed *m/z* | *z* | Observed (M+H) | Calc. mass (M+H) | Mass error (ppm) | Score | \|Log Prob\| | Scan # | Scan Time |
|---|---|---|---|---|---|---|---|---|---|---|---|---|---|---|
| LC | 00023:1 | 1 | S.HMA.A | 86 | 87 | 358.154 | 1 | 358.154 | 358.154 | -0.4 | 93.4 | 1.55 | 24 | 1.2159 |
| LC | 00326:1 | 2 | M.IACVV.V | 101 | 107 | 504.285 | 1 | 504.285 | 504.285 | -0.2 | 89.3 | 1.84 | 400 | 4.3926 |
| LC | 00196:1 | 3 | F.IAM.Q | 109 | 111 | 334.179 | 1 | 334.179 | 334.180 | -0.8 | 217.0 | 2.14 | 231 | 2.9703 |
| LC | 00106:1 | 4 | M.HIL.F | 150 | 153 | 382.245 | 1 | 382.245 | 382.245 | -1.1 | 194.9 | 3.90 | 131 | 2.1218 |
| LC | 00313:2 | 5 | W.YLGGAVEKR.L | 159 | 168 | 496.775 | 2 | 992.542 | 992.552 | -10.3 | 1.7 | 0.77 | 387 | 4.286 |
| LC | 00024:1 | 6 | G.AVEK.R | 164 | 167 | 446.261 | 1 | 446.261 | 446.261 | 0.3 | 111.9 | 1.67 | 25 | 1.2211 |
| LC | 00309:1 | 7 | A.VEKRL.C | 165 | 169 | 644.397 | 1 | 644.397 | 644.409 | -18.0 | 77.4 | 0.59 | 381 | 4.2291 |
| LC | 00201:1 | 8 | G.KLI.V | 173 | 175 | 373.281 | 1 | 373.281 | 373.281 | -1.1 | 121.4 | 1.59 | 237 | 3.0146 |
| LC | 00316:1 | 9 | G.PWFGGL.S | 195 | 200 | 676.351 | 1 | 676.351 | 676.345 | 8.3 | 86.4 | 3.34 | 391 | 4.3199 |
| LC | 00502:7 | 10 | R.GERDPQSGIYLQRGLIIFALIWIVAGWFDLFG MSMANGAHIAGLAVGLAMAFVDSLNARKRK.- | 215 | 276 | 962.516 | 7 | 6731.568 | 6732.567 | 0.5 | 474.8 | 1.74 | 631 | 6.3651 |
| LC | 00505:5 | 11 | Y.LQRGLIIFALIWIVAGWFDLFGMSMANGAHI AGLAVGLAMAFVDSLNARKRK.- | 225 | 276 | 1126.820 | 5 | 5630.072 | 5630.063 | 1.6 | 2325.6 | 41.21 | 635 | 6.3989 |
| LC | 00341:1 | 12 | R.GLII.F | 228 | 231 | 415.291 | 1 | 415.291 | 415.292 | -1.5 | 127.6 | 1.59 | 420 | 4.5616 |
| LC | 00192:1 | 13 | R.GLI.I | 228 | 230 | 302.207 | 1 | 302.207 | 302.207 | -1.0 | 89.3 | 2.14 | 225 | 2.9127 |
| LC | 00103:2 | 14 | W.IVAGWF.D | 237 | 242 | 346.695 | 2 | 692.383 | 692.377 | 9.5 | 74.7 | 1.67 | 127 | 2.0881 |
| LC | 00393:4 | 15 | S.MANGAHIAGLAVGLAMAFVDSLNARKRK.- | 249 | 276 | 721.397 | 4 | 2882.565 | 2882.560 | 1.7 | 764.3 | 6.21 | 485 | 5.1097 |
| LC | 00427:3 | 16 | S.MANGAHIAGLAVGLAMAFVDSLNAR.K | 249 | 273 | 824.096 | 3 | 2470.272 | 2470.269 | 1.0 | 720.7 | 7.78 | 531 | 5.5087 |
| LC | 00042:1 | 17 | N.GAHIA.G | 252 | 256 | 468.256 | 1 | 468.256 | 468.257 | -0.3 | 160.6 | 2.04 | 45 | 1.3892 |
| LC | 00248:1 | 18 | A.HIAGL.A | 254 | 258 | 510.303 | 1 | 510.303 | 510.304 | -0.6 | 218.1 | 2.68 | 300 | 3.5507 |
| LC | 00319:1 | 19 | G.LAVGL.A | 258 | 262 | 472.313 | 1 | 472.313 | 472.313 | -0.5 | 16.7 | 1.84 | 395 | 4.3535 |
| LC | 00195:1 | 20 | L.AVGLA.M | 259 | 263 | 430.266 | 1 | 430.266 | 430.266 | -0.8 | 1309.6 | 15.84 | 229 | 2.9468 |
| LC | 00263:1 | 21 | L.AVGLAM.A | 259 | 264 | 561.303 | 1 | 561.303 | 561.307 | -6.7 | 20.4 | 1.67 | 320 | 3.7186 |
| LC | 00203:1 | 22 | M.AFVD.S | 264 | 268 | 451.219 | 1 | 451.219 | 451.219 | -0.4 | 254.1 | 2.73 | 240 | 3.0435 |

**Table S8. Fitted parameters of DEER data for native and sterically denatured GlpG in bicelles and liposomes.** The background-substracted dipolar evolution data were fitted to non-negative Tikhonov regularization.

Bicelles (DMPC:DMPG:CHAPS)

| | Max Peak (Å) | Median ± 0.5·SD (Å) | $\chi^2$ | Modulation depth | Distance limit (Å) | Shape limit (Å) | Max Time (μsec) |
|---|---|---|---|---|---|---|---|
| $95_N172_M$ | 28.2 | 29.0 ± 9.5 | 1.46 | 0.35 | 58 | 46 | 3.1 |
| $95_N172_M$·mSA$_2$ | 41.1 | 38.6 ± 13.8 | 1.25 | 0.40 | 58 | 46 | 3.1 |
| $172_M267_C$ | 29.5 | 30.6 ± 6.1 | 1.22 | 0.51 | 58 | 46 | 3.1 |
| $172_M267_C$·mSA$_2$ | 42.3 | 45.0 ± 15.3 | 1.24 | 0.29 | 60 | 48 | 3.4 |
| $95_N267_C$ | 42.8 | 38.2 ± 9.7 | 1.06 | 0.40 | 59 | 48 | 3.4 |
| $95_N267_C$·mSA$_2$ | 60.7 | 48.4 ± 19.2 | 1.72 | 0.18 | 62 | 50 | 3.9 |

Liposomes (*E. coli* phospholipids)

| | Max Peak (Å) | Median ± 0.5·SD (Å) | $\chi^2$ | Modulation depth | Distance limit (Å) | Shape limit (Å) | Max Time (μsec) |
|---|---|---|---|---|---|---|---|
| $95_N172_M$ | 30.0 | 29.1 ± 7.7 | 1.12 | 0.50 | 49 | 39 | 1.9 |
| $95_N172_M$·mSA$_2$ | 56.0 | 40.7 ± 20.1 | 1.08 | 0.32 | 60 | 48 | 3.4 |
| $172_M267_C$ | 24.4 | 26.0 ± 8.3 | 1.08 | 0.22 | 60 | 48 | 3.4 |
| $172_M267_C$· mSA$_2$ | 54.3 | 52.7 ± 26.6 | 1.08 | 0.25 | 60 | 48 | 3.5 |
| $95_N267_C$ | 40.6 | 35.8 ± 9.7 | 1.08 | 0.42 | 59 | 47 | 3.3 |
| $95_N267_C$·mSA$_2$ | 55.2 | 56.1 ± 8.1 | 1.01 | 0.25 | 58 | 46 | 3.1 |

**Table S9. Fitted parameters of DEER data for native and sterically denatured GlpG in bicelles and liposomes.** The background-substracted dipolar evolution data were fitted to the single- or double-Gaussian model.

Bicelles (DMPC/DMPG/CHAPS)

| | | Max peak (Å) | Width (Å) | Area | $\chi^2$ | Modulation depth | Distance limit (Å) | Shape limit (Å) | Max time (μsec) |
|---|---|---|---|---|---|---|---|---|---|
| $95_N172_M$ | Double | 26.0 ± 0.3<br>47.4 ± 1.9 | 12.7 ± 0.9<br>25.9 ± 2.8 | 0.72 ± 0.04<br>0.29 ± 0.03 | 1.47 | 0.34 | 56 | 45 | 2.79 |
| $95_N172_M·mSA_2$ | Single | 33.9 ± 1.5 | 47.1 ± 2.2 | 1.01 ± 0.02 | 1.51 | 0.41 | 58 | 46 | 3.11 |
| $172_M267_C$ | Double | 29.9 ± 0.7<br>47 ± 0 | 11.9 ± 2.2<br>33.2 ± 6.6 | 0.71 ± 0.05<br>0.22 ± 0.03 | 1.40 | 0.50 | 57 | 46 | 2.97 |
| $172_M267_C·$ $mSA_2$ | Single | 45.5 ± 0.8 | 32.5 ± 1.6 | 0.96 ± 0.02 | 1.41 | 0.26 | 57 | 46 | 2.99 |
| $95_N267_C$ | Single | 36.8 ± 0.2 | 22.0 ± 0.5 | 1.00 ± 0.01 | 1.77 | 0.37 | 57 | 46 | 2.99 |
| $95_N267_C·mSA_2$ | Single | 49.7 ± 0.3 | 31.9 ± 0.6 | 0.91 ± 0.01 | 3.51 | 0.15 | 59 | 47 | 3.25 |

Liposomes (*E. coli* lipids)

| | | Max peak (Å) | Width (Å) | Area | $\chi^2$ | Modulation depth | Distance limit (Å) | Shape limit (Å) | Max time (μsec) |
|---|---|---|---|---|---|---|---|---|---|
| $95_N172_M$ | Double | 26.7 ± 0.5<br>42.2 ± 4.0 | 10.1 ± 1.5<br>21.0 ± 5.5 | 0.68 ± 0.11<br>0.32 ± 0.1 | 1.24 | 0.50 | 49 | 39 | 1.9 |
| $95_N172_M·mSA_2$ | Single | 43.7 ± 0.9 | 43.2 ± 1.6 | 1.0 ± 0.02 | 1.82 | 0.36 | 59 | 47 | 3.2 |
| $172_M267_C$ | Double | 23.8 ± 0.6<br>49.4 ± 2.4 | 10.7 ± 1.9<br>23.2 ± 4.8 | 0.73 ± 0.06<br>0.27 ± 0.04 | 1.18 | 0.22 | 60 | 48 | 3.4 |
| $172_M267_C·mSA_2$ | Single | 52.4 ± 0.6 | 27.8 ± 1.4 | 0.91 ± 0.02 | 1.43 | 0.24 | 58 | 47 | 3.2 |
| $95_N267_C$ | Single | 33.9 ± 1.8 | 29.5 ± 3.5 | 0.98 ± 0.05 | 1.33 | 0.41 | 59 | 47 | 3.3 |
| $95_N267_C·mSA_2$ | Single | 54.6 ± 0.4 | 21.2 ± 1.1 | 0.97 ± 0.02 | 1.35 | 0.21 | 59 | 47 | 3.3 |

**Table S10. DSE simulations of GlpG alone and the three constructs doubly bound with mSA.** The temperature dependence of the $C_\alpha$-RMSD and $R_g$ are referenced to native GlpG (the mSA are not included). Values are the mean of 20 independent trajectories where errors denote their standard deviation ($n$ = 20). The RMSDs were calculated for all TM helices (TM1–TM6) or the TM helices excluding intrinsically flexible TM5 (in parentheses). Simulations of the "Expanded DSE" were performed at 240 K but with only repulsive side chain interactions. All values are in Å.

| | $T$ | GlpG | $95_N172_M\cdot mSA_2$ | $172_M267_C\cdot mSA_2$ | $95_N267_C\cdot mSA_2$ |
|---|---|---|---|---|---|
| $C_\alpha$-RMSD: all TMHs (TM 1–4 and 6) | 274 K | 5.2 ± 0.9 (3.0 ± 0.6) | 8.9 ± 0.9 (7.2 ± 0.7) | 8.7 ± 1.1 (5.4 ± 0.7) | 7.3 ± 1.2 (5.7 ± 1.1) |
| | 308 K | 9.0 ± 0.8 (5.7 ± 1.2) | 11.8 ± 1.6 (9.4 ± 1.2) | 12.2 ± 2.0 (7.7 ± 1.4) | 10.3 ± 0.9 (7.9 ± 0.9) |
| | 343 K | 14.4 ± 2.4 (9.9 ± 2.0) | 15.3 ± 2.5 (12.0 ± 2.7) | 17.4 ± 3.4 (11.9 ± 2.8) | 15.5 ± 2.5 (10.7 ± 2.3) |
| | 377 K | 26.2 ± 6.0 (22.0 ± 6.1) | 31.7 ± 3.8 (28.5 ± 3.2) | 37.4 ± 7.9 (30.4 ± 7.7) | 27.3 ± 4.8 (22.6 ± 3.7) |
| | 411 K | 39.4 ± 3.6 (34.3 ± 3.8) | 38.5 ± 4.3 (33.0 ± 3.2) | 42.1 ± 6.4 (37.6 ± 5.8) | 37.3 ± 3.5 (32.3 ± 3.8) |
| | Expanded DSE | 22.8 ± 1.4 (18.8 ± 1.4) | 23.6 ± 1.4 (21.2 ± 1.1) | 26.6 ± 2.9 (20.1 ± 1.8) | 22.4 ± 1.6 (18.0 ± 1.6) |
| $R_g$ | 274 K | 16.1 ± 0.3 | 17.2 ± 0.4 | 17.1 ± 0.5 | 17.1 ± 0.3 |
| | 308 K | 16.8 ± 0.3 | 18.0 ± 0.8 | 18.5 ± 1.2 | 18.0 ± 0.4 |
| | 343 K | 19.9 ± 1.6 | 21.1 ± 2.2 | 23.5 ± 3.4 | 22.8 ± 2.6 |
| | 377 K | 32.5 ± 7.1 | 39.4 ± 4.7 | 44.7 ± 7.9 | 35.8 ± 4.7 |
| | 411 K | 47.9 ± 4.2 | 47.3 ± 4.9 | 50.8 ± 6.8 | 45.6 ± 3.2 |
| | Expanded DSE | 30.1 ± 1.9 | 31.7 ± 1.6 | 34.6 ± 3.7 | 30.5 ± 1.8 |
| $C_\alpha$-$C_\alpha$ distance: (without bound mSA) | 274 K | | 31.6 ± 2.6 (10.2 ± 1.5) | 29.0 ± 3.2 (14.4 ± 1.1) | 35.8 ± 3.0 (5.7 ± 1.1) |
| | 308 K | | 34.9 ± 3.8 (13.3 ± 4.5) | 33.4 ± 4.2 (16.5 ± 3.5) | 39.9 ± 4.0 (23.3 ± 4.2) |
| | 343 K | | 38.8 ± 5.8 (14.1 ± 4.1) | 47.3 ± 11.8 (40.2 ± 21.5) | 52.2 ± 10.4 (44.3 ± 20.3) |
| | 377 K | | 66.8 ± 12.2 (41.1 ± 23.9) | 70.3 ± 17.2 (74.8 ± 16.4) | 77.3 ± 14.3 (80.3 ± 22.1) |
| | 411 K | | 69.9 ± 11.9 (71.9 ± 22.0) | 77.9 ± 13.2 (95.2 ± 16.7) | 83.5 ± 16.0 (106.7 ± 21.4) |
| | Expanded DSE | | 61.1 ± 6.9 (43.2 ± 11.9) | 55.9 ± 5.3 (55.3 ± 10.8) | 71.7 ± 10.2 (78.0 ± 17.9) |

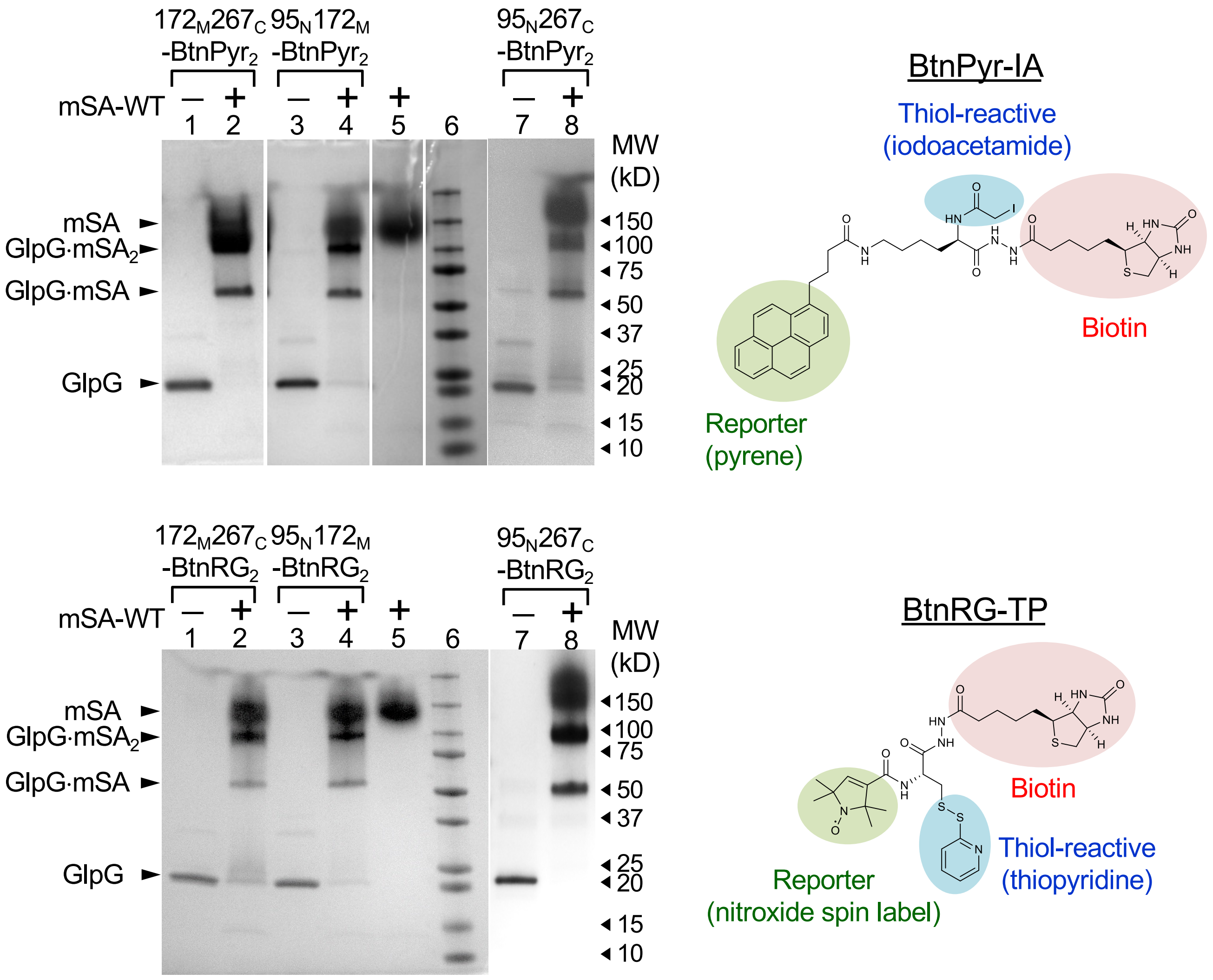


**Fig. S1. Labeling of double cysteine variants of GlpG ($95_N172_M$, $172_M267_C$ and $95_N267_C$) with the thiol-reactive biotin derivatives measured by SDS-PAGE.**

The thiol-reactive biotin derivative contains either fluorescent pyrenyl group (BtnPyr-IA, *top*) or paramagnetic spin label (BtnRG-TP, *bottom*). This assay utilizes the strong noncovalent bond between the biotinyl group and wild-type monovalent streptavidin (mSA-WT), which is resistant to a high concentration of SDS when the samples are not heated. In the presence of excess mSA-WT (lanes 2, 4 and 8), a major portion of double biotin variants of GlpG migrates as GlpG bound with two mSA molecules (GlpG·$mSA_2$) with minor portions of singly bound (GlpG·mSA) and unbound (GlpG) forms. mSA-WT (52 kD, lane 5) migrates as a higher molecular species (~150 kD) probably because the number of bound SDS molecules per a folded tetrameric mSA molecule is not proportional to the number of the total residues in mSA.

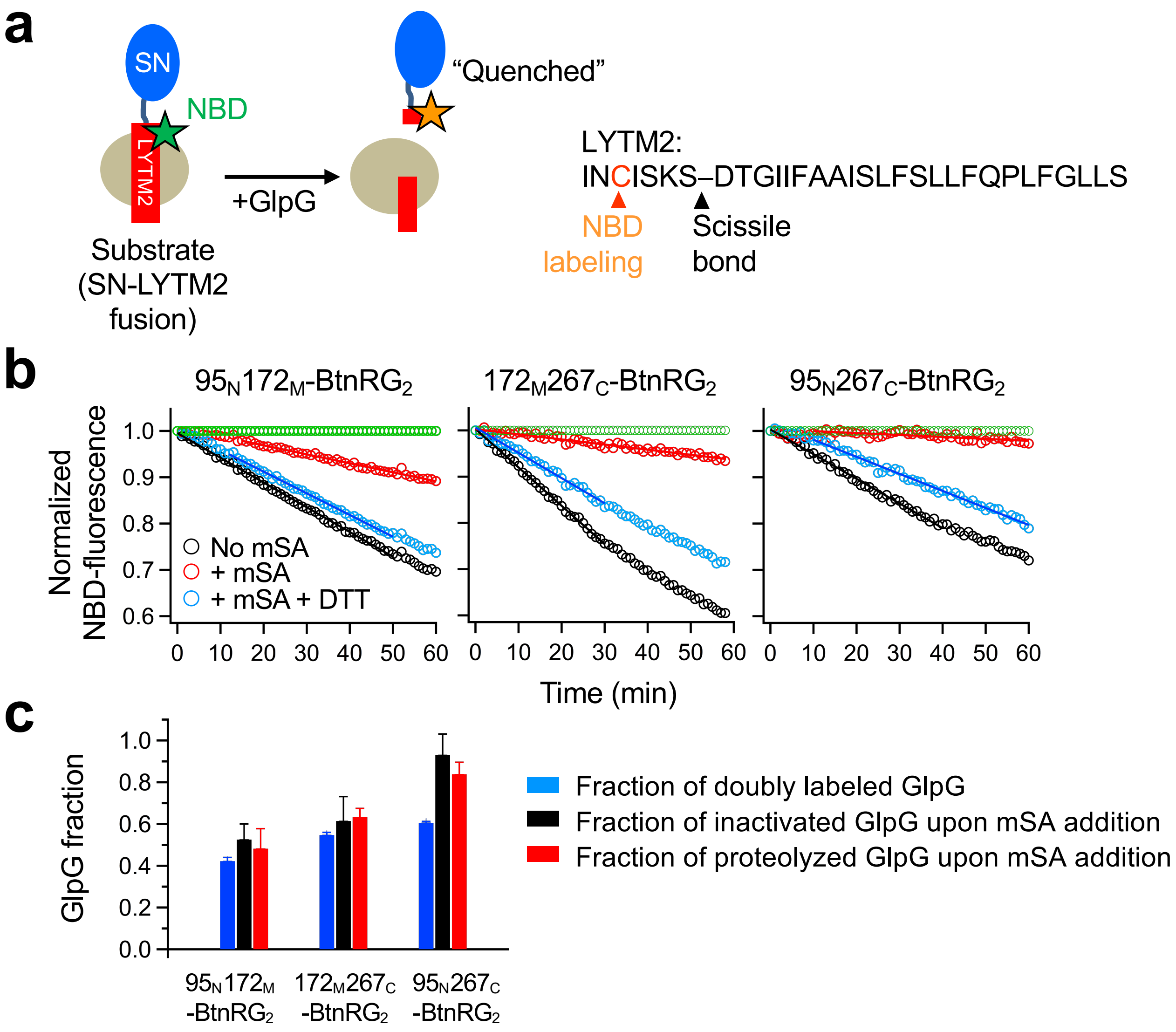


**Fig. S2. The correlation between the labeling efficiency, resistance to proteolysis by ProK and inactivation upon denaturation by steric trapping.**

**(a)** Schematic illustration of the activity assay of GlpG in micelles and bicelles. The model transmembrane substrate of GlpG (LYTM2) was fused to the stable globular protein staphylococcal nuclease (SN). The five-residue upstream of the scissile bond was substituted with cysteine, which was labeled with the environment-sensitive fluorophore NBD. Upon cleavage by GlpG, the NBD label is transferred from the hydrophobic environment to the aqueous phase, which induces quenching of NBD fluorescence. The initial slope of fluorescence decrease reflects activity of GlpG.

**(b)** Denaturation of GlpG (1 μM) doubly biotinylated with BtnRG (**Fig. S1**) upon addition of mSA-WT (10 μM) in DDM micelles. Addition of the reducing agent DTT, which breaks the disulfide linkage between GlpG and the biotin label bound with mSA, induces the regain of activity, indicating refolding of GlpG.

**(c)** The efficiency of double biotinylation of GlpG is correlated with the degrees of inactivation and proteolysis by ProK upon addition of mSA. The fraction of doubly biotinylated GlpG was obtained by quantifying the band intensities on SDS-PAGE (**Fig. S1**). The fractions of inactivated and proteolyzed GlpG were obtained by averaging those values obtained in micelles, bicelles and liposomes (**Figs. 2c** and **2d**, respectively). The error bars in doubly-labeling efficiency denote ± s.e.m. over multiple measurements (n = 3) while the error bars in inactivated and proteolyzed fractions indicate ± s.e.m. of the values obtained in micelles, bicelles and liposomes.

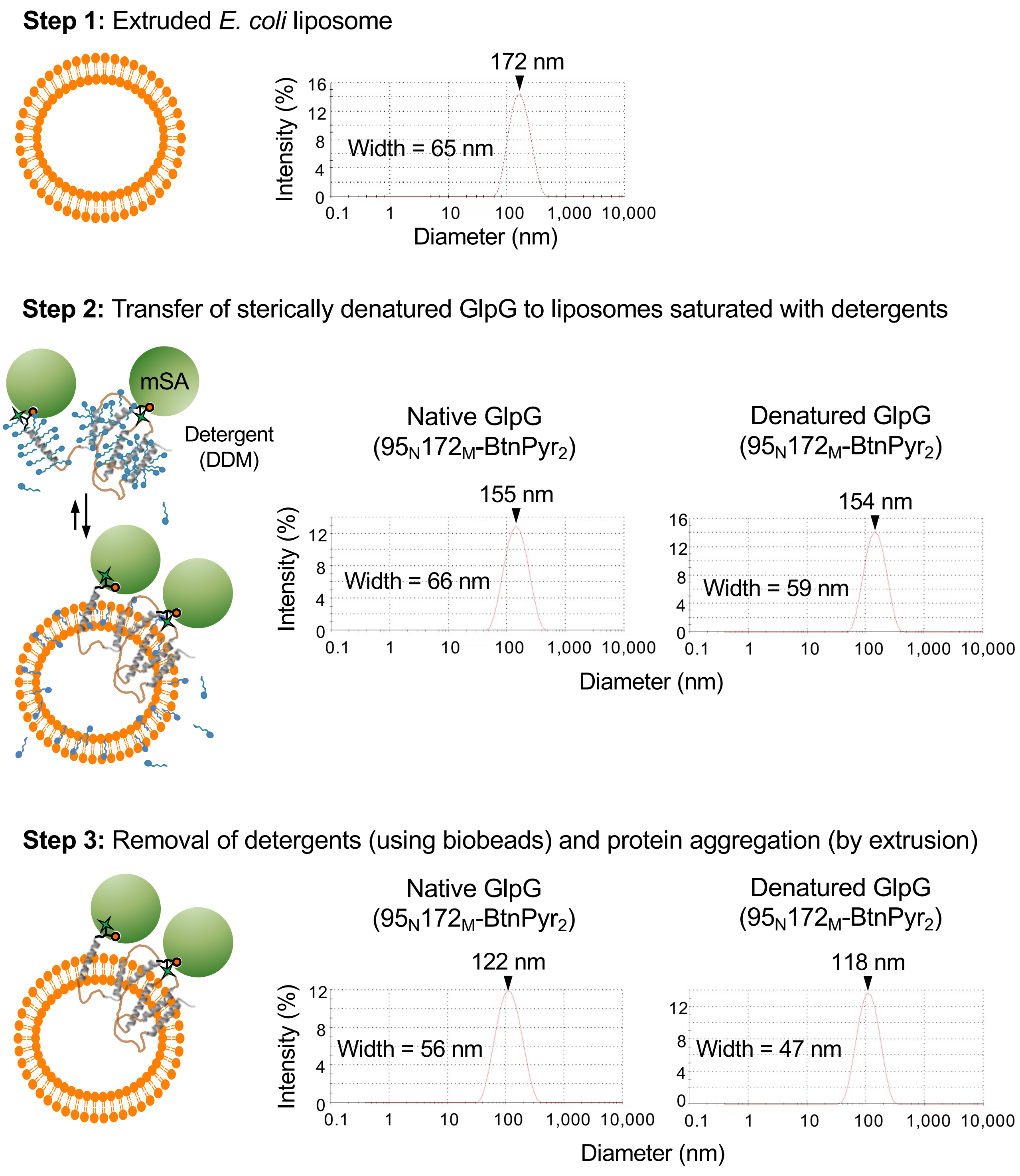


**Fig. S3. Size distributions of the proteoliposomes during the reconstitution steps for native and sterically denatured GlpG measured by dynamic light scattering.** This result indicates that the bilayer structure and size distribution of the liposomes are maintained during reconstitution. Detergent DDM was added to induce defects on liposomes that would facilitate incorporation of GlpG (**Step 2**).

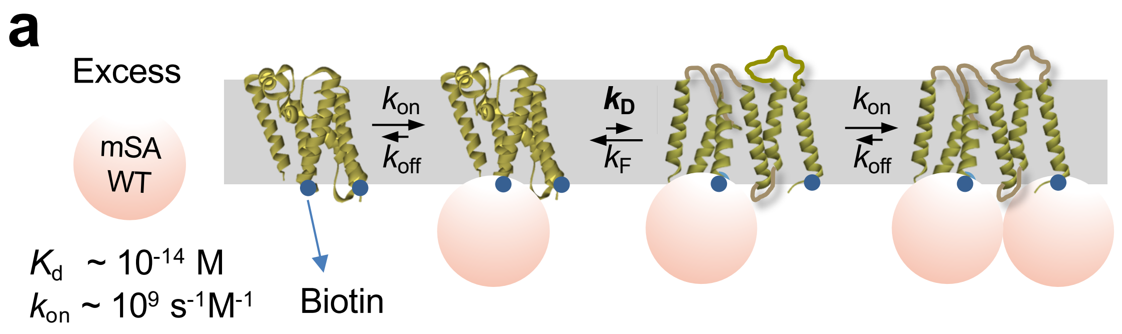


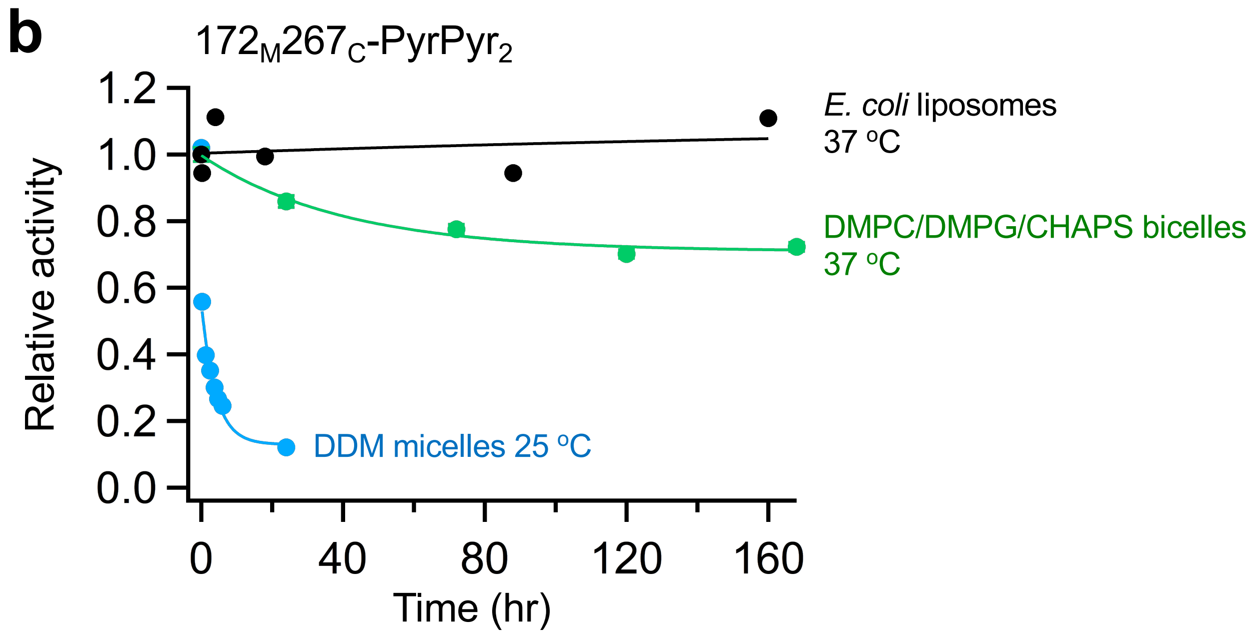


**Fig. S4. Denaturation kinetics of GlpG measured by steric trapping in various lipid environments.**

**(a)** The reaction scheme to measure the denaturation rate ($k_D$) of doubly biotinylated GlpG using steric trapping. When an excess concentration of mSA-WT with a high biotin affinity is added, the on-rate ($k_{on}$) of mSA binding to either biotin label is much faster than the off-rate ($k_{off}$). The denaturation rate of GlpG ($k_D$) is determined by monitoring the activity of GlpG as a function of time because binding of the second mSA binding ($k_{on}$[mSA] ~ $10^4$ $s^{-1}$) is much faster than denaturation of GlpG ($k_D$) under the condition where the folding rate ($k_F$ ~ $10^{-1}$ $s^{-1}$) is slower than the on-rate ($k_{on}$[mSA] ~ $10^4$ $s^{-1}$). That is, the denaturation of GlpG is the rate-determining step in the overall reaction.

**(b)** Denaturation kinetics of doubly biotinylated GlpG ($172_M267_C$-BtnPyr$_2$) measured by steric trapping in micelles, bicelles and liposomes.

**a**

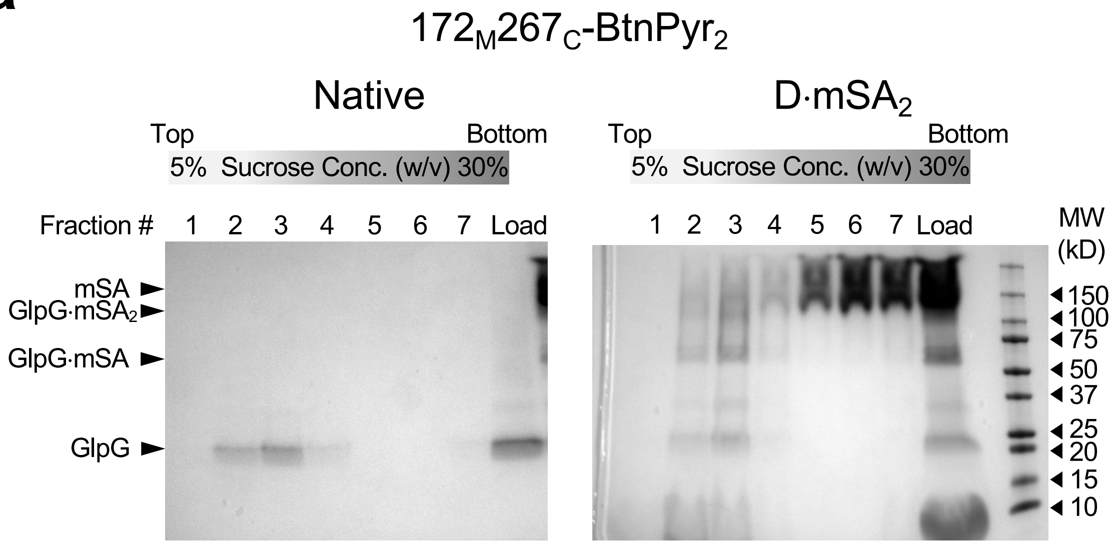


**b**

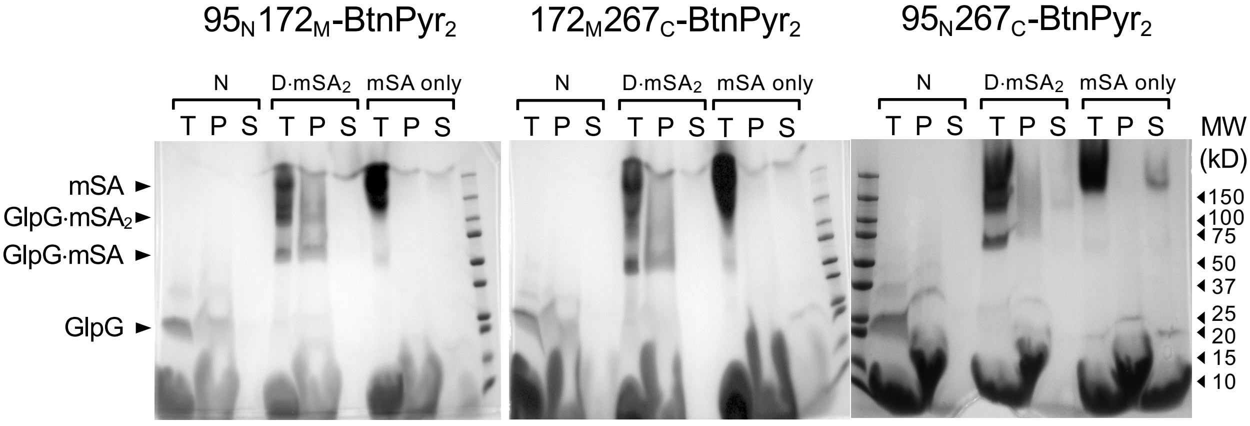


**Fig. S5. Reconstitution of native and sterically denatured GlpG in liposomes composed of *E. coli* phospholipids.**

**(a)** Liposome flotation assay for (*left*) native GlpG or (*right*) sterically denatured GlpG (D·$mSA_2$) reconstituted in liposomes. Sucrose concentration (w/v) increased from 5% (top layer, Fraction 1) to 30% (Fraction 8, bottom layer). The samples were first incubated in 30% sucrose solution and placed at the bottom. Flotation of native or denatured GlpG to the zones of lower sucrose concentration (fractions 2 and 3) indicates the association of the proteins with liposomes (see **Fig. 2b**).

**(b)** Sodium carbonate extraction of native (N) and sterically denatured GlpG (D·$mSA_2$) reconstituted in liposomes. **T:** total samples without carbonate extraction; **P:** pellet after ultracentrifugation; **S:** supernatant after ultracentrifugation. Both native and denatured GlpG samples are partitioned into the pellet, indicating transmembrane integration (i.e., not extracted).

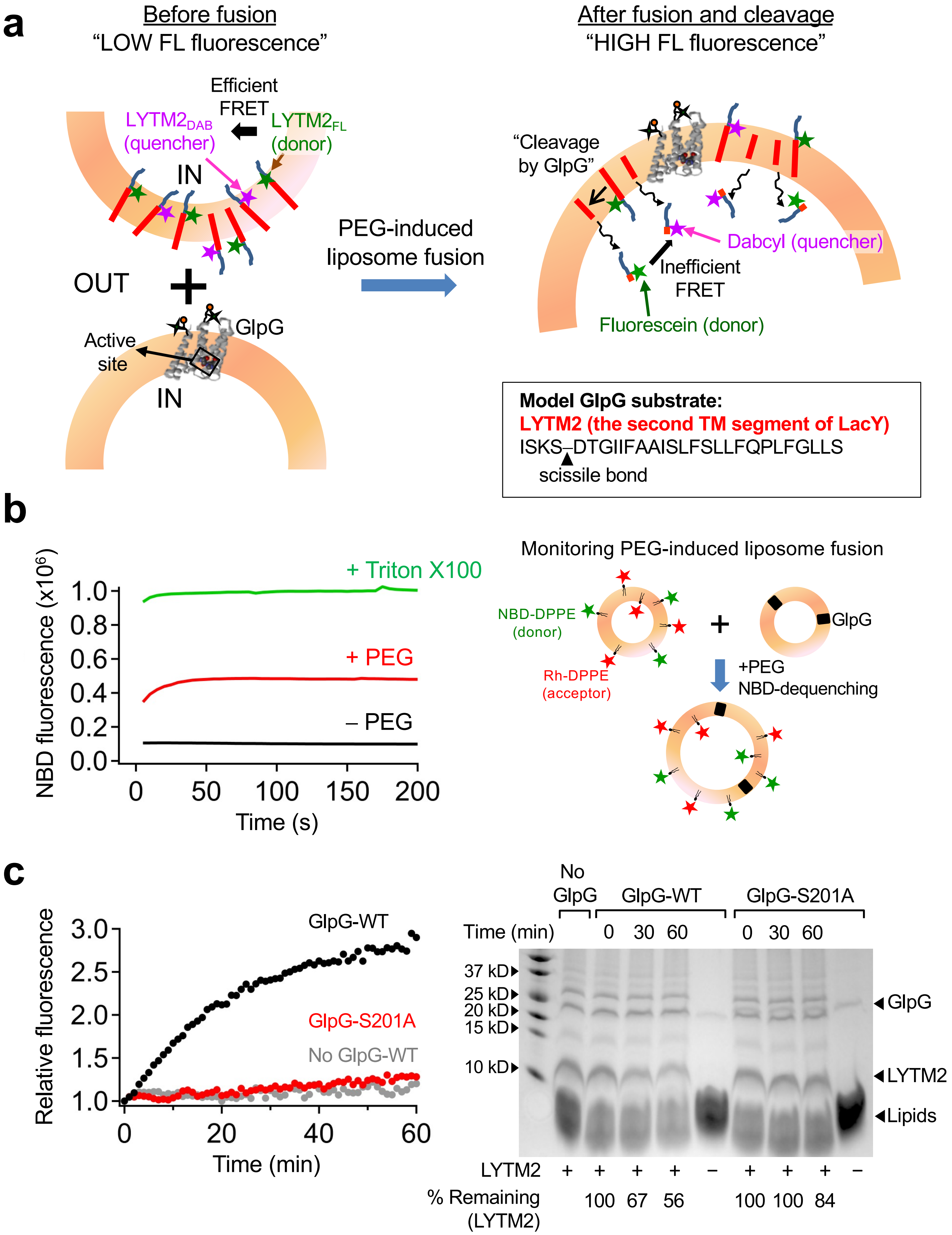

a
Before fusion
"LOW FL fluorescence"
Efficient FRET
LYTM2DAB (quencher)
LYTM2FL (donor)
IN
OUT
GlpG
Active site
IN
PEG-induced liposome fusion
After fusion and cleavage
"HIGH FL fluorescence"
"Cleavage by GlpG"
Dabcyl (quencher)
Inefficient FRET
Fluorescein (donor)
Model GlpG substrate:
LYTM2 (the second TM segment of LacY)
ISKS–DTGIIFAAISLFSLLFQPLFGLLS
scissile bond
b
NBD fluorescence (x10^6)
+ Triton X100
+ PEG
– PEG
Time (s)
Monitoring PEG-induced liposome fusion
NBD-DPPE (donor)
Rh-DPPE (acceptor)
GlpG
+PEG
NBD-dequenching
c
Relative fluorescence
GlpG-WT
GlpG-S201A
No GlpG-WT
Time (min)
No GlpG
GlpG-WT
GlpG-S201A
Time (min) 0 30 60 0 30 60
37 kD
25 kD
20 kD
15 kD
10 kD
GlpG
LYTM2
Lipids
LYTM2 + + + + – + + + –
% Remaining (LYTM2) 100 67 56 100 100 84

**Fig. S6. A fluorescence-based assay for measuring GlpG activity in *E. coli* liposomes.**

Here we developed an assay for precise measurement of GlpG activity in liposomes employing polyethylene glycol (PEG 3500)-induced liposome fusion.

**(a)** An overview of activity assay. First, two types of liposomes are prepared, one containing GlpG and the other containing a 1:1 mixture of two types of the model TM substrate LYTM2 (**Fig. S2**) labeled with the chromophores, fluorescein ($LYTM2_{FL}$, FRET donor) and nonfluorescent quencher dabcyl ($LYTM2_{DAB}$, FRET acceptor), respectively. Next, the GlpG-containing and substrate-containing liposomes are mixed upon addition of PEG to induce liposome fusion. Before fusion, fluorescein fluorescence is highly quenched due to the efficient FRET between $LYTM2_{FL}$ and $LYTM2_{DAB}$ which are confined in the same liposome. After fusion, mixing of GlpG and LYTM2 induces the cleavage of LYTM2, releasing the peptide fragments possessing chromophores into the aqueous phase, leading to lower FRET levels and an increase of fluorescein (donor) fluorescence. The rate of the increase in fluorescence intensity reflects the proteolytic activity of GlpG.

**(b)** Kinetics of PEG-induced liposome fusion which enables mixing of the enzyme GlpG and the substrate LYTM2. Fusion of the two types of proteoliposomes composed of *E. coli* phospholipids (one containing NBD (FRET-donor)- and rhodamine (FRET-acceptor)-labeled DPPE, and the other containing unlabeled wild-type GlpG) was monitored by dequenching of NBD fluorescence ($\lambda_{Ex}$ = 467 nm and $\lambda_{Em}$ = 535 nm) at 37$^{o}$C. Dead time of mixing was ~15 s. 0.5% Triton X-100 was added to induce solubilization of liposomes, which represents a positive control for complete dequenching. This result indicates that liposome fusion for enzyme-substrate mixing occurs in a faster time scale (~tens of s) than the proteolysis reaction (~tens of min, **Fig. S6c**).

**(c)** Assay results. (*Left*) Upon addition of PEG that induces liposome fusion, fluorescein (FL) fluorescence was monitored over time. In the presence of WT GlpG, FL fluorescence increases while Inactivated mutant GlpG S201A induces no change in fluorescence, as does the addition of empty vesicles without WT GlpG. (*Right*) Time-dependent proteolysis of LYTM2 in liposomes monitored by SDS-PAGE. The band intensity of LYTM2 disappears in the presence of GlpG WT, but not in the presence of GlpG S201A. Therefore, dequenching of FL fluorescence is indicative of the proteolysis of LYTM2 by GlpG. "%-remaining" denotes the band intensity of full length LYTM2 at each time point relative to that at time zero. The band intensities were quantified using the ImageJ program.

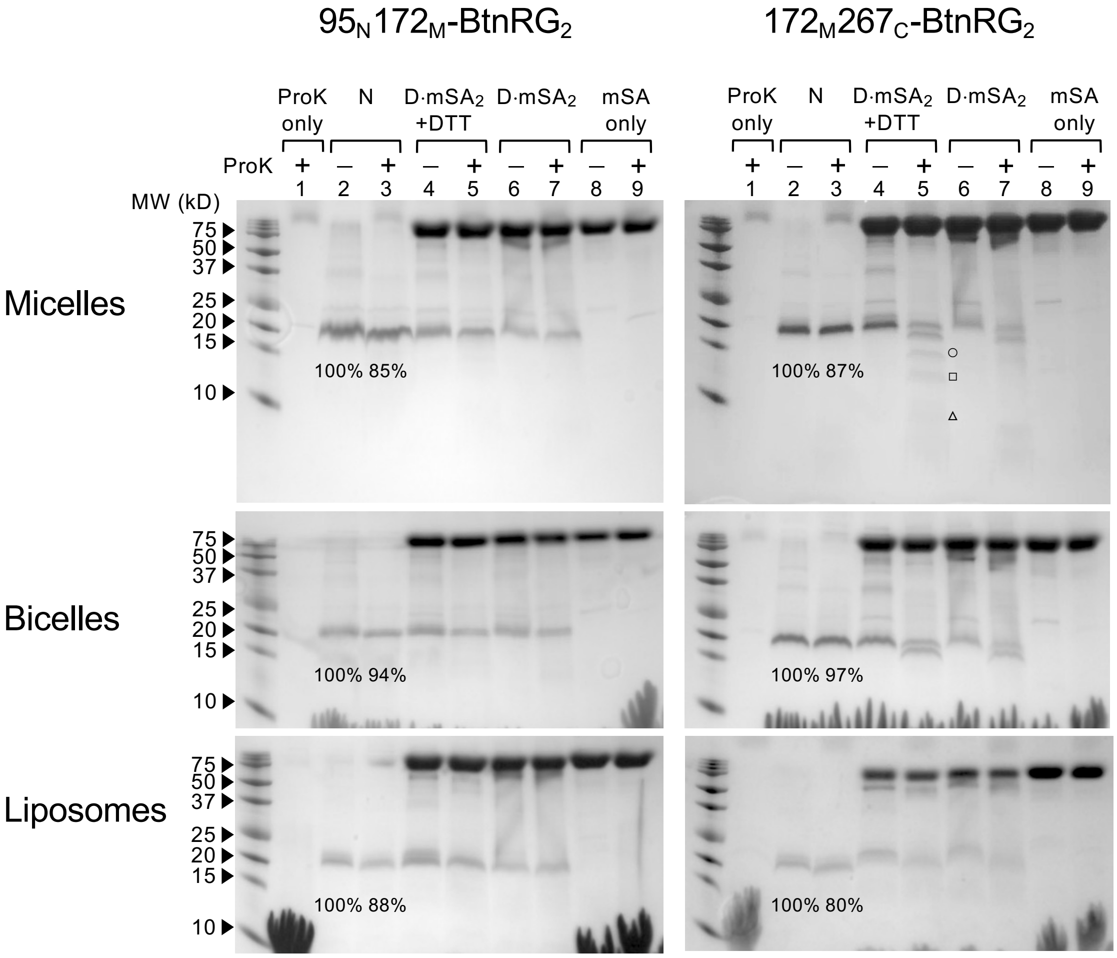


**Fig. S7. Proteolysis of native and sterically denatured GlpG by Proteinase K.**

Native (N) and sterically denatured (D·$mSA_2$) GlpG were prepared in (*top*) DDM micelles, (*middle*) DMPC:DMPG:CHAPS bicelles, and (*bottom*) *E. coli* liposomes. Native GlpG (lanes 2 and 3), GlpG denatured with mSA (lanes 4–7), and mSA only (lanes 8 and 9) were incubated without or with Proteinase K (ProK) for 30 minutes. Denatured GlpG incubated with or without Proteinase K were further treated with DTT (lanes 4 and 5) to break the disulfide bond between GlpG and BtnRG label bound with mSA. Release of bound mSA upon addition of DTT enables direct comparison of the band intensities of GlpG to quantify the fraction digested by ProK. The proteolytic reaction was terminated using PMSF.

89 ± 5% of native $95_N172_M$-$BtnRG_2$ and 88 ± 9% of native $172_M267_C$-$BtnRG_2$ were resistant to proteolysis (lanes 2 and 3) while 52 ± 5% of denatured $95_N172_M$-$BtnRG_2$ and 37 ± 9% of denatured $172_M267_C$-$BtnRG_2$ were resistant to proteolysis, indicating that denatured GlpG was more susceptible to proteolysis. The band intensities were quantified using the ImageJ program.

The proteolytic peptide fragment larger than 10 kDa is marked with a symbol (open circles, 17 kDa; open squares, 13 kDa; open triangles, 11 kDa) on the right side of each band. These larger proteolytic products probably contain the N-subdomain since subglobal denaturation in the C-subdomain induces proteolysis of the C-subdomain and protection of the N-subdomain.

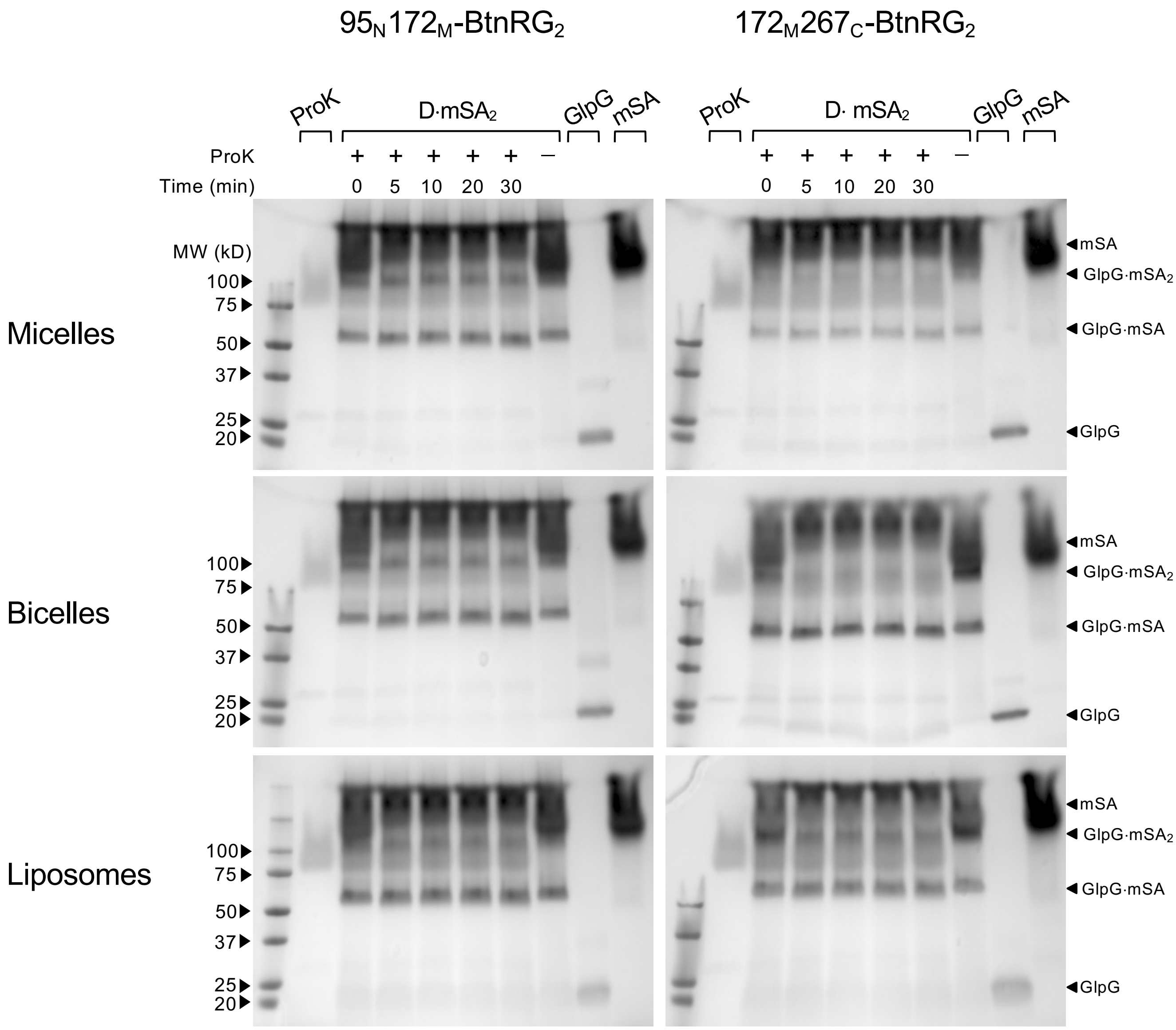


**Fig. S8. Susceptibility of unlabeled (GlpG), single mSA-bound (GlpG·mSA) and double mSA-bound (*i.e.*, sterically denatured, D·mSA$_2$ or GlpG·mSA$_2$) GlpG to proteolysis by Proteinase K.**

Time-dependent proteolysis of GlpG in various mSA-bound states in (*top*) DDM micelles, (*middle*) DMPC/DMPG/CHAPS bicelles, and (*bottom*) *E. coli* liposomes. Denatured GlpG bound with two mSA molecules (D·mSA$_2$ or GlpG·mSA$_2$) is preferentially degraded by ProK, whereas GlpG unbound (GlpG) or bound with one mSA molecule (GlpG·mSA) is relatively resistant to proteolysis.

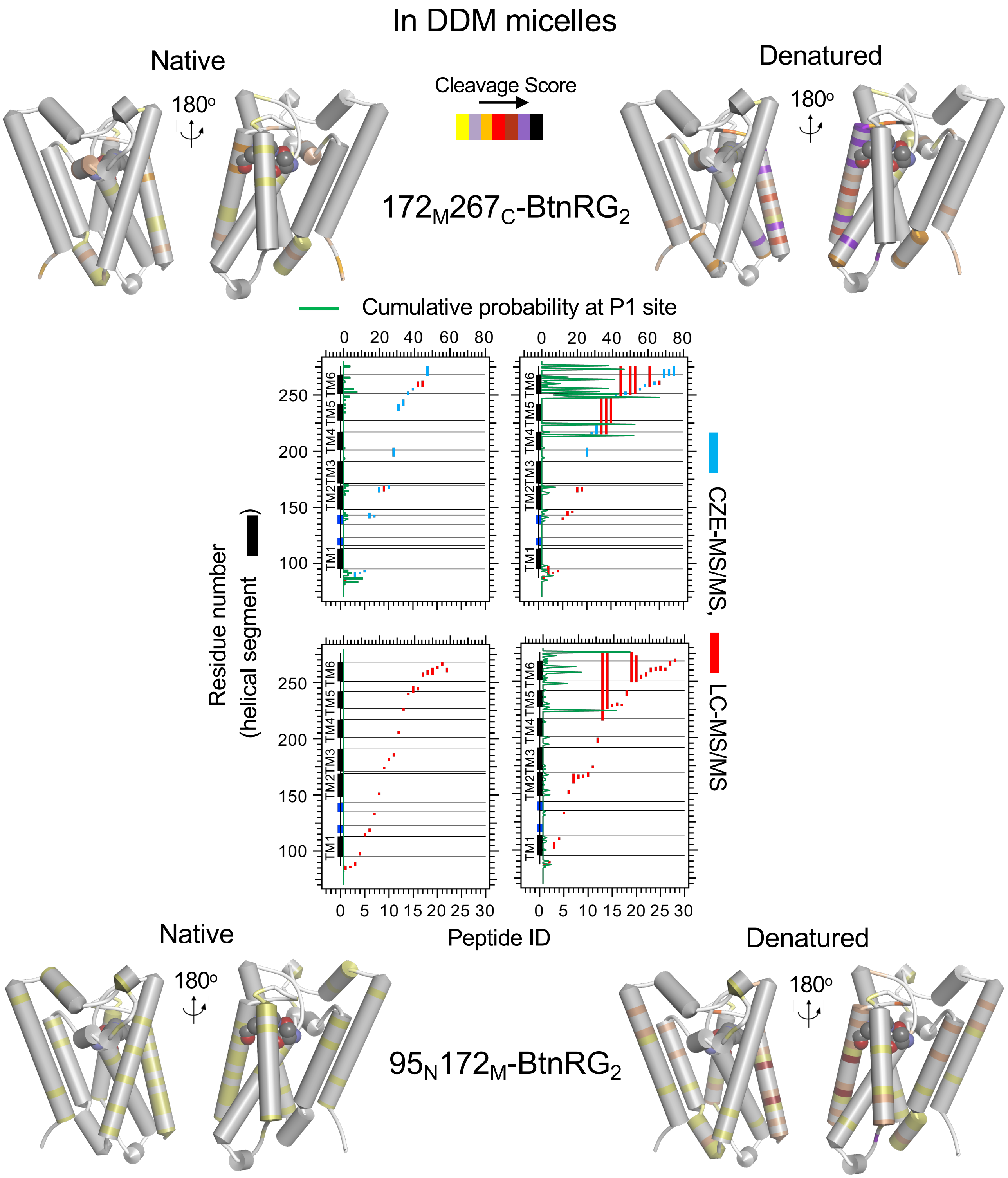


**Fig. S9 Mapping the flexible structural regions in folded and sterically denatured GlpG using limited proteolysis by Proteinase K and mass spectrometry.** Folded and denatured GlpG variants, $172_M267_C$-BtnRG$_2$ (*top*) and $172_M267_C$-BtnRG$_2$ (*bottom*) in DDM micelles were proteolyzed using Proteinase K and the proteolysis products were analyzed using CZE-MS/MS or LC-MS/MS. Analyzed peptides and the confidence scores (log [Probability]) summed at each P1 cleavage site were mapped on to the sequence and secondary structural elements (*center*) and the tertiary structure (*left and right*). The catalytic dyad (Ser201 and His254) are shown in spheres.

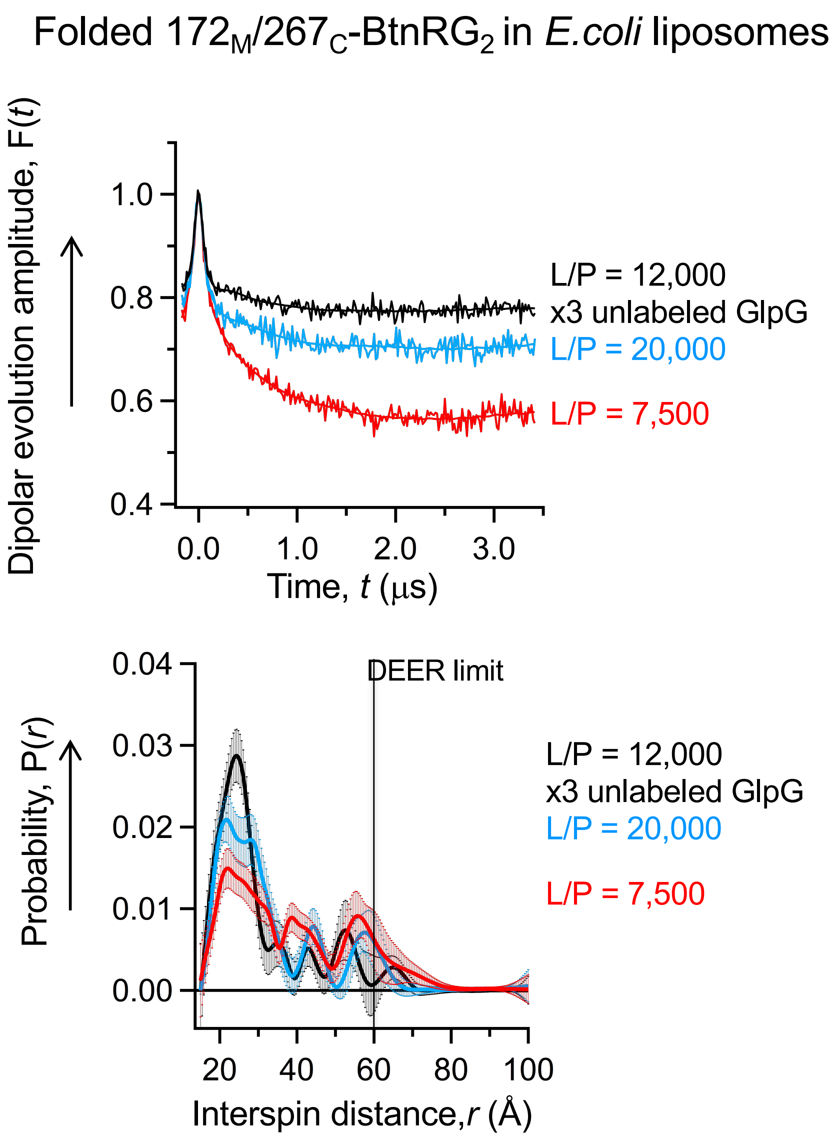


**Fig. S10. Optimization of the sample condition for measuring the intramolecular spin-spin distances in native GlpG in *E. coli* liposomes.**

In the case of $172_M267_C$–$BtnRG_2$ reconstituted in liposomes at the protein-to-lipid molar ratio (L/P) of 7,500, a large portion (~50%) of the long-distance components (~40 Å and ~60 Å) that are absent in micelles and bicelles emerged (**Fig. 4b**). We suspected that these long-distance components may have stemmed from the intermolecular dipolar coupling caused by multiple incorporation of spin-labeled GlpG molecules in a single liposome. To suppress this unwanted contribution, we increased the L/P up to 20,000 or co-reconstituted a molar excess of unlabeled folded GlpG (the inactive variant S201A) with spin-labeled GlpG at L/P = 12,000.

Both attempts substantially reduced the probability of the longer distance components and the modulation depth. The modulation depth is related to the concentration of spin pairs whose distances are within the DEER distance limit (55–60 Å). Co-reconstitution of an excess unlabeled GlpG was more effective than increasing the L/P to suppress the intermolecular dipolar coupling.

Each raw dipolar evolution trace (*top*) was fitted to the non-negative Tikhonov regularization algorithm. The error bar at each distance corresponds to the ± s.d. of fitting. This uncertainty in fitting originates from various factors including simultaneous fitting of multiple distance components, minute perturbation of F($t$) at $t$ = 0, data noise, imperfect background subtraction, *etc*.

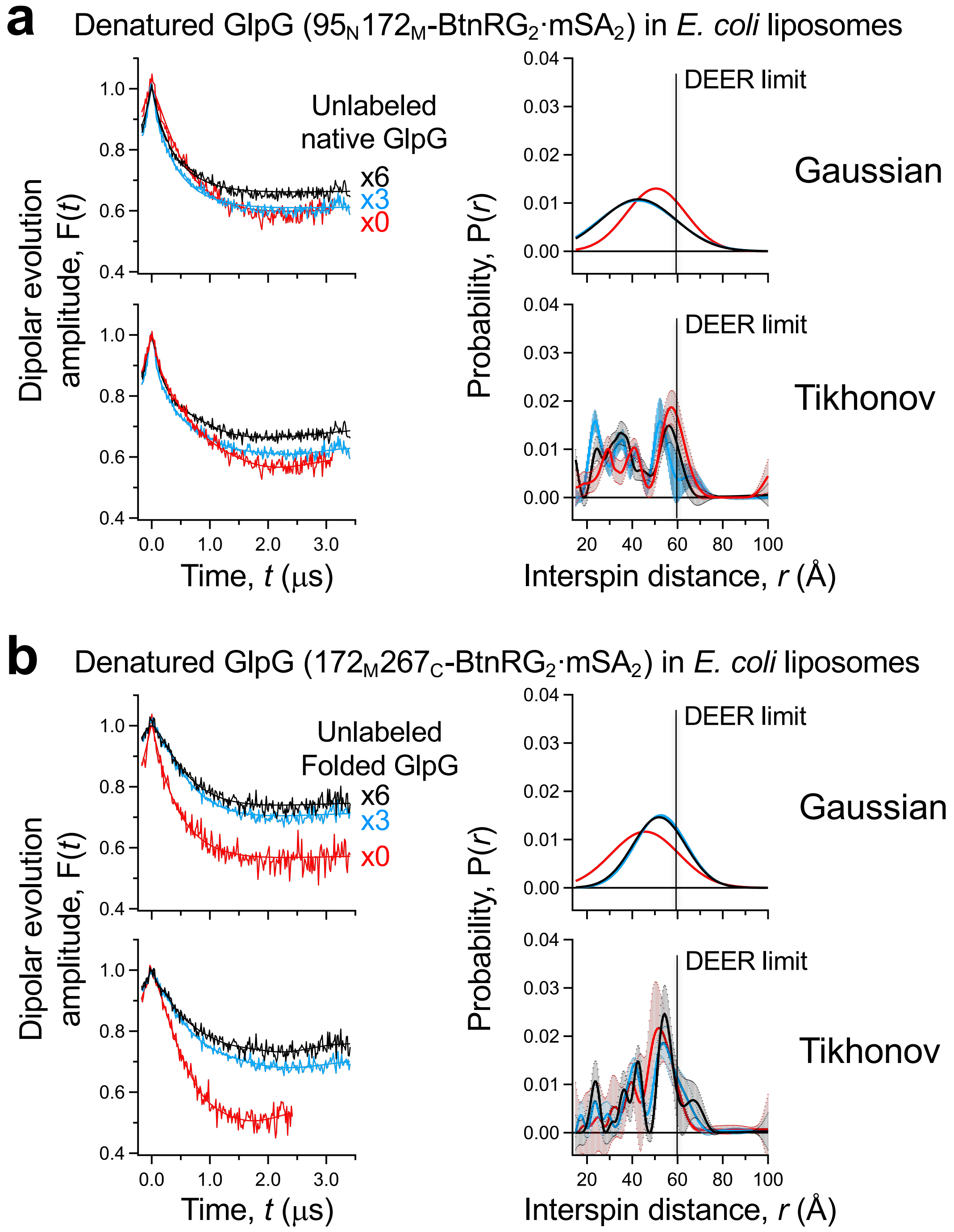


**Fig. S11. Optimization of the sample condition for measuring the intramolecular spin-spin distances in denatured GlpG in *E. coli* liposomes.**

Double-biotin variants used: **a.** $95_N172_M$-$BtnRG_2$**; b.** $172_M267_C$-$BtnRG_2$

To optimize the condition for measuring the intramolecular spin-spin distances in the denatured states in liposomes, we incorporated unlabeled GlpG (S201A) at an increasing molar excess (x0, x3 and x6, relative to spin-labeled GlpG) at L/P = 20,000. While the modulation depth successively decreased with an increasing amount of unlabeled GlpG, the interspin distance distributions did not noticeably change.

Each background-subtracted dipolar evolution trace (*left* panels) was fitted to the single Gaussian model or the non-negative Tikhonov regularization algorithm. The error bar at each distance obtained from non-negative Tikhonov regularization (*lower right* panels in **a** and **b**) corresponds to the ± s. d. of fitting.

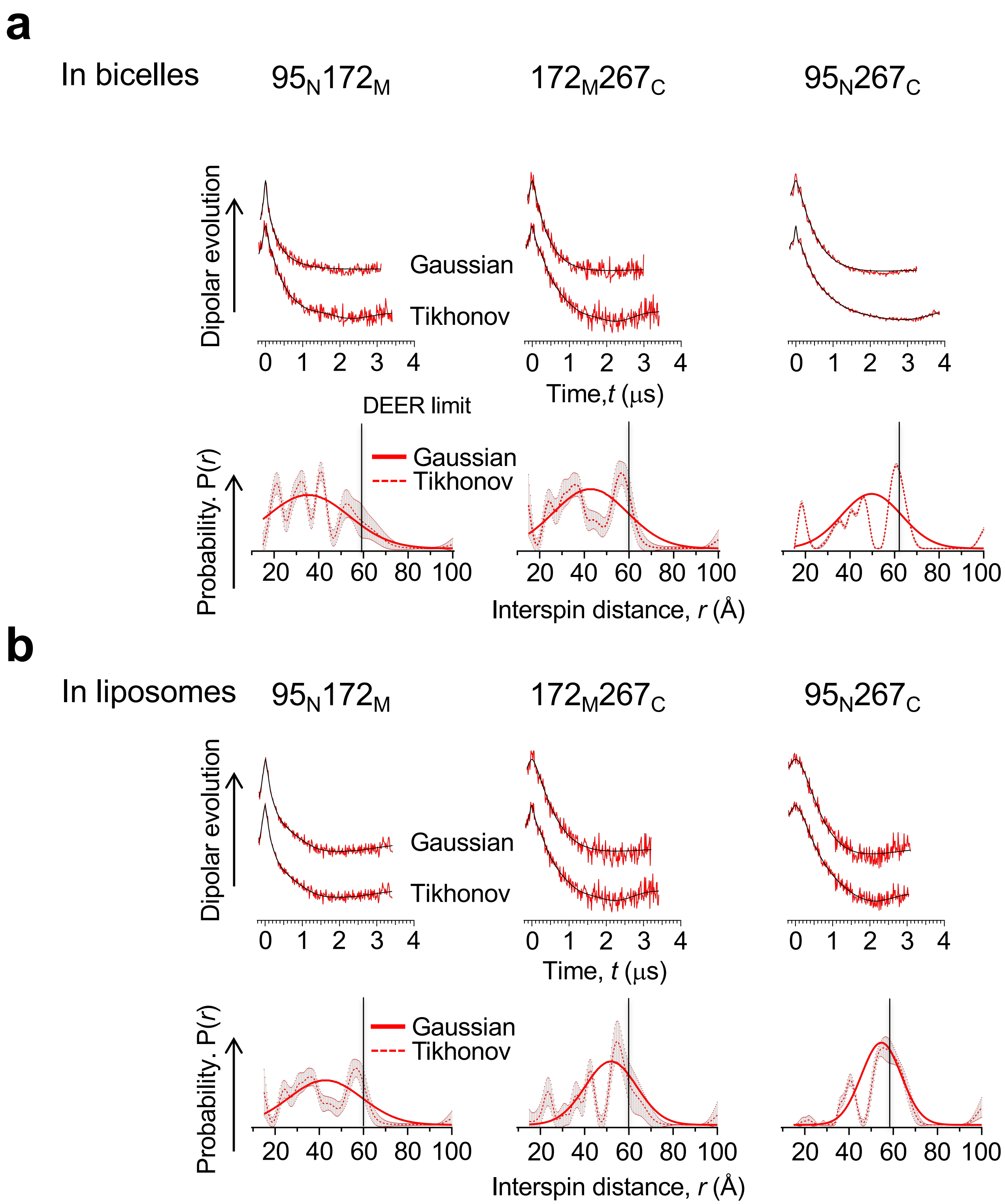


**Fig. S12. Comparison between the Guassian model and non-negative Tikhonov regularization algorithm for distance fitting of the dipolar evolution data.**

The DEER data for the denatured states of three double biotin variants bound with mSA in **(a)** bicelles and **(b)** liposomes.

The error bar at each distance obtained from non-negative Tikhonov regularization corresponds to ± s. d. from fitting.

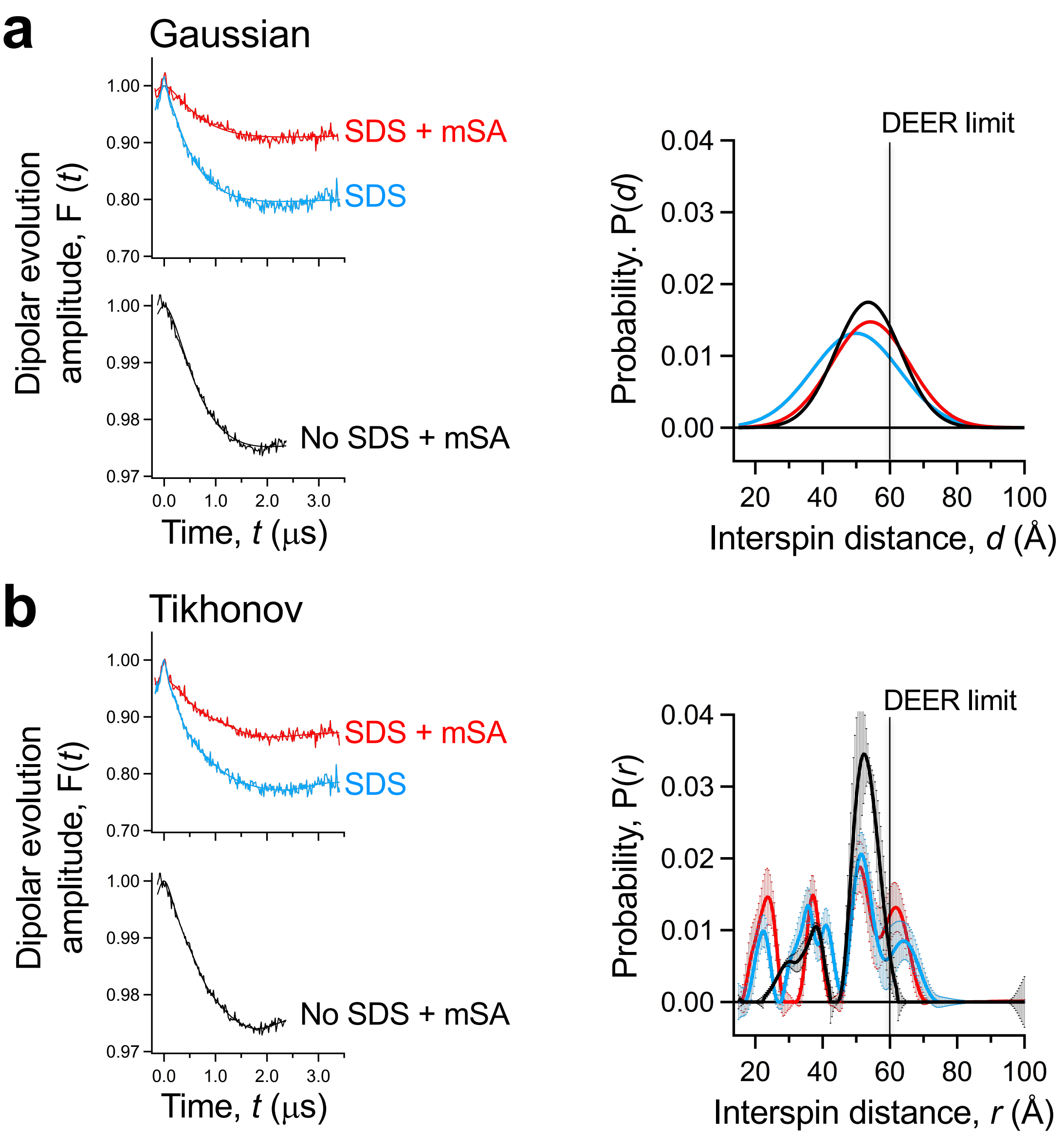


**Fig. S13. The effect of doubly bound mSA molecules on the compactness of the denatured state ensemble (DSE).**

The DEER data obtained for the double biotin variant of GlpG $172_M267_C$–$BtnRG_2$ were analyzed using the single Gaussian model (**a**) or non-negative Tikhonov (**b**) algorithm. The error bar at each distance in (**b**) corresponds to ± s. d. of fitting. The final SDS concentration was ~150 mM (~4% w/v).

To test if doubly bound mSA molecules affected the degree of expansion of the DSE, the interspin distances were measured for the SDS-induced DSE in the absence and presence of bound mSA molecules. Once GlpG is denatured by SDS, bound mSA induced slight expansion of the DSE (red vs cyan traces) indicating that bound mSA induced additional expansion of the DSE.

Another potential concern is that attractive or repulsive intermolecular interactions between bound mSA molecules may have affected the degree of expansion of the DSE. The interspin distances with bound mSA in SDS were similar to those with mSA in nondenaturing DDM micelles (red vs black traces). Addition of negatively charged SDS molecules would bind to mSA and SDS-bound mSA molecules would exert repulsive forces with each other. The finding that the interspin distances remain unchanged in the presence and absence of SDS indicates that repulsive or attractive interactions between bound mSA molecules, even if they exist, do not noticeably affect the compactness of the DSE.

**a**

At $T_{av}$ = 274 K (271 → 277 K in 0.34 K steps)

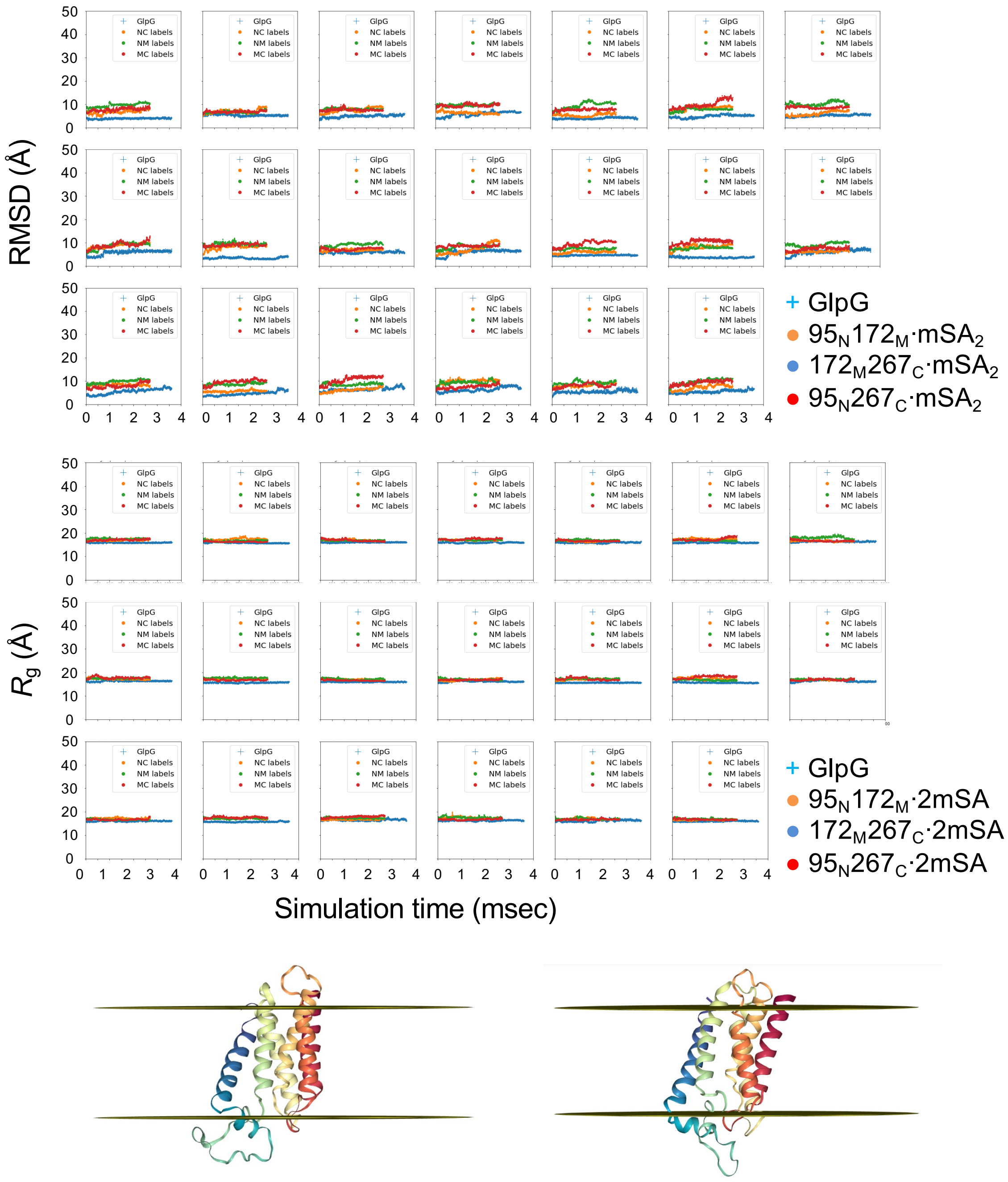


**Fig. S14. *Upside* simulations of GlpG alone and the three constructs doubly bound with mSA.** Time-dependent fluctuations in $C_\alpha$-RMSD (*top*) and $R_g$ (*middle*) are referenced to native GlpG (the mSA molecules are not included in calculation) under the constraints of the depth-dependent statistical membrane burial potential. (*Bottom*) Representative structural snapshots during simulation. The 20 independent trajectories were run over a narrow temperature range 271-277 K in 0.34 K steps. Data shown do not include a 0.4 msec equilibration period.

**a)** At $T_{av}$ = 274K, the RMSD and $R_g$ are stable over the simulations, and the protein remains close to the native structure.

**b** At $T_{av}$ = 308 K (304 → 312 K in 0.34 K steps)

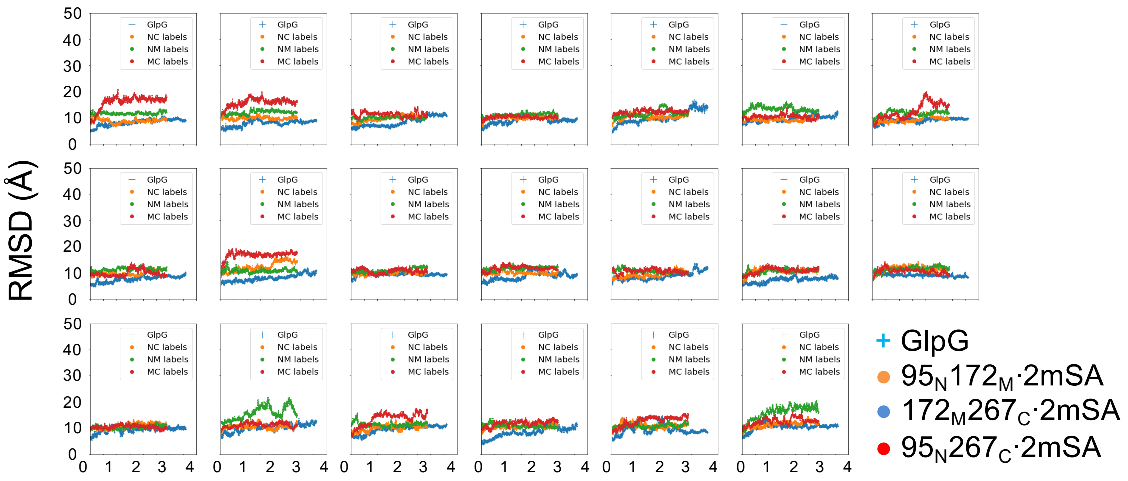


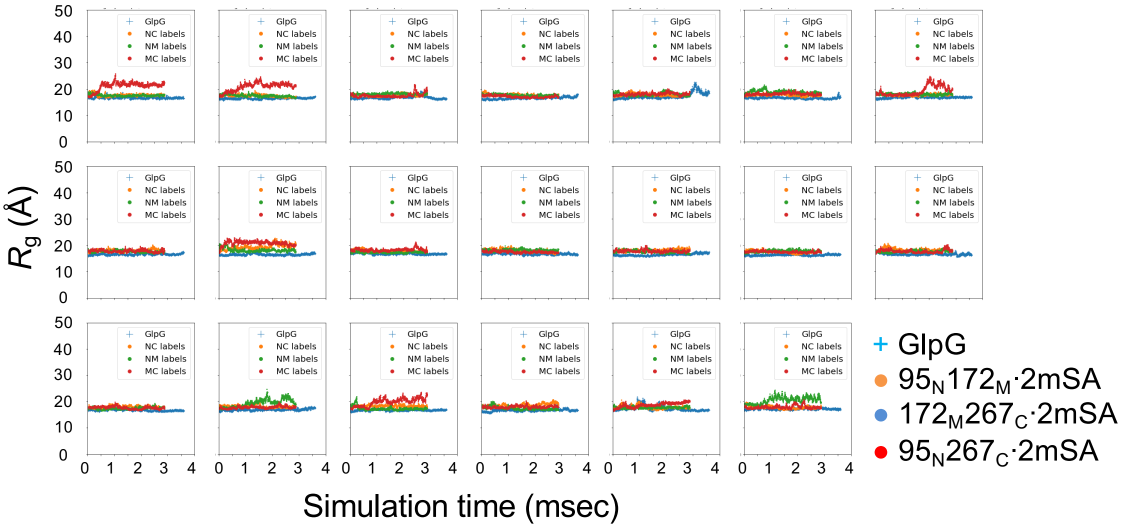


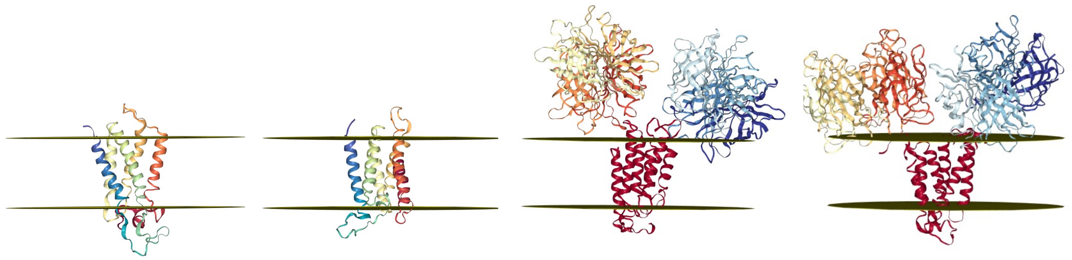

**Continued-Fig. S14. *Upside* simulations of the DSEs of GlpG alone and the three constructs doubly bound with mSA .**

**b:** At $T_{av}$ = 308 K, the conformations are still compact while the RMSDs and $R_g$s start to deviate from the native state. Thus, this state is defined as a “collapsed denatured state”

**c** At $T_{av}$ = 343 K (340 → 346 K in 0.34 K steps)

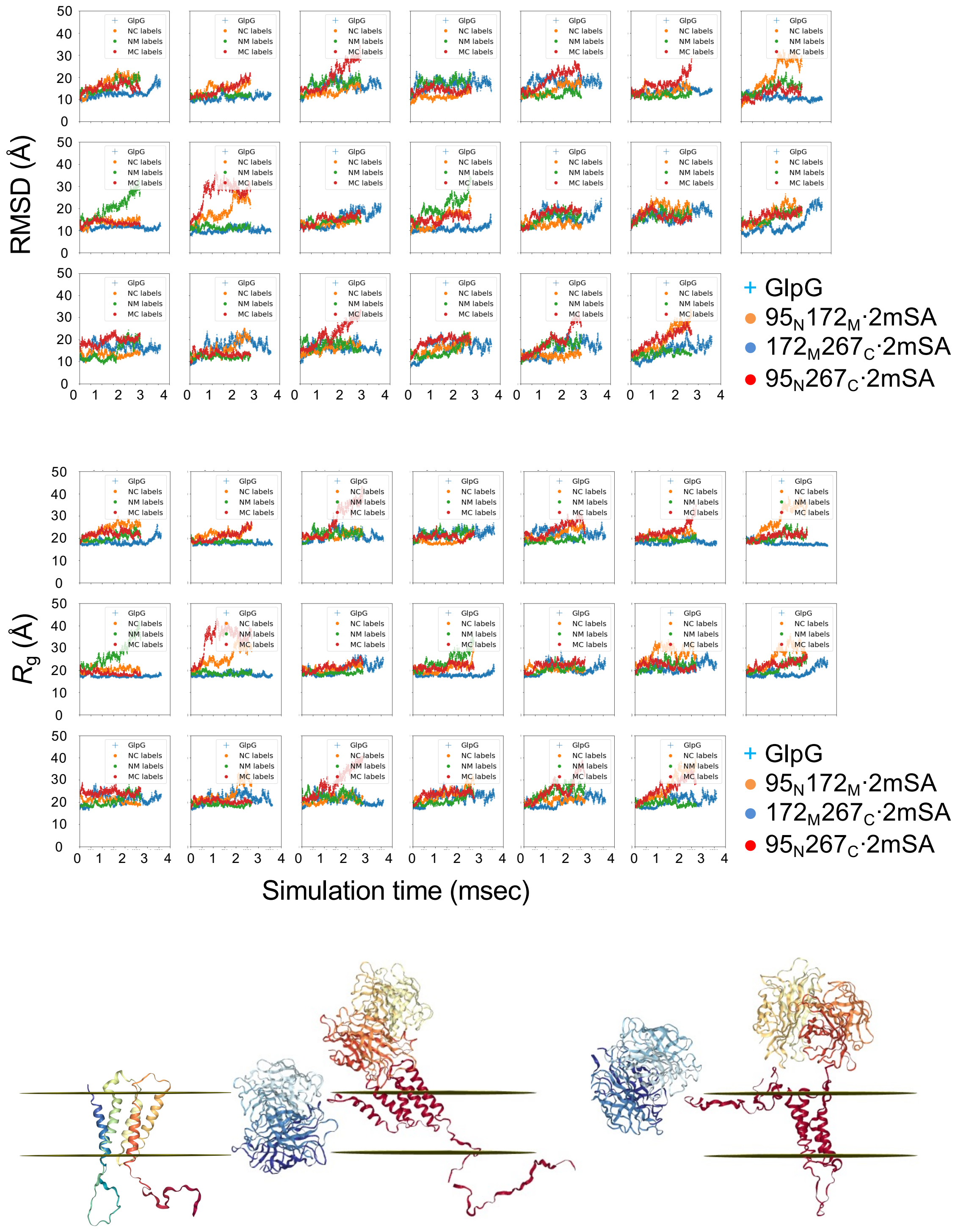


**Continued-Fig. S14. *Upside* simulations of the DSEs of GlpG alone and the three constructs doubly bound with mSA .**

**c:** At $T_{av}$ = 343 K, GlpG fluctuates with a large occasional increase in RMSD and a moderate increase in $R_g$. TM helical structures are largely retained with transient unfolding at the membrane surface.

**d**

At $T_{av}$ = 377 K (373 → 380 K in 0.34 K steps)

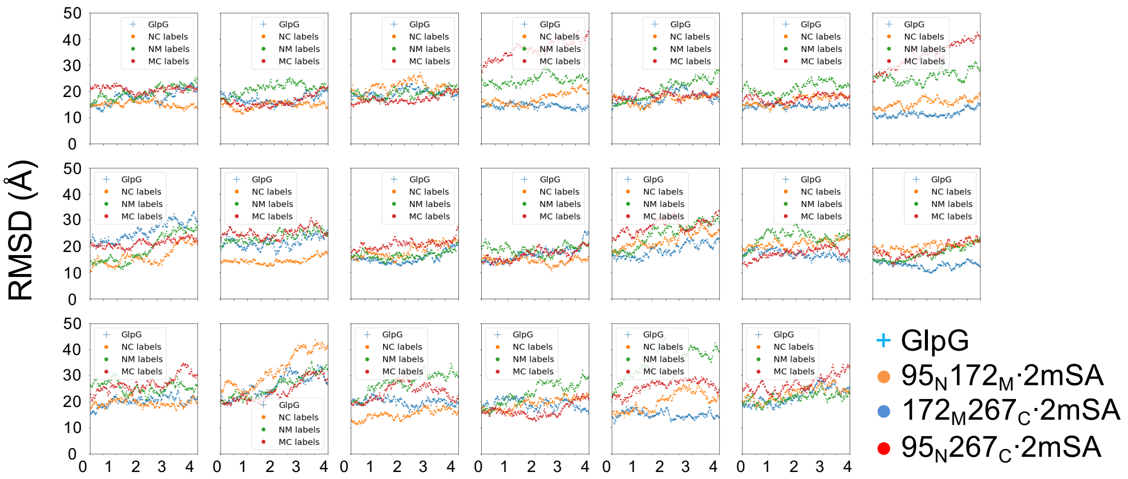


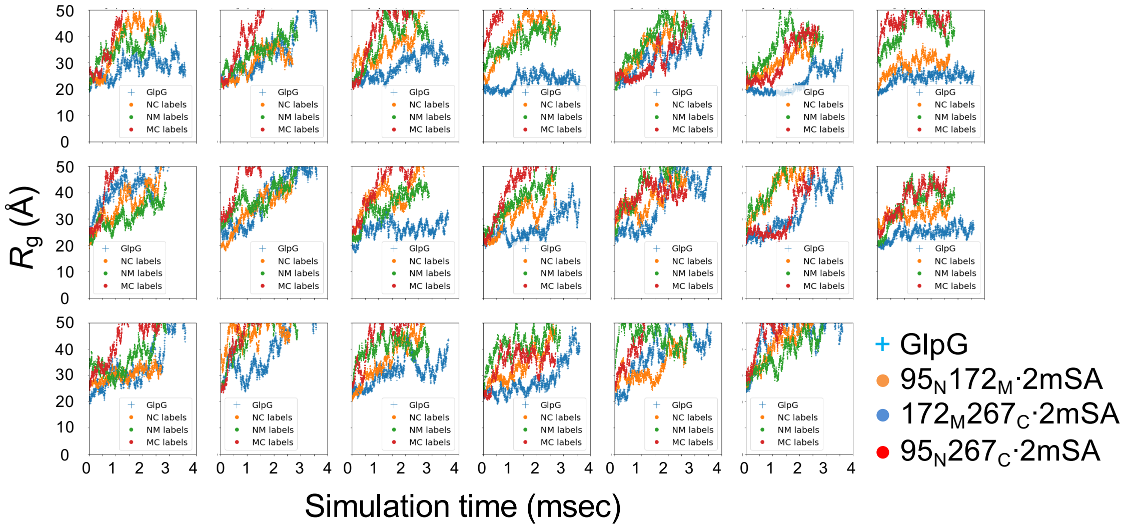


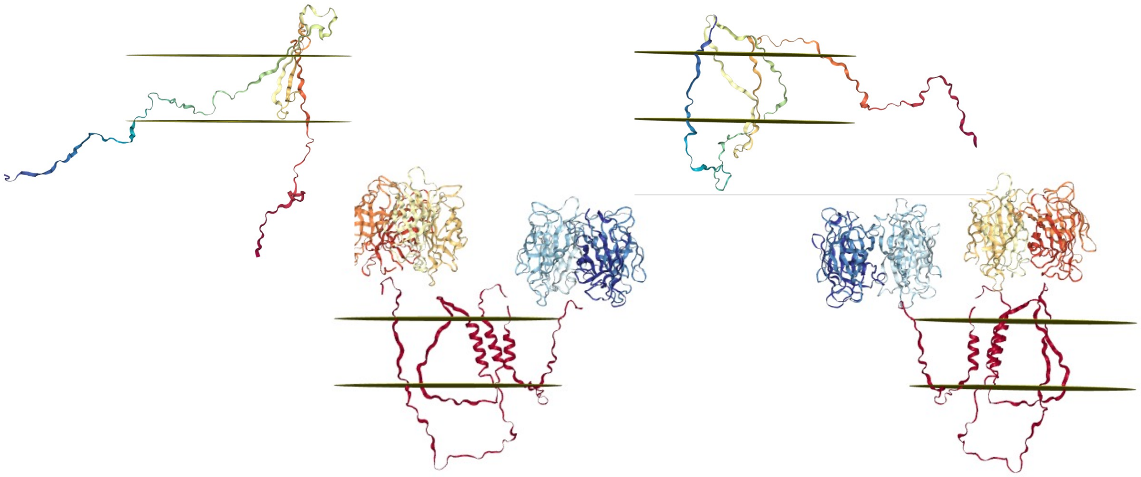

**Continued-Fig. S14. *Upside* simulations of the DSEs of GlpG alone and the three constructs doubly bound with mSA .**

**d:** At $T_{av}$ = 377 K, GlpG substantially unfolds losing the TM helical structures inside and outside of the membrane.

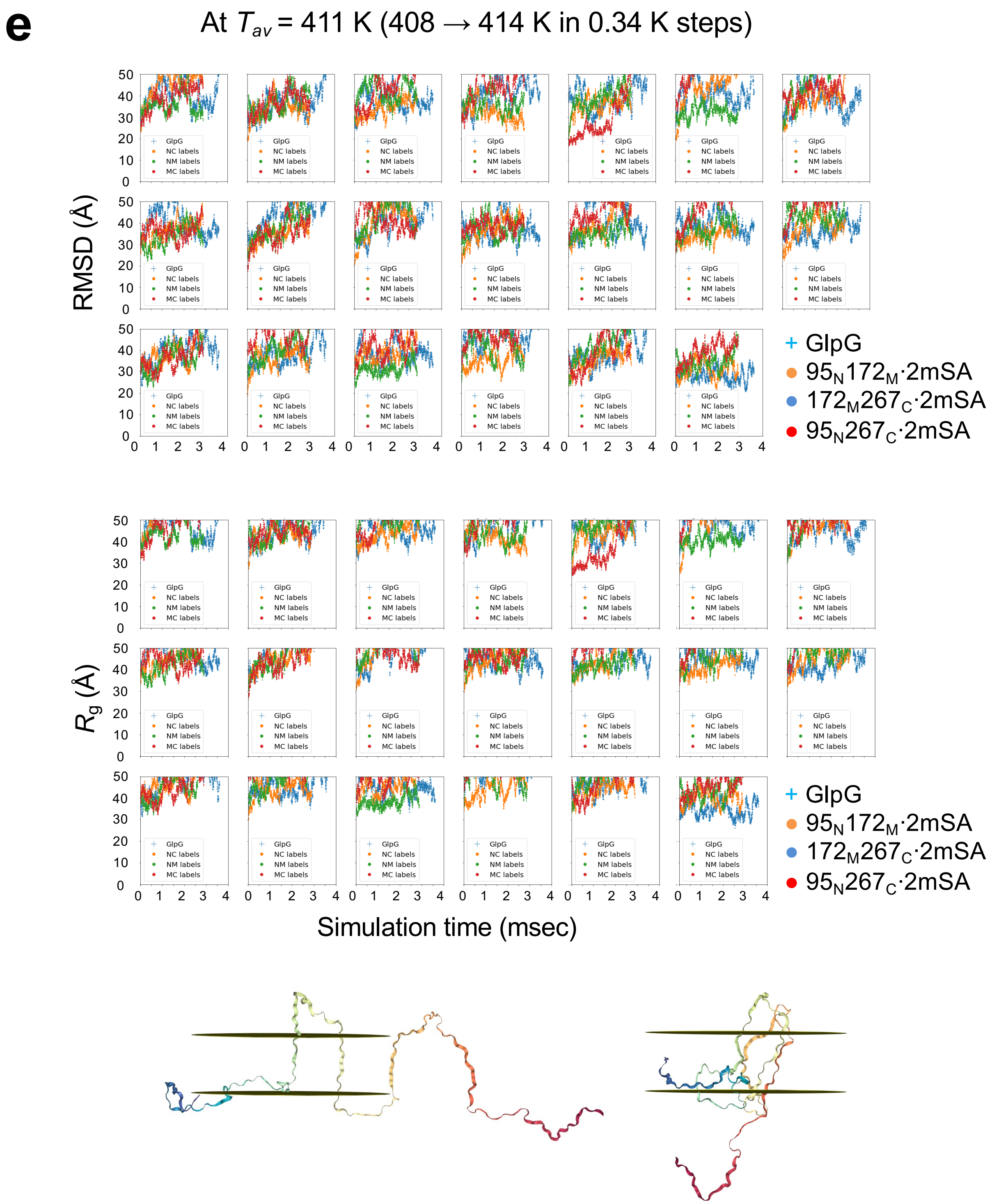


**Continued-Fig. S14.** ***Upside*** **simulations of the DSEs of GlpG alone and the three constructs doubly bound with mSA .**

**e:** At $T_{av}$ = 411 K, the TM segments almost completely unfold interconverting between transmembrane and water-exposed topologies.

**f** At $T_{av}$= 240 K (236 → 244 K in 0.34 K steps)

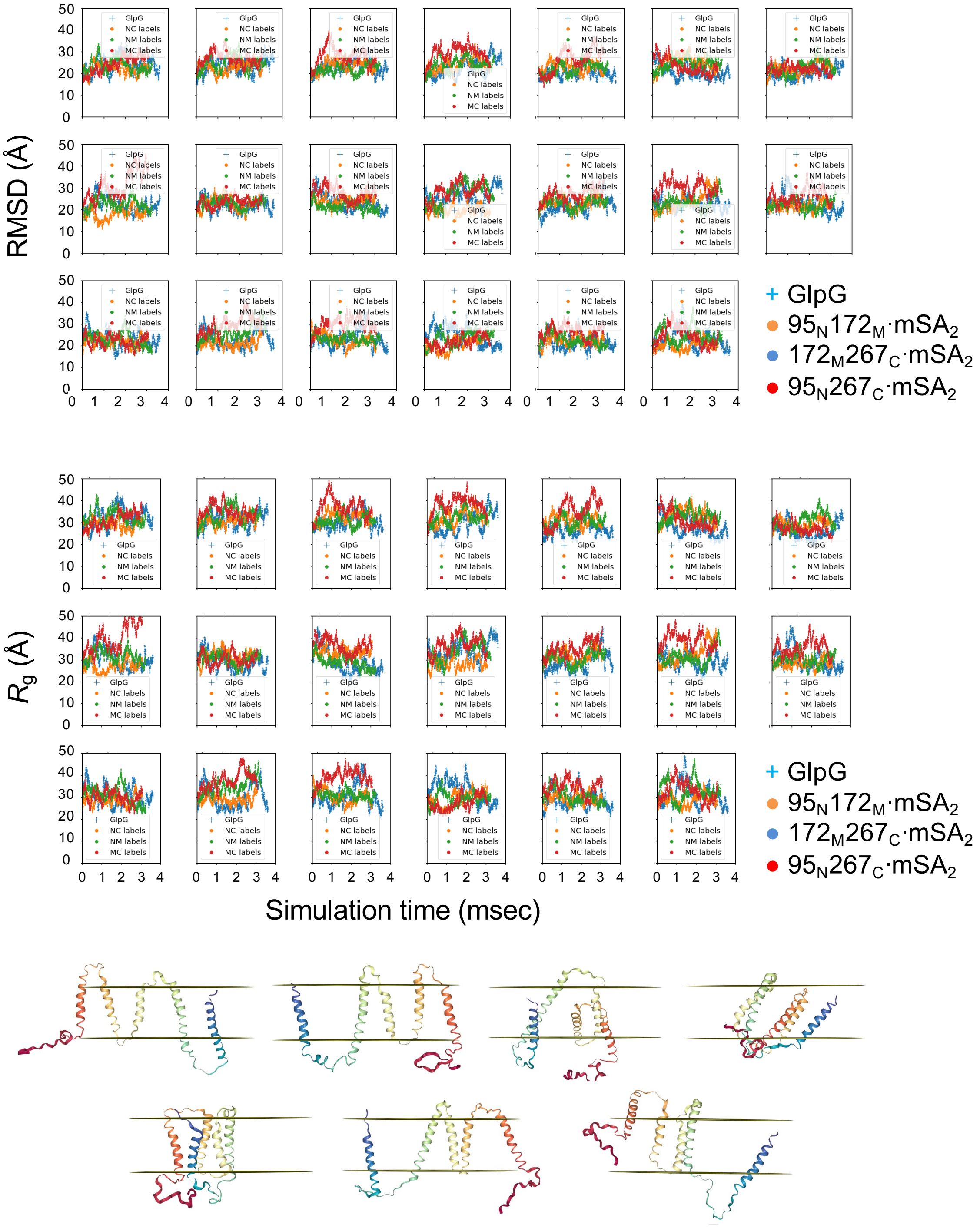


**Continued-Fig. S14.** ***Upside*** **simulations of the DSEs of GlpG alone and the three constructs doubly bound with mSA .**

**f:** Simulation at $T_{av}$ = 240 K. Here, the backbone hydrogen bonds in each TM segment are allowed to form while the side chain attractive terms are turned off. The simulation was performed at a low temperature to maintain the TM helices to a maximal extent. Thus, this state is defined as a “expanded denatured state”.

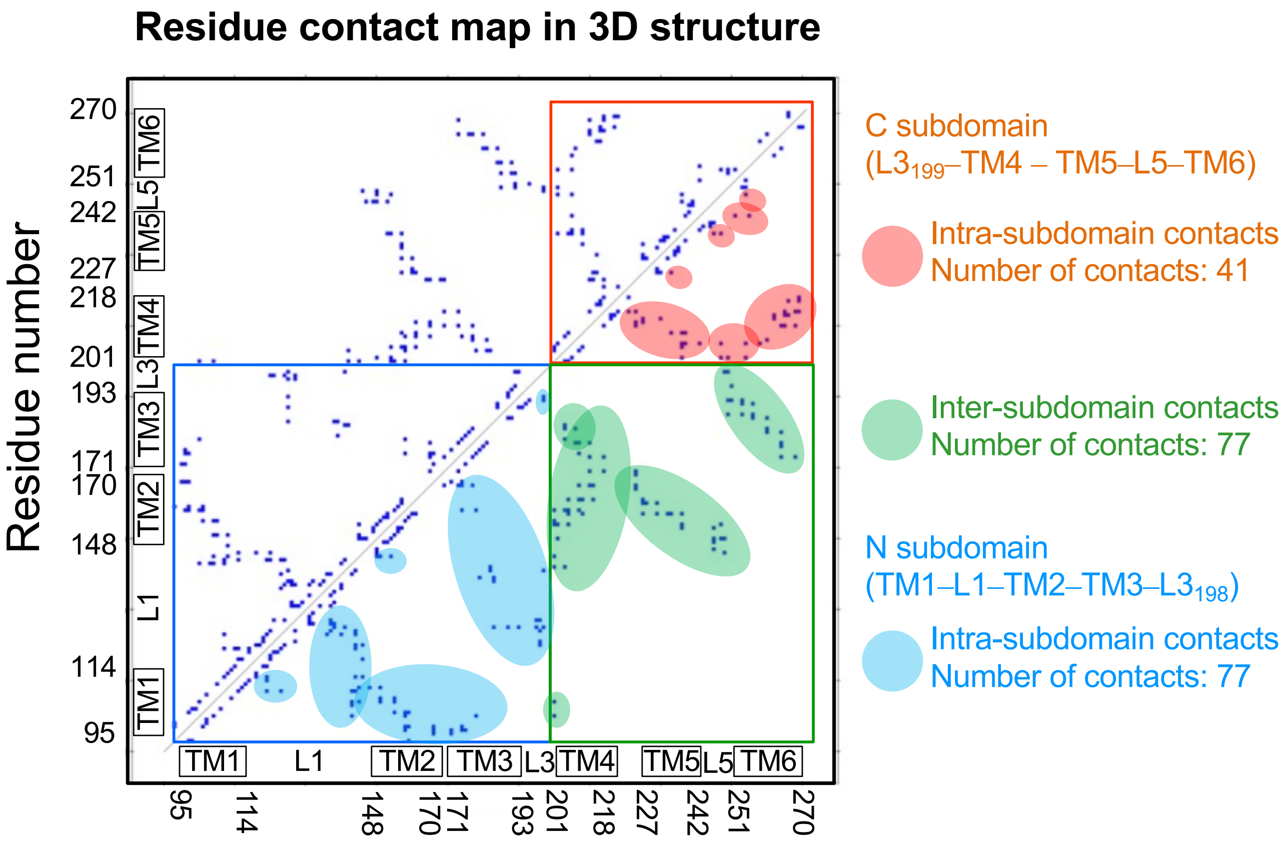


**Fig. S15. Analysis of residue contacts in the structure of GlpG (PDB: 3B45).**

The residue pairs whose $C_\alpha$–$C_\alpha$ distances are within 6 Å are mapped (Discovery Studio program, Accelrys). N-subdomain spans TM1–L1–TM2–TM3–$L3_{198}$ while C-subdomain covers $L3_{199}$–TM4 – TM5–L5–TM6.
